# Loss of an extensive ciliary connectome induces proteostasis and cell fate switching in a severe motile ciliopathy

**DOI:** 10.1101/2024.03.20.585965

**Authors:** Steven L. Brody, Jiehong Pan, Tao Huang, Jian Xu, Huihui Xu, Jeffrey Koenitizer, Steven K. Brennan, Rashmi Nanjundappa, Thomas G. Saba, Andrew Berical, Finn J. Hawkins, Xiangli Wang, Rui Zhang, Moe R. Mahjoub, Amjad Horani, Susan K. Dutcher

**Author notes:** Correspondence: Co-corresponding, S. L. Brody. Co-corresponding, S. K. Dutcher.

## Abstract

Motile cilia have essential cellular functions in development, reproduction, and homeostasis. Genetic causes for motile ciliopathies have been identified, but the consequences on cellular functions beyond impaired motility remain unknown. Variants in *CCDC39* and *CCDC40* cause severe disease not explained by loss of motility. Using human cells with pathological variants in these genes, *Chlamydomonas* genetics, cryo-electron microscopy, single cell RNA transcriptomics, and proteomics, we identified perturbations in multiple cilia-independent pathways. Absence of the axonemal CCDC39/CCDC40 heterodimer results in loss of a connectome of over 90 proteins. The undocked connectome activates cell quality control pathways, switches multiciliated cell fate, impairs microtubule architecture, and creates a defective periciliary barrier. Both cilia-dependent and independent defects are likely responsible for the disease severity. Our findings provide a foundation for reconsidering the broad cellular impact of pathologic variants in ciliopathies and suggest new directions for therapies.

## Introduction

Fluid propelled by motile cilia is an essential function during development at the embryonic node, and flow in the brain, reproductive, and respiratory tracts^1–3^. These tasks are accomplished by molecular motors with companion regulatory complexes docked along the length of the cilia’s microtubule skeleton, the axoneme^4^. Seminal studies of cilia from *Chlamydomonas reinhardtii* using freeze-fracture with transmission electronic microscopy (EM) uncovered a 96-nm repeating pattern of motors and other structures on the outer circumference of the nine doublet microtubules (DMT)^5,6^ (**Figure 1A**). Recent single particle cryo-EM of cilia provides exquisite detail that allows the identification of proteins and their position along the ciliary axoneme^7–14,15^. In parallel, advances in proteomics and genomics provide a catalog of motile ciliary components and pathogenic gene variants ^16–21^. These complementary discovery tools moved the understanding of cilia biology and human disease forward. However, a major gap exists in understanding how the ciliary proteins are placed in a precise, repeating pattern and how those processes are disrupted in disease variants.

**Figure 1.**
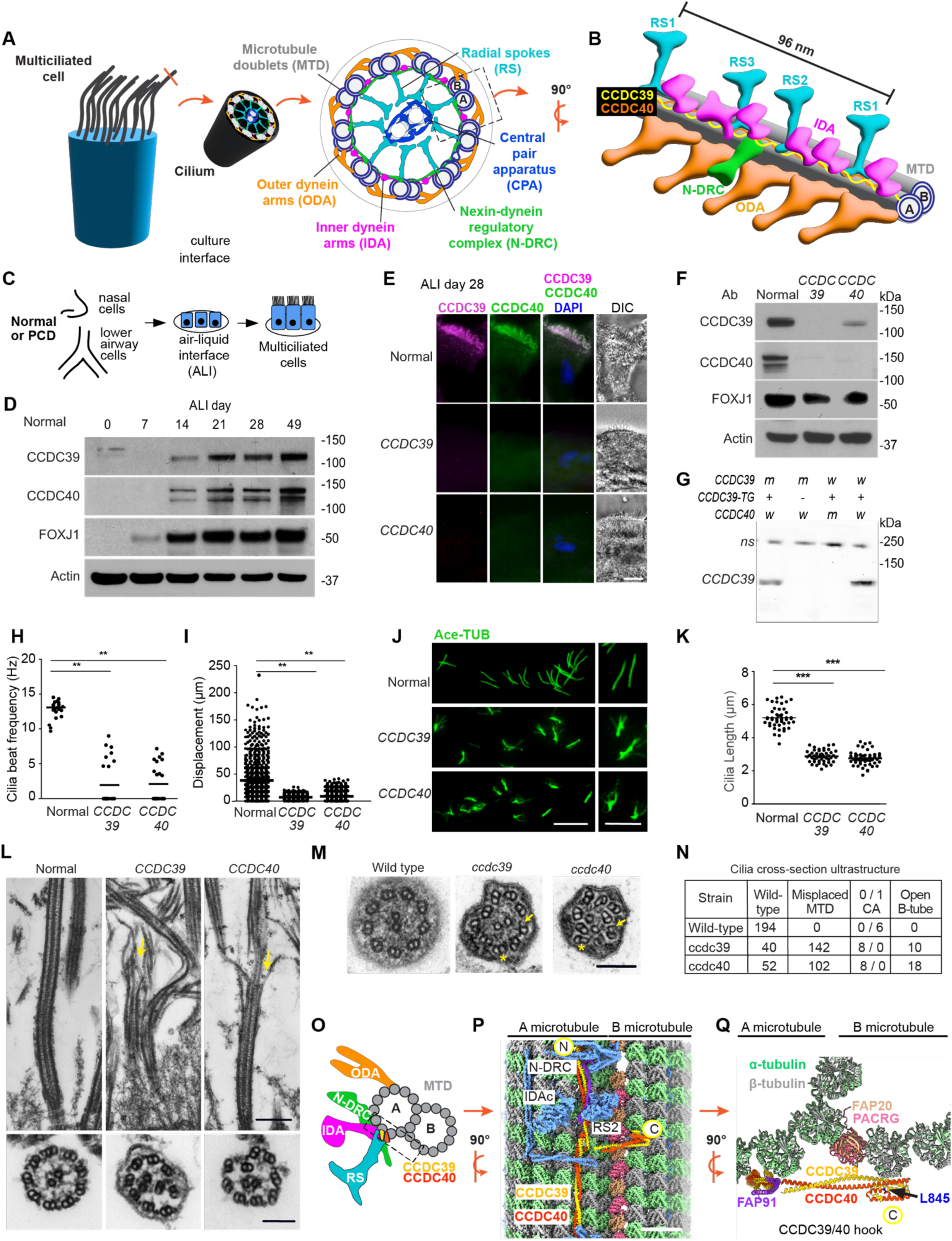
Loss of ciliary microtubular integrity of human multiciliated cells in *CCDC39* and *CCDC40* variants. **(A)** Diagram of multiciliated cell, motile cilium cross-section with the nine microtubule doublets (DMT) linked to the major complexes. Dashed rectangle is expanded in Figure 1B. **(B)** Diagram of a 96-nanometer unit showing the location of the CCDC39/CCDC40 heterodimer (yellow and orange ribbon) on the DMT. The major ciliary complexes are shown. **(C)** Scheme for obtaining primary airway epithelial cells for culture at air-liquid interface (ALI). Cells are from normal individuals and those with *CCDC39* and *CCDC40* variants. **(D)** Immunoblot (IB) detection of CCDC39, CCDC40, and FOXJ1 in normal ALI cultured cells during differentiation. **(E)** Immunofluorescent (IF) detection of CCDC39 and CCDC40 in normal and variant cells cultured at ALI 28. Bar=10 μm (WU 182, WU146, respectively and in panels H-L). **(F)** IB detection of CCDC39 and CCDC40 in normal and human variant cells. **(G)** IB detection of CCDC39 and CCDC40 in *Chlamydomonas* wild-type and *ccdc40* mutant. **(H)** Cilia beat frequency in normal (n=2) and *CCDC39* and *CCDC40* variant cells. **(I)** Cilia transport of microbeads on the surface of well-differentiated normal, C*CDC39* and *CCDC40* variant cells. Each point represents one bead. **(J)** IF detection of acetylated α-tubulin (Ace-TUB) in cilia isolated from human normal and variant cells. Detail shows examples of cilia with splaying in variants. Bar=10 μm (left) and 5 μm (right) **(K)** Quantification of cilia length from Panel J; n=47-50 cilia measured for each genotype from normal (n=3), *CCDC39* and *CCDC40*. **(L)** Transmission electron microscopy (TEM) of cilia from normal and variant cells showing the microtubules in longitudinal and cross section. Bar=500 nm (top) and 100 nm (bottom). **(M)** TEM of cross-sections of cilia isolated from *Chlamydomonas* wild-type, *ccdc39*, and *ccdc40*. Asterisks indicate disorganized DMT. Arrows indicate an opening of the B-microtubule. **(N)** Quantification of open B-microtubules from panel M; n=200 of each genotype. The total loss of the central apparatus in the *ccdc39* and *ccdc40* mutant strains is significantly different from wild-type (p<0.0001) by chi-squared and Fisher’s exact testing. **(O)** Diagram of a cross-section of a DMT showing the ciliary complexes and the location of CCDC39/ CCDC40 on the DMT. Structures are as in panels A and B. The box outlined represents the region viewed at 90^0^ in panel P. **(P)** Three-dimensional structure of the *Chlamydomonas* axoneme resolved by cryo-EM showing CCDC39 and CCDC40 and indicated proteins on the surface of the DMT. The N terminus of CCDC39/40 is labeled (N). The C-terminus extends from the A- to the B-tubule (C). Bar=10 nm. **(Q)** Detail of CCDC39/40 hook region from a predicted conformation of the C-terminal region of the CCDC39/40 heterodimer spanning from the A- to the B-microtubule. L845 is the location of the mutant residue in the temperature-sensitive *ccdc39 Chlamydomonas* mutant. **In H, I, K.** The bar indicates the medium. Differences between groups were determined using Kruskal-Wallis with Dunn’s Multiple Comparison Test; *p<0.05, **p<0.01 are shown.

Insufficient cilia motor function in humans results in altered organ laterality, infertility, and respiratory tract infection, which are hallmarks of the genetic disease primary ciliary dyskinesia (PCD)^22^. Pathogenic variants in nearly 60 genes cause PCD^23,24^; the phenotypes provide powerful tools for uncovering cilia biology^25,26^. Variants of cilia genes code for dynein arms, the radial spokes (RS), nexin dynein regulatory complex (N-DRC) or central apparatus^27–32^ (**Figure 1A, 1B**). Paired with model organism genetics, these variants provide structure-function relationships, but do not inform the ciliary assembly process. A common cause of PCD includes pathologic variants in *CCDC39* and *CCDC40*^33–35^. For unknown reasons, patients with *CCDC39* or *CCDC40* variants exhibit worse disease as indicated by decreased lung function and increased mucus plugging of airways, compared to individuals with variants in 20 other PCD genes^36–39^. Current dogma suggests that PCD is caused simply by ciliary dysmotility, however this concept does not explain the range of patient phenotypes. Thus, a second gap in understanding is how variants in different genes impact multiciliated cells.

CCDC39 and CCDC40 are alpha-helical proteins that form a heterodimer (CCDC39/40) repeating every 96-nm along the length of the DMT^10^ (**Figure 1B**). The dimer directly contacts most of the structures within the repeating unit^7,9^. These include proteins of the N-DRC and inner dynein arms (IDAs) that are missing in patients with *CCDC39* and *CCDC40* variants^33,34^. CCDC39/40 was suggested to be a molecular ruler for positioning of axonemal structures^40^. The ruler model proposed that CCDC39/40 measures a specific distance along the microtubule to direct the placement of IDAs and N-DRC^40^. An alternative model is that CCDC39/40 provides specific sites or “addresses” to dock ciliary complexes^7,41^.

We aim to identify how CCDC39/40 provides addresses and how its loss leads to severe lung disease. The variant cilia show reduced motility and are short. Proteomics of cilia from *CCDC39* human variants show a devastating loss of 90 ciliary proteins, which defines a large “connectome” dependent on CCDC39/40 for anchoring. Loss of this connectome leads to the absence of major ciliary structures, impaired ciliary integrity, and an unexpected cytoplasmic burden of unanchored proteins, leading to cell stress. The findings provide insight into the cilia-independent impact of pathologic genetic variants on patients with motile ciliopathies.

## Results

### Loss of ciliary integrity in *CCDC39* and *CCDC40* variants

To understand the function of the human CCDC39/40 heterodimer, we obtained freshly isolated, primary airway epithelial cells from the respiratory tract of normal individuals and those with biallelic pathologic variants in *CCDC39* or *CCDC40*. Most variants have frameshift mutations likely resulting in nonsense mediated decay of the RNA (**Table S1**). To avoid secondary insults on cilia that are caused by the inflamed airway environment in PCD, we isolated and expanded the airway basal progenitor cells in cultures with medium supplemented with antimicrobial drugs, then differentiated cells using the air-liquid interface condition (ALI) to generate multiciliated cells (**Figure 1C**). The onset of CCDC39 and CCDC40 expression during ALI differentiation coincides with the expression of the cilia transcription factor FOXJ1 (**Figure 1D**). The specificity of antibodies to CCDC39 and CCDC40 used for these studies was validated in cultured cells obtained from individuals with pathologic *CCDC39* or *CCDC40* variants by immunofluorescence and immunoblot analysis (**Figure 1E, 1F**). In some human variants we found that the unaffected partner protein is absent and in others, present by immunoblot analysis and is likely retained in the cytoplasm (**Figure 1F, Figure S1A, S1D**). CCDC39 protein is absent in the cytoplasm of the *Chlamydomonas ccdc40* mutant (**Figure 1G**), which suggests that formation of the heterodimer enhances stability. Functional analysis of cilia in *CCDC39* and *CCDC4*0 variant human cultures show most are immotile or flicker (**Figure 1H and Videos V1, V2)**. The surface transport of microbeads in variants is markedly decreased as expected (**Figure 1I; Figure S1B, S1C; Videos V3-5**).

We previously observed that cilia of *Chlamydomonas* with null alleles in *ccdc39* and *ccdc40* were short with ciliary splaying^42^. Cilia in human *CCDC39* and *CCDC40* variants are approximately half the length of normal, when measured intact on the surface of cultured cells or following treatment of cells with a detergent-containing buffer that removes cilia from cells (**Figure 1J, 1K**). Immunostaining using an anti-tubulin antibody shows splaying in some cilia isolated from variants, likely caused by the loss of the nexin-dynein regulatory complex that links the DMT pairs. TEM of human variant axonemes in cross-section show the loss of organization of the nine DMTs as reported previously^33,34,43^. Notably, imaging the length of the cilia showed splayed microtubules (**Figure 1L**), which was not previously described in PCD patients. Analysis of TEM images of *Chlamydomonas ccdc39* and *ccdc40* identify other ultrastructural defects including misplaced DMTs and open B-tubules at the inner junction (**Figure 1M, 1N)**, indicating the extensive loss of DMT integrity.

To determine how loss of CCDC39 or CCDC40 in the variants may lead to ciliary splaying and therefore instability, we used single particle cryo-EM of cilia from wild-type *Chlamydomonas* cells. We previously reported that CCDC39/40 is restricted to the groove between protofilaments A02 and A03, adjacent to IDAs, RS, and the N-DRC (**Figure 1O, 1P**)^10^. Previously, we were unable to resolve the C-terminus. With improved cryo-EM data processing and 3D classification, the last ∼30 residues of the C-termini of CCDC39/40 are now resolved near the inner junction (**Figure 1P**). They extend as a hook from the A- to the B-tubule, forming a 4-helices bundle that agrees well with the AlphaFold prediction (**Figure 1Q**). The importance of the A-to-B tubule extension was illustrated by analysis of a *CCDC39* variant (WU152), which carries a frameshift that is predicted to truncate the last 32 amino acids and shows persistence of CCDC39 and CCDC40 protein (**Figure S1D**). This individual has clinically severe PCD; cilia show lowered expression of CCD39 and CCDC40 and beat frequency (**Figure S1E, S1F**), suggesting an essential role for the CCDC39 C-terminal structure. A temperature-sensitive mutation in *ccdc39* (L845P) is also within this hook region (**Figure 1Q**) and causes loss of the heterodimer at the restrictive temperature but reduced function at intermediate temperatures^42^. Taken together, the loss of CCDC39/40 function accounts for a unique defect in ciliary structure.

### CCDC39/40 is required for assembly of a connectome of ciliary proteins via CARPs

Prior analysis of *Chlamydomonas* cilia in *ccdc39* and *ccdc40* mutants identified deficiencies in IDAs, N-DRC and tektin^40,42^. Other proteins were preserved, although studies were limited by antibody availability^42^. Proteomics of isolated cilia would provide a comprehensive assessment of the role of CCDC39/40. Cilia obtained from ALI preparations of cells from four normal donors identified 440 proteins for comparison to cilia from a *CCDC39* variant individual (**Figure 2A, 2B**).

**Figure 2.**
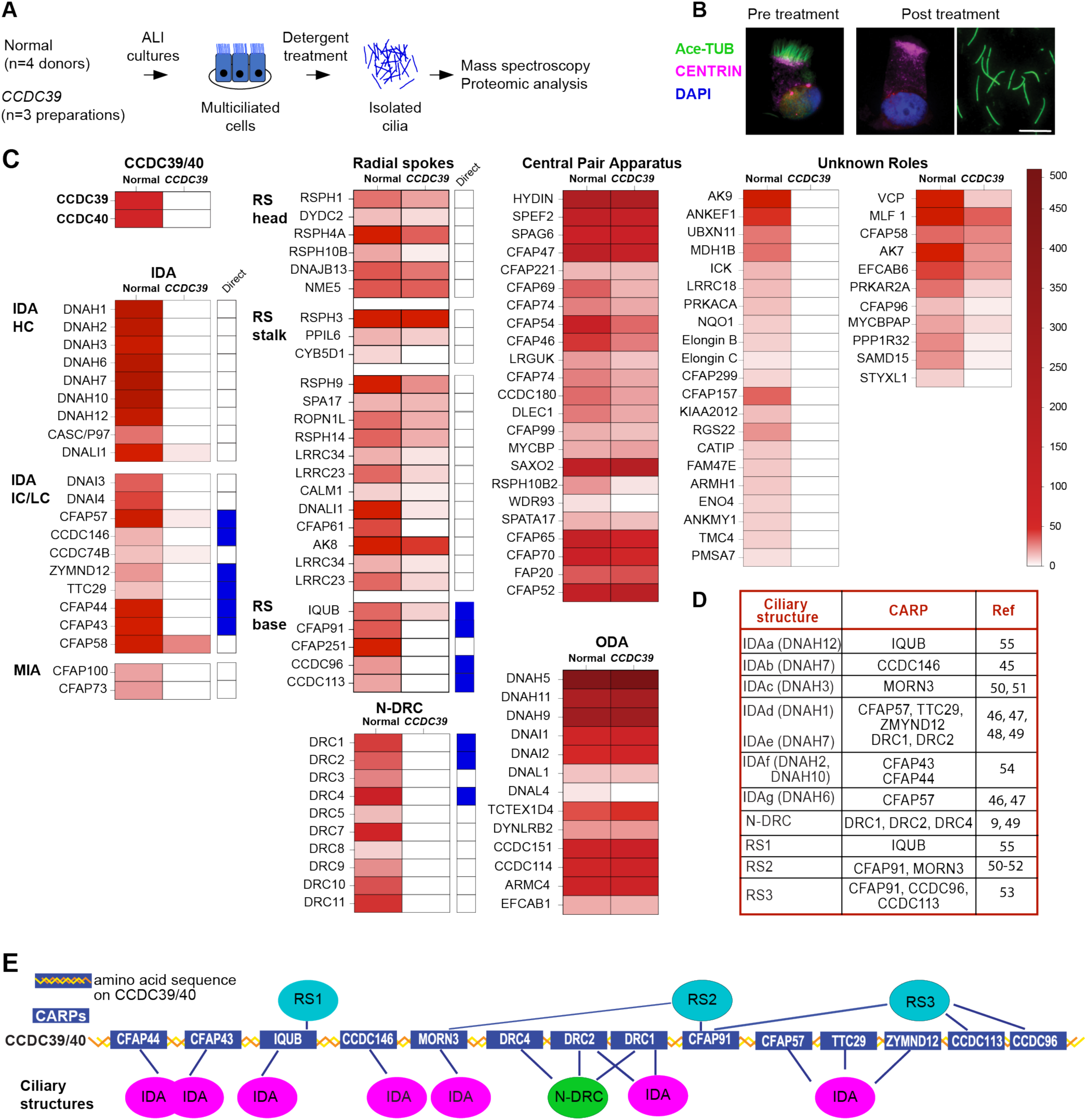
CCDC39/40 is required for assembly of a connectome of ciliary proteins. **(A)** Scheme of cilia isolation from cultured human cells from normal (n=3) and *CCDC39* variant (WU182) analyzed by mass spectrometry. **(B)** Immunofluorescent (IF) detection of Ace-TUB and basal body protein centrin in cells after cilia isolation. Bar=10 μm. **(C)** Heat map of mean levels of proteins normalized for tubulin in the *CCDC39* variant compared to normal cilia. Proteins with direct CCDC39/40 contact noted by blue box. **(D)** CCDC39/40 ciliary address recognition protein (CARP) with associated structure and references to experimental validation. **(E)** Diagram of the relationship of CCDC39/40 with CARPs and ciliary structures (see Figure 1A for abbreviations).

Proteomics of cilia from the *CCDC39* variant shows extensive loss of at least 90 proteins (**Figure 2C, Table S2)**. Outer dynein arms are retained, but many of the known structural proteins are lost or reduced. All the IDA proteins are missing, as are 9 of the 12 N-DRC proteins; DRC5, DRC6, DRC12 were not detected in the normal cilia. CFAP73 and CFAP100, which regulate the normal waveform via the two-headed IDA^44^, are missing. Many radial spoke (RS) proteins are missing or reduced, as are several proteins within the central apparatus (CA), consistent with the loss of this structure in the cross-sections of the *ccdc39* or *40* mutants (**Figure 1L-1N**). Many proteins with unknown roles and locations are also depleted or reduced (**Figure 2C**). Inspection of single particle cryo-EM reconstructions shows that at least 13 of the missing proteins are in direct contact with the CCDC39/40 heterodimer (**Figure 2C**)^7–12,14,45–55^. Each of these 13 missing proteins, plus MORN3, which is not present in our proteomics, is linked to a major cilia complex (**Figure 2D**). Due to this special arrangement, we refer to these 14 proteins as ciliary address recognition proteins (CARPs). The CARPs are diagrammed to indicate their position relative to ciliary structures (**Figure 2E**).

### Ciliary radial spokes are differentially depleted in *CCDC39/40* variants

RS act as a mechanotransducer to control ciliary waveform^9,56^. *Chlamydomonas* has two full radial spokes (RS1, RS2) and a shorter third RS (RS3) structure while humans have three full radial spokes within each 96-nm repeat unit (**Figure 3A**)^7,9,15,57,58^. Each RS contains unique proteins, as well as shared head and stalk proteins.

**Figure 3.**
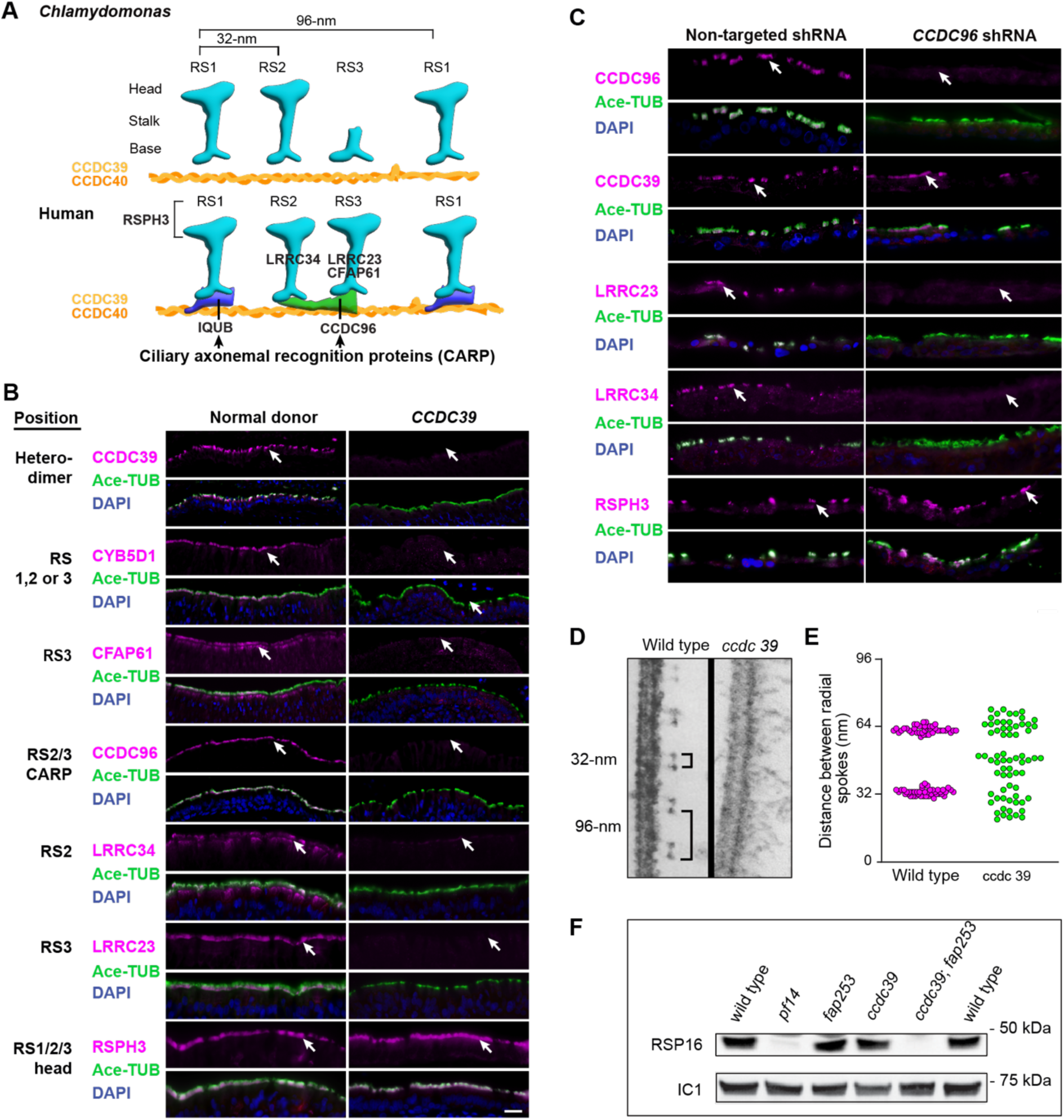
Ciliary radial spokes are differentially depleted in *CCDC39/40* variants. **(A)** Diagram of the radial spokes (RS) in *Chlamydomonas* and human in a 96-nm repeat along the DMT. *Chlamydomonas* has two complete and one partial RS. The human has three complete RS proteins at the bases. RSPH3 is shared by all RS. **(B)** Immunofluorescent (IF) detection of Ace-TUB, CCDC39, and RS proteins in normal and *CCDC39* variant airways. Arrow shows indicated protein. Bar=25 μm. **(C)** IF detection of Ace-TUB, CCDC39, and radial spoke (RS) proteins in normal cells transduced with non-targeted or *CCDC96*-specific shRNA. Arrow shows indicated protein. Bar=25 μm. **(D)** TEM of isolated *Chlamydomonas* wild-type and *ccdc39* axonemes. Repeating distances of radial spokes within a 96-nm repeat (32 nm) and between repeats (96 nm) are indicated. **(E)** Quantification of distance from panel D. **(F)** Immunoblot detection of RS head protein RSP16 shared by all RS in *Chlamydomonas* strains.

We found that radial spokes are differentially depleted in the *CCDC39/40* variants. Cryo-EM analysis shows that each of the three radial spokes is anchored to CCDC39/40 by different CARPs^7,9^. The loss of RS2 and RS3 proteins were investigated in *CCDC39* and *CCDC40* variants based on the availability of antibodies (**Figure 3B, Figure S2A**). By proteomic analysis the RS2 and RS3 CARP, CFAP91^7,52^ is absent. Likewise, RS3-specific CARP, CCDC96, is absent and confirmed by immunostaining (**Figure 2B, Figure S2A**). Immunostaining confirms the reduction of RS2 stalk protein LRRC23, RS3-specific proteins CFAP61 and LRCC34. RSPH3, a shared RS protein, is present in the cilia of normal and variant cells by immunostaining, consistent with the assembly of some radial spokes. We used depletion of the RS3 CARP, CCDC96 in human cells to ask about its role for RS3. *CCDC96* shRNA resulted in loss of CCDC96 and decrease in the surface transport of beads (**Figure S2B-S2D**). CCDC39 and RSPH3 were preserved (**Figure 3C**). There was loss of both LRRC34 and LRRC23, suggesting that CCDC96 plays a role in both RS2 and RS3 (**Figure 3C**).

Additionally, analysis shows that RS1 CARP (IQUB) is diminished. *Chlamydomonas* was used to confirm the retention of RS1 and loss of RS2 in the *ccdc39* mutant. Analysis of longitudinal TEM images of *ccdc39* show loss of the periodicity and rigidity of RS (**Figure 3D and 3E**). RSP16, a shared radial spoke head protein, is present in cilia isolated from both *ccdc39* and *fap253* (CARP and *IQUB* ortholog) by immunoblot analysis, confirming the presence of RS (**Figure 3F**). However, immunoblots of cilia from the *ccdc39*; *fap253* double mutant lacks RSP16, indicating the complete loss of RS. The finding suggests that in the absence of CCDC39/40, RS1 still binds the microtubule lattice without address information but with variable positioning. Together, our observations suggest that CCDC39/40 provide unique addresses for the attachment of all RS via designated ciliary address recognition proteins (CARP).

### Altered microtubule inner proteins in *CCDC39/40* variants

Microtubule inner proteins (MIPs) are a network of proteins within the lumen of the A and B tubules that show 8-, 16-, and 48-nm periodicities^10,12,13,59^. Some MIPs directly contact outer surface proteins. In our proteomics of *CCDC39* variant cilia, three of the four tektin proteins (TEKT2,3,4) are decreased compared to normal cilia (**Figure 4A**). TEK3 does not contact CCDC39/40 but was absent in variant cilia by proteomics and verified by immunolocalization (**Figure 4A, 4B**). Decreased TEK2 and TEK4 may result from an inability to form the TEK3-TEK2 and TEK3-TEK4 heterodimers^7^. Additionally, CFAP90 (inner junction) and CFAP141 (outer junction) are absent in our proteomics (**Figure 4A**). CFAP90 is localized at the inner junction^10^, near the hook of the CCDC39/40 heterodimer (**Figure 1Q**). There was no difference in normal and *CCDC39* in the total number of peptides of other MIPS (**Figure 4A**). The 48-nm MIPS are present in the cilia by immunostaining (**Figure 4C**).

**Figure 4.**
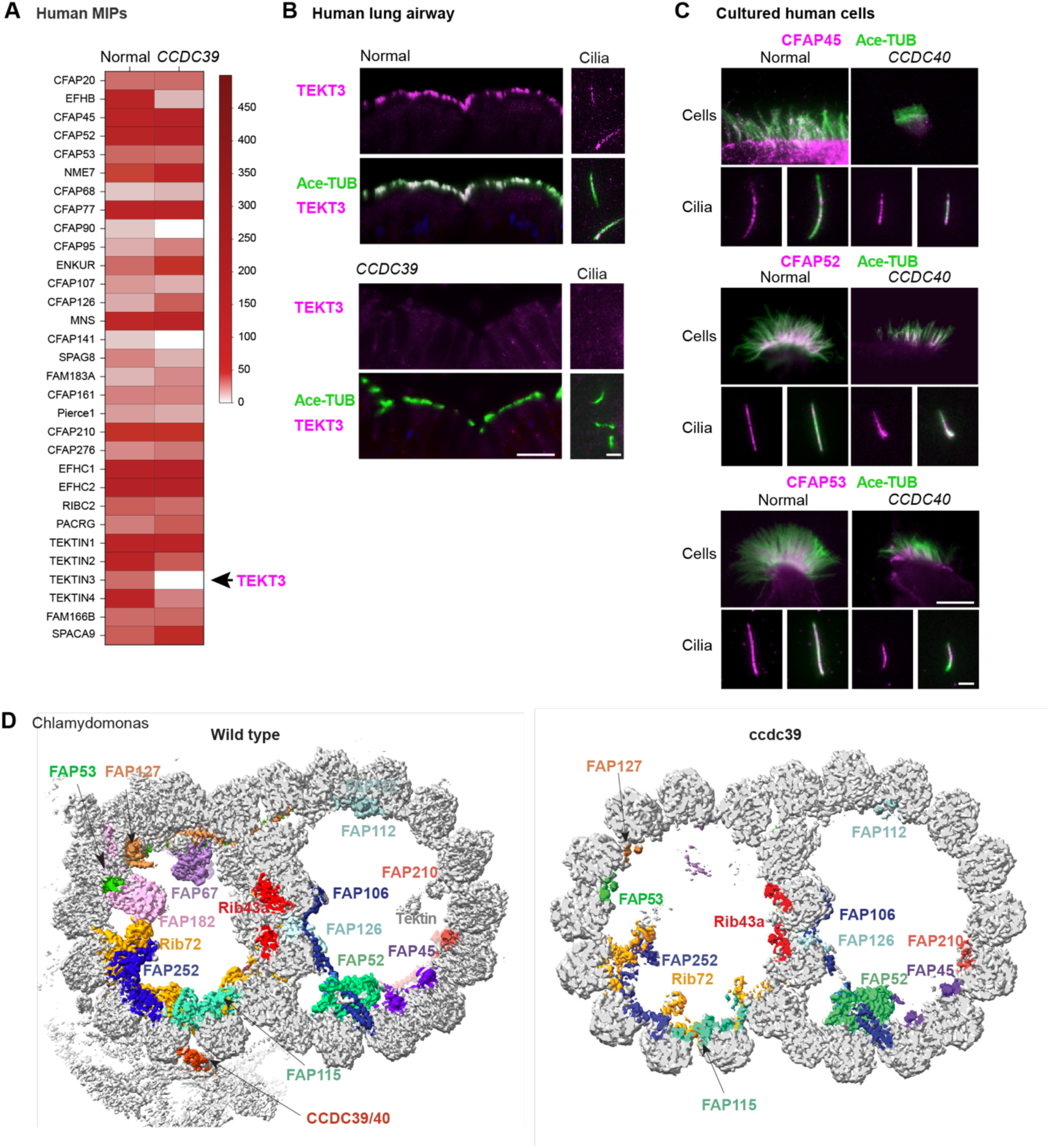
Disorganization of microtubule inner proteins (MIPs) in *CCDC39/40* variants. **(A)** Proteomic analysis of MIPs in normal and *CCDC39* variant human cilia **(B)** Immunofluorescent (IF) detection of TEKT3 in human cells (*CCDC39*, WU182). Airway, Bar=25 μm; cilia, Bar=2 μm. **(C)** IF detection of MIPs in cilia cultured cells (*CCDC40*, WU146). Cells, Bar=5 μm; cilia, Bar=2 μm. **(D)** MIPs in wild-type and *ccdc39-2 Chlamydomonas* identified by cryo-EM. DMT structure from wild-type has 48-nm periodicity applied and *ccdc39-2* has 16-nm periodicity applied.

By cryo-EM data of *Chlamydomonas* cilia, the 48-nm MIPs are disrupted in the *ccdc39* mutant compared to wild-type (**Figure 4D**). However, the MIPs with 8-nm and 16-nm periodicity are largely preserved (**Figure 4D**). The disorganization of *Chlamydomonas* MIPs contrasts with the structural analysis that only found the loss of tektin in human *CCDC39/40* variants^7^. Like in our human proteomics, tektin was diminished in *ccdc39*^42^ and in N-DRC mutants (*DRC*1 and *DRC2*)^60^ (**Figure 4D**). We predict that missing and misplaced MIPs in the variants contribute to abnormal ciliary function.

### CCDC39/40 connectome localization during cilia assembly

Our data suggest that the CCDC39/40 heterodimer creates a set of addresses for anchoring the CARPs on the ciliary axoneme. By this hypothesis, we predict a temporal pattern of trafficking, whereby the heterodimer would enter the cilia early, followed by the CARPs and finally those complexes that preassemble in the cytoplasm^61^. Guided by stages of ciliogenesis (**Figure 5A**), we examined trafficking of CCDC39/40 in normal cells. During early stages of multiciliated cell differentiation marked by centriole amplification, CCDC39 and 40 are expressed present in the cytoplasm, but do not directly colocalize with centrioles (**Figure 5B**). Just prior to cilia assembly, CCDC39 and CCDC40 are enriched at the apical domain but infrequently colocalize (**Figure 5C**). Subsequently, CCDC39/40 colocalize in short cilia as they begin to grow (**Figure 5C**).

**Figure 5.**
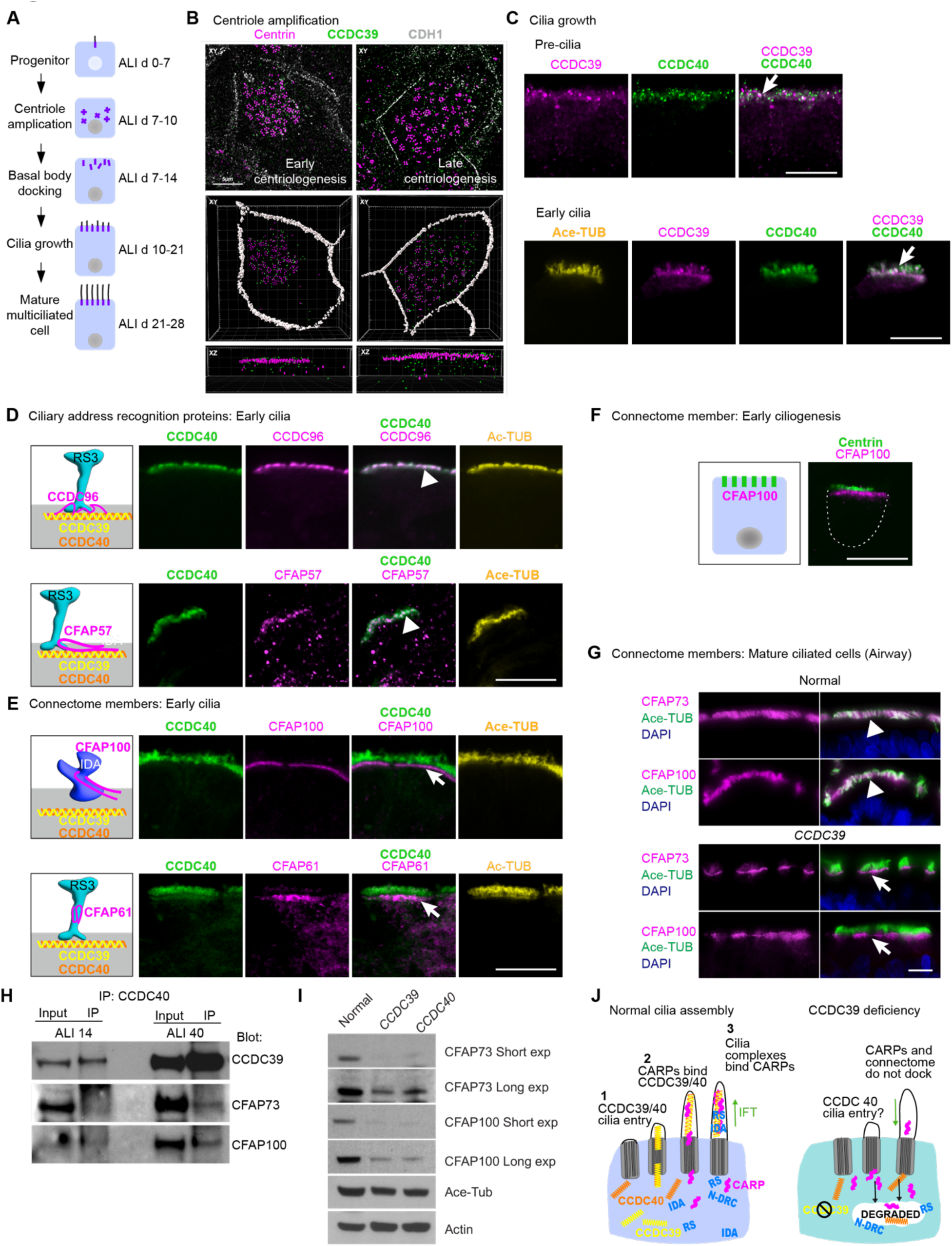
CCDC39/40 traffics independently of connectome proteins. **(A)** Scheme of motile ciliogenesis in air-liquid interface (ALI) cultures. **(B)** Immunofluorescence (IF) detection of basal body protein centrin, CCDC39, and CDH1 during basal body amplification. **(C)** IF detection of CCDC39 and CCDC40 at pre-cilia stage (arrow indicates co-localization). **(D)** IF detection of Ace-TUB and CCDC40 with proteins that contact CCDC39/40 in cells with emerging cilia (arrow indicates co-localization). **(E)** IF detection of Ace-TUB and CCDC40 with proteins that do not directly contact CCDC39/40 in cells with emerging cilia. **(F)** IF detection of connectome member CFAP100 and basal body protein Centrin during early cilia growth. **(G)** IF detection of Ace-TUB, CFAP73, and CFAP100 in mature normal and *CCDC39* cells. **(H)** Immunoprecipitation of CCDC40 in cells from early (ALI 14) and late (ALI 40) stages. **(I)** IB of CCDC39, CCDC73, and CCDC100 in normal, and variant mature cells. Short (S) and long (L) exposure. **(J)** Proposed schema of cilia assembly in normal and *CCDC39* variant. Bar=5 μm in B-G

We then tested the temporal pattern of trafficking of CCDC39/40, the CARPs, and the connectome during normal cilia assembly. The RS3 CARP (CCDC96) and IDA CARP (CFAP57) are localized with CCDC40 in very short cilia (**Figure 5D**). In contrast, non-CARPs (CFAP100, CFAP61), remain within the cytoplasm, near the basal bodies (**Figure 5E, 5F**). Localization of these connectome proteins in normal multiciliated cells in early cilia assembly may identify a staging site for later transport into the cilia. In later stages of ciliogenesis, connectome proteins are in the cilia (**Figure 5G**). In normal cells, immunoprecipitation with CCDC40 pulls down connectome proteins CFAP100 and CFAP73 to a greater degree at late compared to early stages of cilia growth (**Figure 5H**).

In *CCDC39/40* variants, connectome proteins remain within the apical domain (**Figure 5G**), a location similar to the assembly staging site in normal early cilia (**Figures 5E**). Ultimately, the unbound connectome members are degraded (**Figure 5I**). We suggest cilia assembly proceeds with CCDC39/40 binding the DMTs, followed by CARP binding to the heterodimer, and subsequently, the remaining connectome. We cannot rule out that CARPs are co-transported with CCDC39/40. In sum, there is an orderly assembly sequence, dependent on CCDC39/40 and in the absence of the heterodimer, the proteins remain at the staging site or degrade (**Figure 5J**).

### Single cell RNA transcriptomics reveals increased proteostasis in *CCDC39/40* variants

To interrogate multiciliated cell dysfunction from CCDC39/40 loss, we used single cell RNA sequencing (scRNAseq). Transcriptomes were compared from normal (n=6) and variant (n=3) cell cultures. Cell types were identified using unsupervised clustering and annotated with known airway epithelial markers (**Figure 6A**)^17^. *CCDC39* and *CCDC40* are primarily expressed in the cluster of mature multiciliated cells (Cil3) (**Figure 3SA**). In this cluster, over 1800 genes are differentially expressed in the variants compared to normal (**Extended Data Table**). Enrichment analysis of differential gene expression in multiciliated cells (Cil3), identified cilia structure and function, proteostasis, cell stress, oxidative phosphorylation, and Notch signaling pathways (**Figure 6B, Figure S3B-S3F**). Genes upregulated have roles in: (1) proteostasis, including components of the E3 ubiquitin ligase complex^62^, multiple proteasome subunits, heat shock chaperone proteins that handle unfolded proteins and aggregates^63^ (**Figure S3B, S3E**); (2) cell stress response genes that include *SAA1*, *SAA2*, and *SAA4*, which are secreted acute phase reactants in sterile inflammation^64,65^ (**Figure 6B**), (3) mitochondrial complex I subunits of NADH ubiquinone oxidoreductase and complex V subunits of ATP synthase indicating upregulation of ATP production (**Figure S3C**); and (4) *HES1*, a target of Notch signaling that increases secretory cell differentiation and airway secretory cell proteins^66–68^ (**Figure 6B**). Together, these data indicate the presence of cellular stress in the variant cells, possibly due to the need to degrade the large burden of undocked connectome proteins. We examined the proteasome in variant cells using an antibody against PSMB6, a component of the 20S core subunit (**Figure 6C**). PSMB6 is located throughout in the cytoplasm of normal cells. In contrast, PSMB6 is localized just beneath the cilia in the variant cells. The pattern is coordinate with the location of undocked connectome members in variant cells (**Figure 5H**), suggesting a site-specific requirement for proteostasis. Consistent with our finding, increased proteostasis leads to increased ATP demand and oxidant production. These factors likely cause cell stress that may drive Notch-signaling^69–72^.

**Figure 6.**
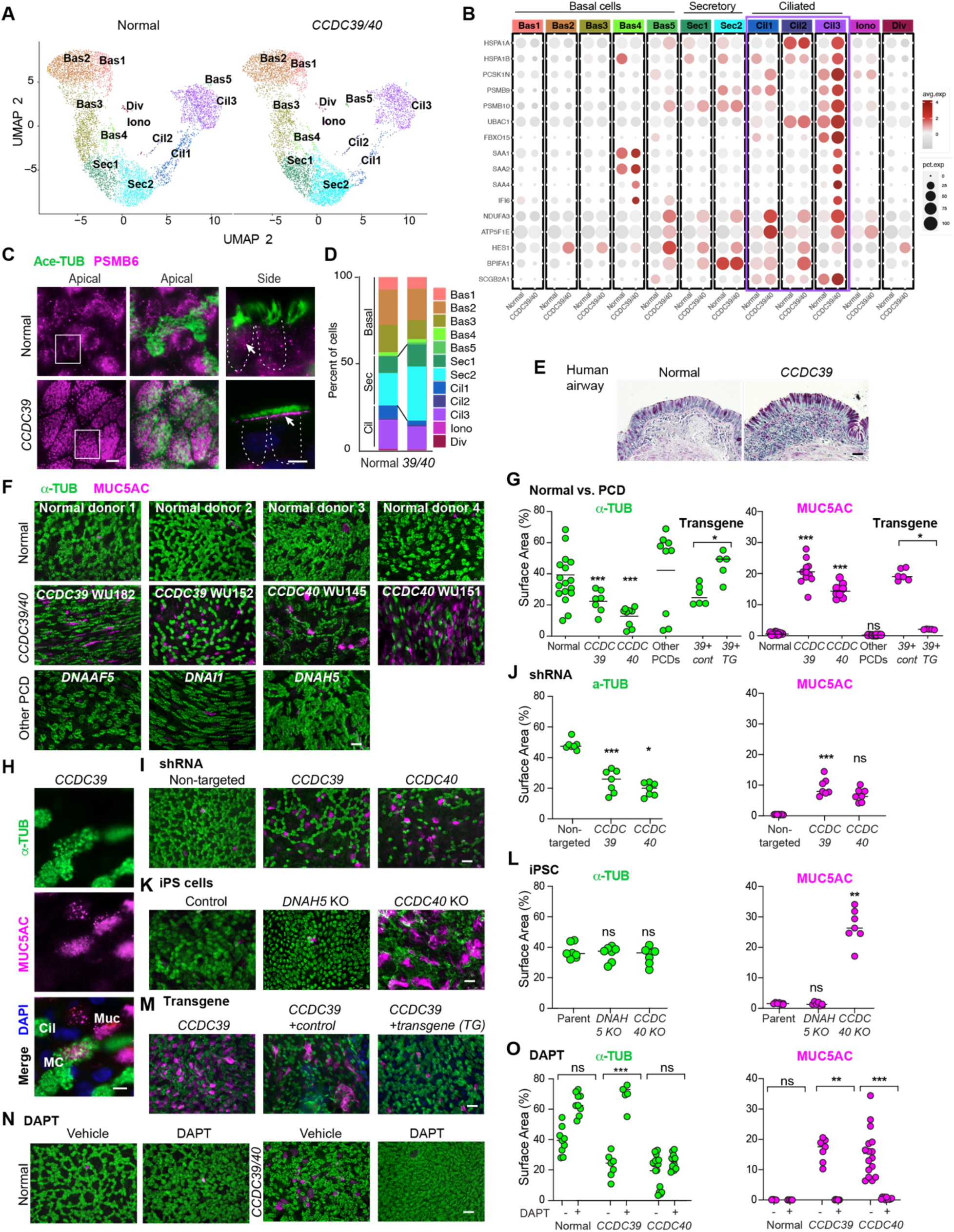
CCDC39/40 variant affect differentiation via Notch signaling. **(A)** UMAP reductions of normal (n=6) and *CCDC39/40* variant (*CCDC39*, WU182 and WU 157; CCDC*40* WU146) differentiated airway epithelial cells showing clusters for basal (bas); secretory (sec); multiciliated (cil) cells. **(B)** Dotplot shows manually annotated, differentially expressed genes in each cluster that show significantly increased expression in the variant compared to multiciliated cells. Ciliated cells are in purple. **(C)** Immunofluorescent (IF) detection of Ace-TUB and PSMB6 in normal and *CCDC39* (WU182) variant cells. Box identifies an example of PSMB6 in multiciliated cells. Arrow indicates the location of PSMB6 in the cytoplasm. **(D)** Quantitation of cell numbers in each cluster from A. Comparison of Sec2 normal vs. *CCDC39*/*CCDC40* (*39*/*40*) variant, p=0.0025. **(E)** Immunohistochemistry for detection of PAS-positive mucous cells in the airway from a lung of a normal donor and a *CCDC39* variant. **(F)** IF detection of α-tubulin (α-TUB) and MUC5AC in cell cultures from unique normal donor (n=4), unique *CCDC39/40* variants (n=4), and PCD variants *DNAAF5*, *DNAH5*, *DNAI1* (n=1 per genotype). **(G)** Quantitation of normal cells with genotypes *CCDC39*, *CCDC40* and other PCD variants from Panels F and L. *CCDC39* variant cells expressing control (Cont) and the *CCDC39* Transgene (*TG*) are compared. **(H)** IF detection of α-TUB and MUC5AC in *CCDC39* variant cells. Cil, multiciliated cell; Muc, MUC5AC cell; MC, Multiciliated-MUC5AC cell. **(I)** IF detection of MUC5AC and Ace-TUB in normal cells transduced with non-targeted *CCDC39* or *CCDC40* shRNA. **(J)** Quantitation of panel I. Normal cells transduced with non-targeted, *CCDC39*, or *CCDC40* shRNA are compared. **(K)** IF detection of α-TUB and MUC5AC in iPSC cells that are cells derived from the control original iPSC from a normal donor, or CRISPR-Cas9 mediated deficiency of *DNAH5* or *CCDC40* in the normal iPSC. Bar=25 μm **(L)** Quantitation of panel K. Each point represents a microscope field of cells from the control original iPSC line, *DNAH5* CRISPR knockout, or *CCDC40* CRISPR knockout. The control cells are compared to the knockout lines. **(M)** IF detection of α-TUB and MUC5AC in transgene (*TG*) rescued variant cells transduced with a control (*Cont*) or *FOXJ1-CCDC39* transgene. Quantitation of M is in panel G. **(N)** IF detection of α-TUB and MUC5AC in unique normal and *CCDC39* and *CCDC40* variant cultures (ALI >28) day then treated with vehicle or NOTCH inhibitor DAPT for two weeks. **(O)** Quantitation of panel N. Vehicle cells are compared to DAPT-treated for each group. Each point is the mean of fields of each condition in at least three independent experiments. Normal (n=3); *CCDC39* (WU182, n=2); *CCDC40* (WU146, n=2). The bar indicates the medium. Differences between groups were determined using Mann-Whitney in D, G, O or Kruskal Wallis with Dunn’s Multiple comparison test in J and L; ns=non-significant, *p<0.05, **p<0.01, ***p<0.001. Bar=5 μm in C, H; Bar=25 μm in E, F, K, I, M, N.

### *CCDC39/40* variants switch fates in airway epithelial cells

Comparing variant and normal cells, scRNAseq analysis shows an increased number of secretory cells (Sec1 and Sec2) (**Figure 3D**), suggesting a shift in differentiation. The balance of airway multiciliated versus secretory cell differentiation is controlled by Notch^66,73^. *In vivo*, chronic airway inflammation is found in patients with asthma, cystic fibrosis, chronic obstructive pulmonary disease (COPD). These airways have increased mucous cells (also called goblet cells), marked by increased MUC5AC, a gel-forming, secreted mucin^74^. There is also an abundance of mucous cells in tissue samples obtained from an airway of a patient with a *CCDC39* variant who underwent lung transplantation (**Figure 6D**). We examined the production of MUC5AC in ALI cultured cells. Control cultures show a high percentage of multiciliated cells, with few MUC5AC-expressing cells. In contrast, *CCDC39*/*40* variant cultures have significantly more MUC5AC-expressing cells at the expense of multiciliated cells (**Figure 6F, 6G**). Cultured cells from PCD patients with variants in *DNAAF5*, *DNAI1*, and *DNAH5* show significantly fewer MUC5AC-staining cells than the *CCDC39/40* cultures (**Figure 6F, 6G**). A small number of cells expressing both MUC5AC and cilia proteins are present in the *CCDC39*/*40* cultures and represent a transition state in cell differentiation^75^ (**Figure 6H)**.

It is possible that the chronic inflammatory environment in the airways of *CCDC39* and *CCDC40* patients account for the increased mucous cells *in vitro*, as observed in cultures of cells from COPD subjects^76,77^. Accordingly, we depleted *CCDC39* or *CCDC40* from normal donor airway cells using shRNA (**Figure 6I, 6J, Figure S4A-D**). *CCDC39* or *CCDC40* depleted cultures show increased numbers of MUC5AC-expressing cells and fewer multiciliated cells than controls. Thus, the shift in the differentiation phenotype is unlikely to be attributable to effects of a chronically infected, inflamed PCD airway. To exclude the possibility that the MUC5AC phenotype was the result of exposure of cells to an airway environment, we assessed cultures of inducible pluripotent stem cells (iPSC) that underwent *CCDC40* or *DNAH5* deletion by CRISPR-CAS9 editing^78^. Cells were differentiated to airway basal cells, then cultured using ALI conditions. *CCDC40* deficient cells show more MUC5AC-expressing cells compared to those with a *DNAH5* deletion or the control iPS cells (**Figure 6K, 6L; Figure S4E**). These findings suggest that the *CCDC39* or *CCDC40* patient cells or their airway basal cells are not altered by the environment. Finally, lentivirus-mediated delivery of a transgene expressing *CCDC39* under control of the *FOXJ1* promotor (Transgene, TG) in the *CCDC39* variant cells showed rescue compared to a control lentivirus (Cont) (**Figures 6M, 6G, Figure S5A-S5E)**. We observed a decrease in the percentage of MUC5AC cells and an increased percentage of ciliated cells (**Figure 6G, 6L**). Thus, the shift in cell fate likely occurs by the transition of multiciliated to secretory cells via a genetics loss of *CCDC39/40* and is consistent with increased mucus plugging of airways observed in computed tomography imaging of patients with *CCDC39/40* variants^39^.

To ask if the shift in differentiation from multiciliated to mucous cells in the *CCDC39/40* cultures was mediated by Notch, we inhibited Notch signaling with the γ-secretase inhibitor DAPT^79^. We delayed treatment of the variant cultures with vehicle or DAPT until after ALI day 28, when MUC5AC cells were increased. In cells treated with DAPT, the number of MUC5AC-expressing cells was markedly decreased and acetylated α-tubulin expressing cells was increased compared to a persistence of MUC5AC-expressing cells in vehicle-treated variant cells (**Figure 6N, 6O**). We conclude that Notch signaling is required to shift the differentiation of the multiciliated variant cells to mucous cells and that continuous inhibition of Notch signaling blocks the switch from multiciliated to mucous cells. As expected, the *CCDC39* and *CCDC40* variants remain immotile after treatment (**Figure S4F**). Taken together, CCDC39/40 deficiency results in loss of ciliary proteins, increased intracellular stress, activated Notch activity, and altered maintenance of multiciliated cell differentiation.

### *CCDC39/40* variants disrupt the periciliary barrier

We defined the intracellular impact of a failed connectome, but the effect of the altered cilia integrity remains unaddressed. In addition to ciliary motility, the cilia are an intrinsic component of the periciliary barrier. Seminal studies defined the periciliary barrier as containing a brush-like polymer of tethered mucins (MUC1, MUC4, MUC16) that surround the cilia and act as a selective filter to particle entry^74,80^. One striking feature of the *CCDC39/40* variants is ciliary splaying (**Figure 1L-N**). To better visualize the DMT splaying, we used scanning EM. Cells were untreated or treated with detergent-containing buffer that disrupts the ciliary membrane (**Figure 7A**). Normal cilia exhibit intact DMTs in both conditions. *CCDC39/4*0 variants show DMT splaying, particularly at the distal ends in the untreated condition and further unraveling of the DMT with detergent treatment (**Figure 7A-C**). Normal isolated cilia show little splaying regardless of treatment. In contrast, at least 15% of variant isolated cilia show splaying with no treatment, while about 50% splay when the membrane is removed. Splaying, as well as cilia length and cilia beat frequency were rescued using a *CCDC39* transgene (**Figures 7F-J, Supplement S5A-F; Video V6**).

**Figure 7.**
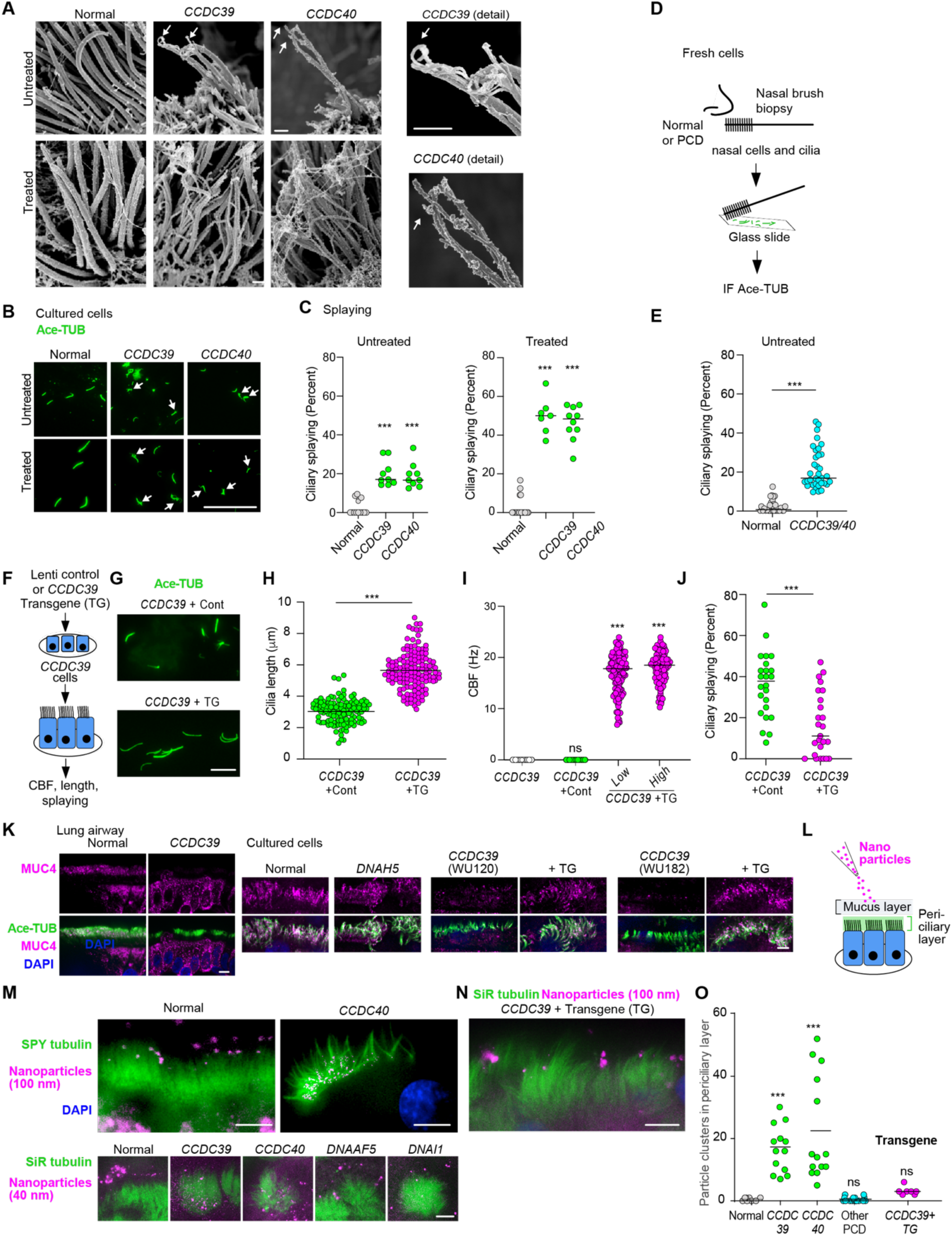
*CCDC39/40* variants have disrupted periciliary barrier. **(A)** Scanning EM of normal and variant (*CCDC39*, WU152; *CCDC40*, WU151) cilia from cells untreated and treated to disrupt the ciliary membrane. Bar=1 μm. (B) Immunofluorescent (IF) detection of Ace-TUB to identify splaying in cilia untreated and treated with buffer to remove cilia membranes from cultured cells. Bar=10 μm. (C) Quantitation of splaying in cilia from cultured cells from normal (n=2) and variant cells (*CCDC39*, WU152, WU182; *CCDC40*, WU146, WU151). (D) Scheme for isolation of fresh cilia by nasal brush biopsy and IF staining. (E) Quantitation of IF detection of splaying by Ace-TUB in cilia from normal (n=3) and variant cells (*CCDC39*, WU120, WU152; *CCDC40*, WU149, WU151). There was no difference between *CCDC39* and *CCDC40*. (F) Scheme of normal *CCDC39* transgene delivery to *CCDC39* variant cells (WU182) using control and *FOXJ1-CCDC39* rescue transgene lentiviruses (Lenti). (G) IF detection of Ace-TUB to identify cilia length in cells from *CCDC39* variant transduced with control (top) and rescue (bottom) lentiviruses in panel F. Bar=10 μm. (H) Quantification of cilia length in *CCDC39* variant cells from panel G, transduced with control or transgene (TG) lentivirus; n=150-200 cilia per condition in over 20 fields. (I) Quantitation of cilia beat frequency (CBF) in *CCDC39* variant cells and cells transduced with control lentivirus, low or high concentration of the *CCDC39* TG. (J) Quantification of cilia splaying in cells transduced with control and TG. n=300-500 cilia per condition from over 20 fields. (K) IF detection of MUC4 in the periciliary region of cultures from normal, *DNAH5* (WU165) variant, and two *CCDC39* variant cells non-transduced and following transgene rescue. Bar=5 μm. (L) Scheme of periciliary barrier assessment performed by addition of microparticles to the apical surface for 15 min prior to fixation and imaging. (M) Fluorescence particles relative to cilia in cultures from normal and variant (*CCDC39*, WU182; *CCDC40*, WU146; *DNAAF5*, WU108; *DNAI1*, WU103). Bar=5 μm. (N) Detection of fluorescent particles in *CCDC39* variant (WU182) transduced with the rescue *CCDC39* transgene as in panel F, Bar=5 μm. (O) Quantitation of fluorescent particles in the periciliary space from panels and L and M. Differences in the indicated conditions were determined compared to normal cells. In C, E, I, N the bar indicates the medium. Difference between groups determined by Kruskal-Wallis and with a Dunn’s Multiple Comparison Test. In H the unpaired two-tailed t-test. In J, the Mann-Whitney test ns=non-significant; ***p<0.001.

To determine the effect of the respiratory tract environment, we examined the extent of cilia splaying from freshly obtained samples (**Figure 7D**). Using non-detergent conditions, the median number of splayed cilia in freshly obtained and cultured samples were similar (**Figure 7C, 7E**). Surprisingly, untreated, fresh samples were significantly more variable in their splaying phenotype (F=0.03) (**Figure 7D, 7E)**, suggesting that the respiratory tract environment destabilizes the ciliary membrane.

The periciliary barrier relies on both intact cilia and inter-ciliary tethered mucins^74,80^. MUC4 is present within the ciliary region of normal and *DNAH5* variant cells but absent in the periciliary layer of cells with *CCDC39* variants (**Figure 7K**). Its presence in the periciliary layer of the *CCDC39* variant can be rescued by the transgene (**Figure 7K**), suggesting that the periciliary barrier is abnormal due to the variant. While it is difficult to visualize the polymer-like structure of this barrier, it can be functionally defined as being impervious to particles that are greater or equal to 40 nm in diameter^80^. To test the barrier function of variant multiciliated cells, we labeled the cilia of live cells using a fluorescent reporter, SPY-tubulin. Fluorescent-coated spherical particles (100 nm or 40 nm) were added to the apical surface (**Figures 7L**). The particles penetrate the periciliary space of the *CCDC39* and *CCDC40* variant cells, but not those of control cells (**Figures 7M, 7O**). Likewise, there is no detectable entry of microparticles in cells from individuals with PCD due to *DNAI1* or *DNAAF5* variants, which also have non-beating cilia. The periciliary space function in *CCDC39* cells was restored with the transgene (**Figures 7N, 7O**). The findings suggest that there is an incompetent periciliary barrier in *CCDC39* and *CCDC40* variant cells, increasing susceptibility to injury and infection that is not related to ciliary beating, contributing to severe lung disease in affected patients.

## Discussion

The CCDC39/40 heterodimer appears by cryo-EM as a ribbon-like structure that repeats every 96-nm along the length of the outer doublet microtubules^7,10^. We propose that the heterodimer guides multiple structural complexes to their correct positions by providing addresses for their docking. *CCDC39/40* anchors an extensive connectome of over 90 proteins that is much greater than suggested by earlier studies^33,34,40,42^. The severe lung disease in *CCDC39/40* patients may be explained by the large burden of the undocked connectome. This proteotoxic load is postulated to cause alterations in the multiciliated cell transcriptional and differentiation programs. In addition to the intracellular effects, we suggest that the microtubular disorganization results in a change in the periciliary barrier from signals in the extracellular environment.

### Ciliary address recognition proteins (CARP) identify addresses on CCDC39/40 for docking major ciliary complexes

Cilia of the *CCDC39* variant were missing 50 proteins while 40 other proteins were significantly decreased. Notably, 14 of the missing proteins directly contact CCD39/40 and are likely to be the ciliary address recognition proteins (CARPs) for IDA, the N-DRC, and the RSs. CARPs attach to specific amino acid sequences of CCDC39/40 (the addresses) (**Figure 2D, 2E**)^7–12,14,45–55^. During assembly, CARPs enter cilia coincident with CCDC39/40. The IDA and RS are prefabricated in the cytoplasm into megadalton complexes but do not include the CARPs^61,81–83^. The entry and docking of these structures follow the CARPs (**Figure 5C-E**). Therefore, the loss of CCDC39/40, and thus the addresses for CARPs in the variants result in the failure of docking of entire complexes (**Figure 3B**, **Figure 5H**). Our findings are relevant as variants in several CARPs are diseases-causing. *CFAP57*, *DRC1*, *DRC2* and *DRC4* variants result in PCD associated with the loss of the respective IDA or N-DRC complex^47,84–86^. Variants in *CCDC146* and *ZYMND12* result in human male sterility. We do not know the extent of protein loss in these variants, or their impact on multiciliated cell function.

### CCDC39/40-dependent ciliary proteins with unknown localization are missing or reduced

There are approximately 32 proteins with unknown roles that are missing or reduced in cilia from the *CCDC39* variant; most of these are transcribed primarily in multiciliated tissues or testes (**Figure 2C**). VCP and UBXN11 are reduced; UBXN11 is reported to interact with the proteosome quality control AAA-ATPase, VCP^87^. A related protein, UBXN10 along with VCP, controls primary cilia length. The reduction of UBXN11 may contribute to the short cilia of the *CCDC39/40* variants^87^. Other proteins that are absent may belong to RS3, as current structural maps are incomplete. In future studies it will be important to understand the interaction of these uncharacterized proteins with CCDC39/40 using high resolution cryo-EM and proteomics.

### *CCDC39/40* variants have an impaired periciliary barrier

Two striking features of the *CCDC39/40* variant cilia is their short length and ciliary splaying via the loss of N-DRC (**Figure 2C**)^8,41,88^. Loss of N-DRC proteins also causes microtubule misalignment in PCD patients with *DRC4* variants, but splaying was not investigated. Our SEM of the *CCDC39/40* variants in cultured cells shows that removal the ciliary membranes increased splaying substantially (**Figure 7A-C**). Interestingly, the degree of cilia splaying observed in fresh nasal biopsies of individuals with *CCDC39/40* variants was more variable (**Figure 7E**). We suggest that this significant variability in splaying occurs from inflammatory environmental insults on the ciliary membrane, which normally functions to support ciliary integrity.

The tethered mucin MUC4 was decreased in the periciliary layer (**Figure 7K**), suggesting a role of the ciliary membrane to establish or maintain the interconnected polymer structure. The relationship between ciliary integrity and a functional periciliary barrier has not been explored beforehand. The periciliary space plays an important role in separating the mucus layer and its contents from the airway epithelia^74,80^. We observed that fluorescent-labeled microbeads enter the periciliary space of variant cells but not in normal cells, which suggests that pathogens and foreign particles may more easily perturb multiciliary cells in the variants. Short cilia could also impact the function of the periciliary barrier. Disruption of the periciliary layer would further impair airway host defense and contribute to severe lung disease in these patients.

### *CCDC39/40* variants have mucous cell metaplasia *in vitro*

We were surprised to find that *in vitro* differentiation of *CCDC39/40* variant cells led to mucous cell metaplasia (**Figure 6F, 6G**). These findings are consistent with increased airway mucus plugging observed by x-ray imaging lungs of patients with *CCDC39* and *CCDC40* variants. Cultures of airway cells from individuals with COPD and cystic fibrosis show a similar phenotype that is proposed to arise from altered basal cell differentiation in response to a chronically inflamed environment^76,77,79,89^. Four lines of evidence suggest that the altered cell fate switching observed in *CCDC39/40* variants is associated with the genotype and not the environment. First, an increase in mucous cells was not observed in cultures of cells with other PCD variants that have immotile cilia and were isolated from an inflamed environment. Second, mucous cell metaplasia was observed in cells from normal individuals after depleting *CCDC39* and *CCDC40* by shRNA (**Figure 6I, 6J**). Third, an increased number of mucous cells was present in cultures of *CCDC40* knockout iPS cells derived from peripheral blood cells of a healthy donor. Fourth, transgene rescue with a normal *CCDC39* gene in the *CCDC39* variant cells reversed the mucous cell phenotype (**Figure 6K, 6L**). These lines of evidence strongly suggest that the loss of CCDC39/40 affects the shift in the cell fate of airway cells.

Single cell transcriptomics shows alterations in multiple pathways that may contribute to the mucous cell phenotype. Oxidant stress is reported to increase the number of mucous cells and expression of MUC5AC in airway epithelial cells and associated with intracellular ER stress or oxidants from neutrophilic inflammation, and cigarette smoke^71,72,74^. SAA1, SAA2 and SAA4 are upregulated in the variant multiciliated cells (**Figure 6B**). These secreted proteins are known to respond to injury, and activate inflammatory pathways in neighboring cells that may provide Notch signaling to multiciliated cells^65^. The observation that DAPT treatment decreased mucous cell number in the variant suggests inhibition of Notch signaling as an adjunctive therapy for patients with CCDC39/40 variants^66^.

Increased transcription of proteosome subunits, heat shock chaperones, and E3 ligases is consistent with increased proteostatic stress^63^. We estimate that proteins from the 6 inner dynein arms, on each ∼6 μm long cilium, from an average of 200 cilia per cell will result in one billion kilodaltons of protein that must be continually degraded by each cell. We propose that the unbound connectome proteins are responsible for generating the proteotoxic stress. The dramatic re-localization of the proteasome protein PMSB6 to the apical region of the multiciliated cell is concurrent with accumulation of unbound connectome proteins at what we term the “assembly staging zone” (**Figures 5H, 6C**). This similar localization of the proteosome and unbound connectome may indicate a demand for local quality control. Proteostasis requires significant ATP, which is consistent with the increased transcription of mitochondrial genes (**Figure S3C**). *CCDC39/40* variants share transcriptional pathways with neurodegenerative diseases (Alzheimer’s, Parkinson’s, and Huntington Disease), which have increased protein burden resulting in proteotoxicity and ATP demand^63,90^ (**Figure S3E**). Of note, neurodegenerative disease and a growing number of other diseases are being treated using inhibitors of ubiquitin and proteosome pathway modulators^91^.

### Limitations of the study

We report an extensive disruption the ciliary proteome from the *CCDCC39* variant, but we have not examined the proteome from the *CCDC40* variant. However, there is interdependence between CCDC39 and CCDC40 proteins as described in *Chlamydomonas*^22^, and we validated a similar loss of selected proteins in both *CCDC39* and *CCDC40* variant cells (e.g., **Figure 3, Figure S3**). Second, our proposal is that CCDC39/40 provides amino acid sequences as addresses for attachment of CARPs to CCDC39/40 as supported by multiple cryo-EM models and experimental data (**Figure 2D**). The mutagenesis of these addresses will be the focus of future studies. Finally, we do not know if other PCD variants will lead to similar proteostatic stress or ciliary barrier impairment by the disruption of ciliary membranes due to airway inflammation.

## Supporting information

Brody et al. scRNAseq cil3 cluster

## Acknowledgements

Transmission EM was performed with the assistance of Dr. Wandy Beatty in the Department of Molecular Microbiology at Washington University and scanning EM by Dr. Sanja Sviben in the Washington University Center for Cellular Imaging (WUCCI), which is supported by Washington University School of Medicine, The Children’s Discovery Institute of Washington University and St. Louis Children’s Hospital (CDI-CORE-2015-505 and CDI-CORE-2019-813), and the Foundation for Barnes-Jewish Hospital (3770). Proteomics was performed with the assistance of Dr. Shin-Cheng Tzeng in the Donald Danforth Plant Science Center Proteomic and Mass Spectrometry Core. We thank Greg Longmore (Washington University) for the UbiC-YFP lentivirus transfer plasmid and Mehmet Kesimer (University of North Carolina, Chapel Hill) for the anti-MUC4 antibody.

## Contributions

Conceptualization: S.L.B., S.K.D. and M.R.M.

Methodology: S.L.B, S.K.D., M.R.M., and R.Z.

Formal Analysis: J.P., J.K., and A.B.

Investigation: J.P., T.H. J.X, H.X., R.N., S.K.B, A.B., F.J.H, R.Z., X.W., M.R.M., A.H., and S.K.D.

Writing, Original Draft: S.L.B. and S.K.D.

Review and Editing: M.R.M. and A.H.

Visualization: S.L.B and J.K.

Funding acquisition: S.L.B., M.R.M., and S.K.D.

Resources: T.G.S, F.J.H, and A.H.

Supervision: S.L.B. and S.K.D.

## Supplement

### Methods Resource Table

#### Extended Data Table

Table of differential gene expression in normal compared to CCDC39/CCDC40 variant ciliated cell cluster Cil3

#### Videos

#V1. Normal cells cilia motility

#V2. Variant *CCDC39* cells cilia motility

#V3. Cilia transport of microbeads on the surface of normal cells

#V4. Cilia transport of microbeads on the surface of *CCDC39* variant cells

#V5. Cilia transport of microbeads on the surface of *CCDC40* variant cells

#V6. Variant *CCDC39* cilia motility following CCDC39 transgene rescue

**Supplement Figure S1.**
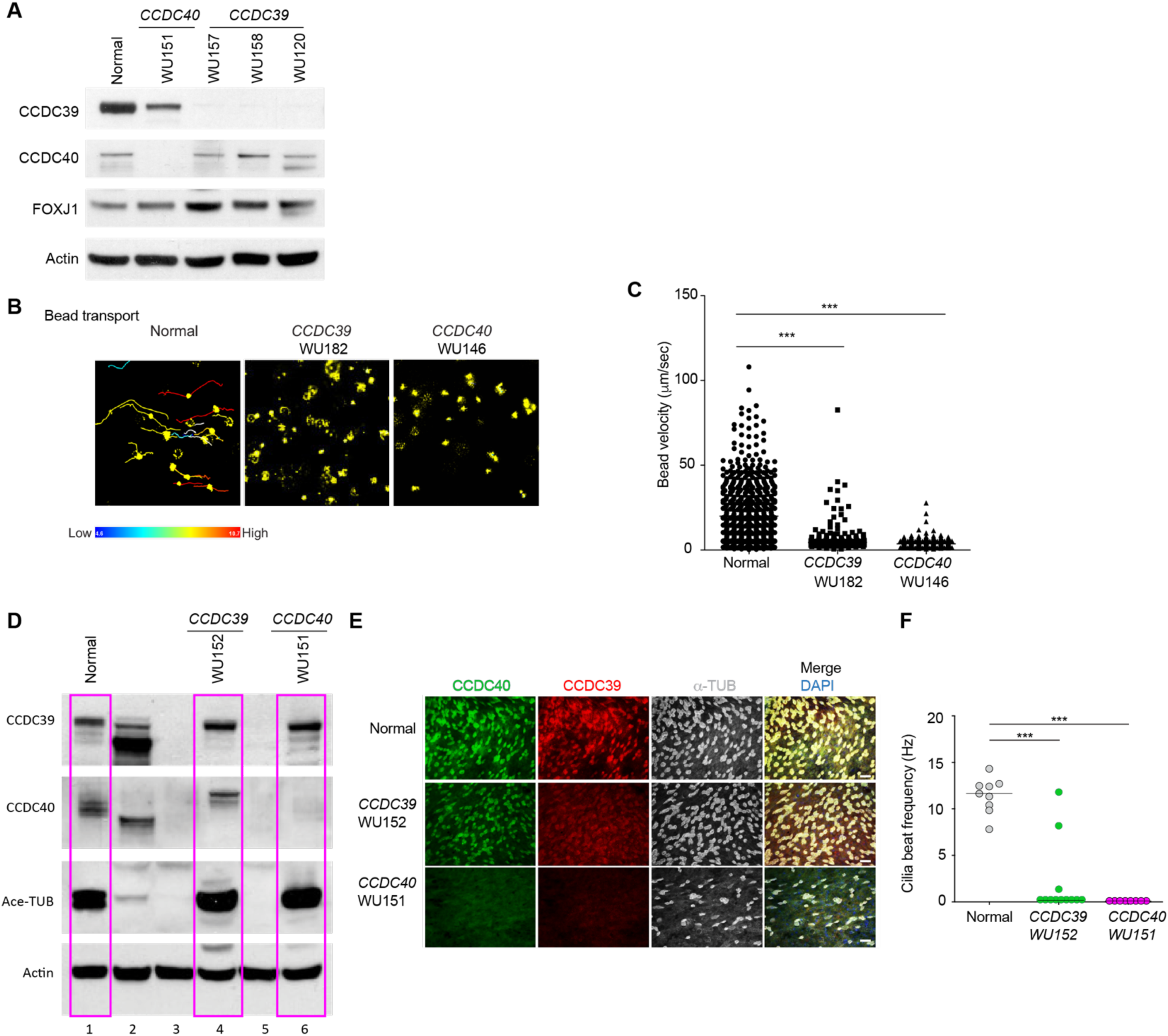
Phenotype of *CCDC39* and *CCDC40* variant cells. **(A)** Immunoblot (IB) detection of CCDC39, CCDC40, and FOXJ1 and Actin in normal and *CCDC39* and *CCDC40* variants. **(B)** Cilia transport of microbeads on the surface of well-differentiated normal, C*CDC39* (WU#182), and *CCDC40* (WU#146), from the data in Figure 1I. Shown are static images recorded over 5 sec. Trails show bead distance in normal cells. Color scale indicates relative distance. **(C)** Cilia transport determined as bead velocity. Each point represents a measure of one bead. **(D)** IB detection of CCDC39, CCDC40, and acetylated a-tubulin (Ace-TUB) and Actin in normal and *CCDC39* and *CCDC40* variants in magenta boxes (lanes 1, 4, 6). Samples in lanes 2 (normal), 3 (CCDC39 variant), 4 (*CCDC40* variant) did not differentiate into multiciliated cells and provide no information. **(E)** Immunofluorescent (IF) detection of CCDC40 and CCDC39 in the indicated *CCDC39* variant (WU152) and *CCDC40* variant (WU151). Bar=25 μm. **(F)** Cilia beat frequency in normal and *CCDC39* (WU152) and *CCDC40* (WU151) variant cells. **C**, **F**. Bar indicates median. The p value was determined by Kruskal-Wallis and with a Dunn’s Multiple Comparison Test; ***p<0.001.

**Supplement Figure S2.**
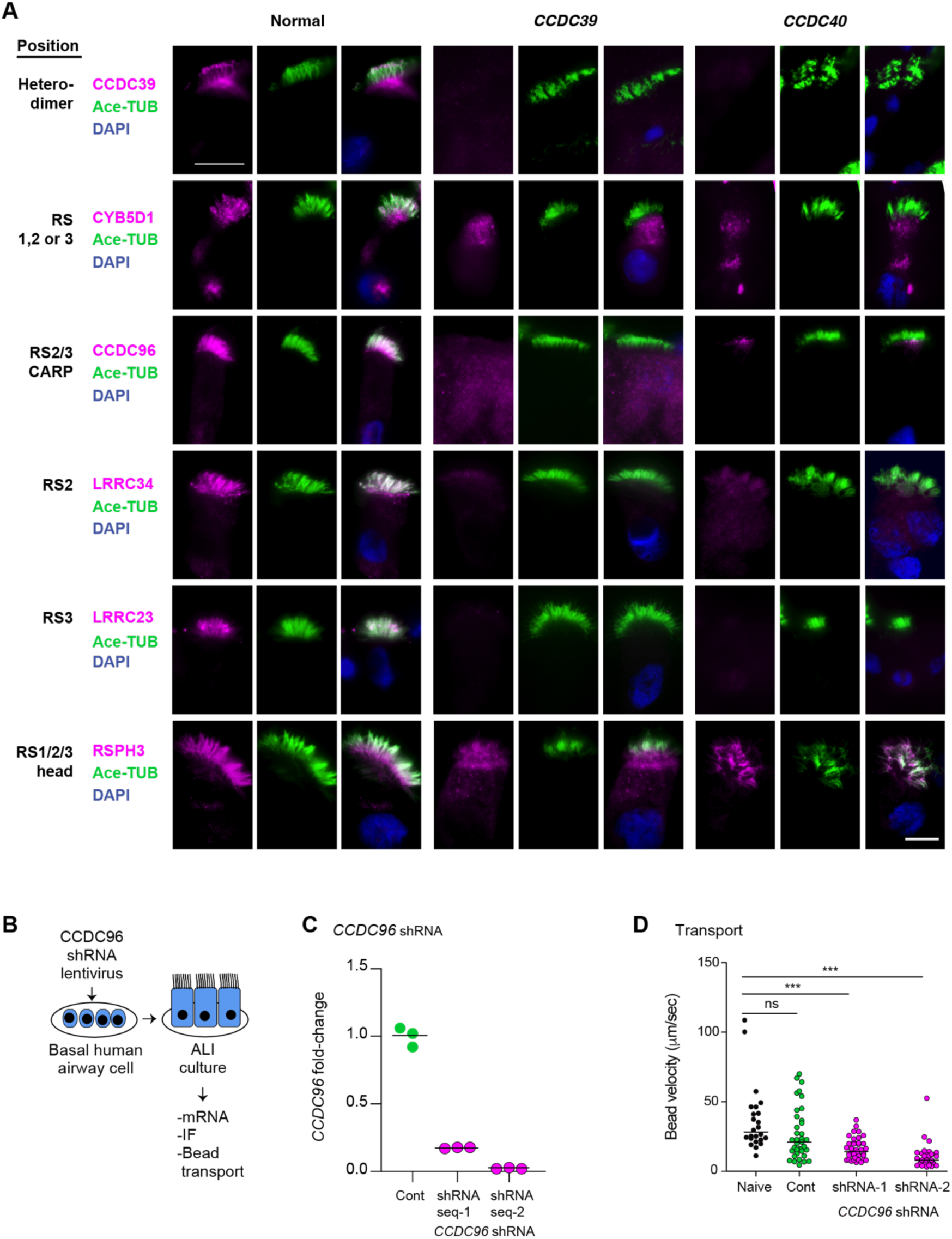
Loss of radial spokes proteins in *CCDC39/40* variants and *CCDC96* shRNA. **(A)** Immunofluorescent (IF) detection of Ace-TUB, CCDC39, and radial spoke (RS) proteins in cells from the cultures of normal and *CCDC39* and *CCDC40* variant cells. Bar=25 mm. **(B)** Scheme for lentivirus-mediated shRNA transduction of normal primary culture airway cells. **(C)** Detection of *CCDC96* mRNA in well differentiated primary culture cells. Each point represents a technical replicate of naïve cells, non-transduced, transduced with a Lentivirus Ubi-C-YFP control (Cont), or *CCDC96*-specific shRNA sequences. **(D)** Cilia transport of microbeads measured as velocity on the surface of cells non-transduced (naïve) or transduced with the indicated plasmid. Bar indicates median. The p value was determined by Kruskal-Wallis and with a Dunn’s Multiple Comparison Test; ***p<0.001.

**Supplement Figure S3.**
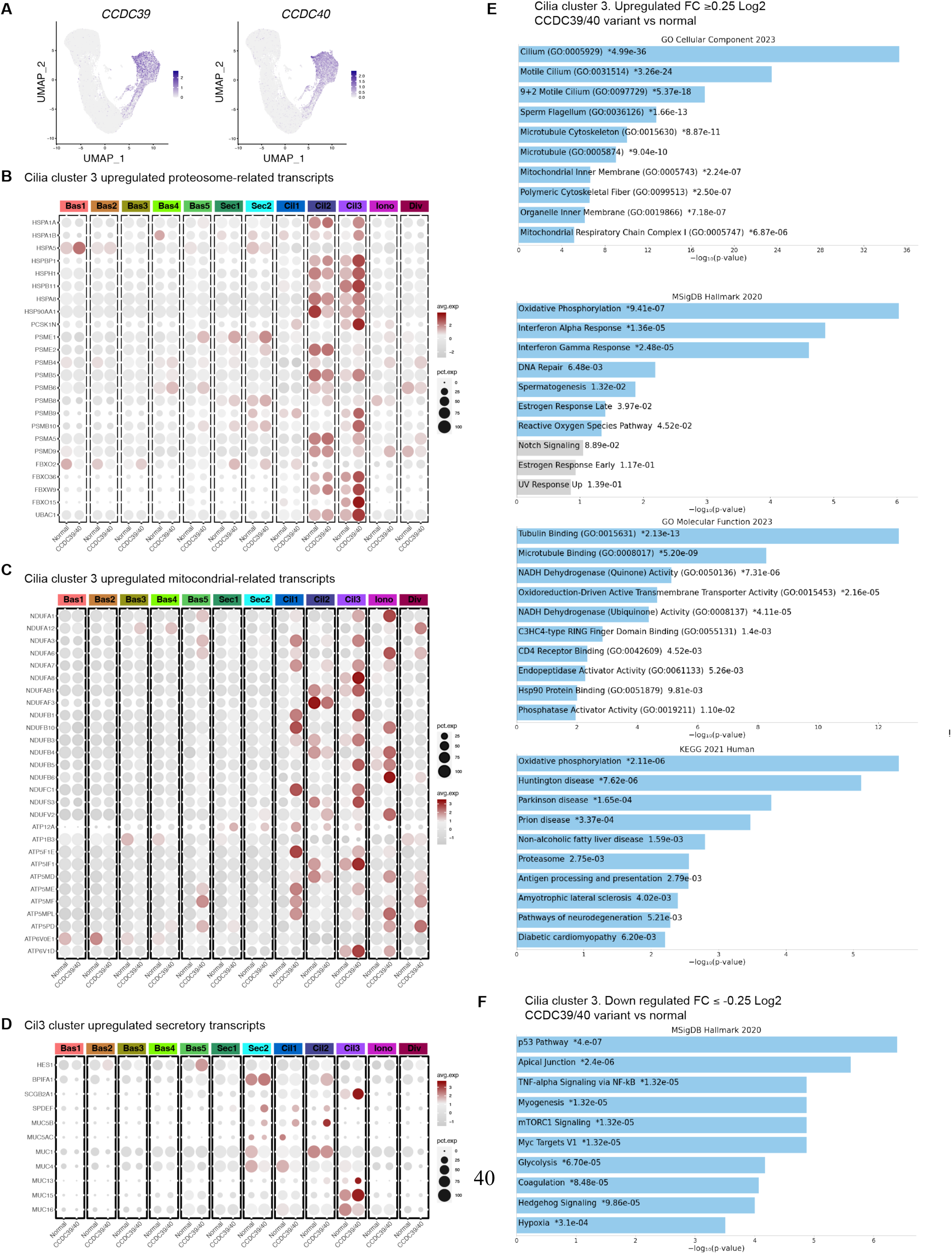
Analysis of differential expressed genes in multiciliated cells from *CCDC39*/*CCDC40* compared to normal determined using scRNA sequencing. **(A)** UMAP reductions of normal (n=5) and *CCDC39/40* variant (*CCDC39*, n=2; *CCDC40*, n=1) airway epithelial cells showing the cells that express *CCDC39* and *CCDC40*. **(B)** Dotplot representation of gene expression levels of selected mRNA related to proteostasis including heat shock proteins, subunits of the proteosome and ubiquitin E3 ligases in of *CCDC39* and *CCDC40* compared to normal cells. Clusters are shown in Figure 6A. **(C)** Dotplot representation of gene expression levels of selected mRNA of mitochondria related genes including subunits of the NADH ubiquinone oxidoreductase of mitochondrial complex and ATP synthase subunits comparing *CCDC39* and *CCDC40* to normal cells. Clusters are shown in Figure 6A. **(D)** Dotplot representation of gene expression levels of selected secretory cell genes in *CCDC39* and *CCDC40* compared to normal cells. Clusters are shown in Figure 6A. **(E)** Gene ontology and pathway analyses of upregulated transcripts from *CCDC39* and *CCDC40 cells* compared to normal cells. All genes with fold change ≥0.25 log2 were analyzed. **(F)** Pathway analysis of downregulated transcripts from *CCDC39* and *CCDC40 cells* compared to normal cells. All genes with fold change ≤-0.25 log2 were analyzed.

**Supplement Figure S4.**
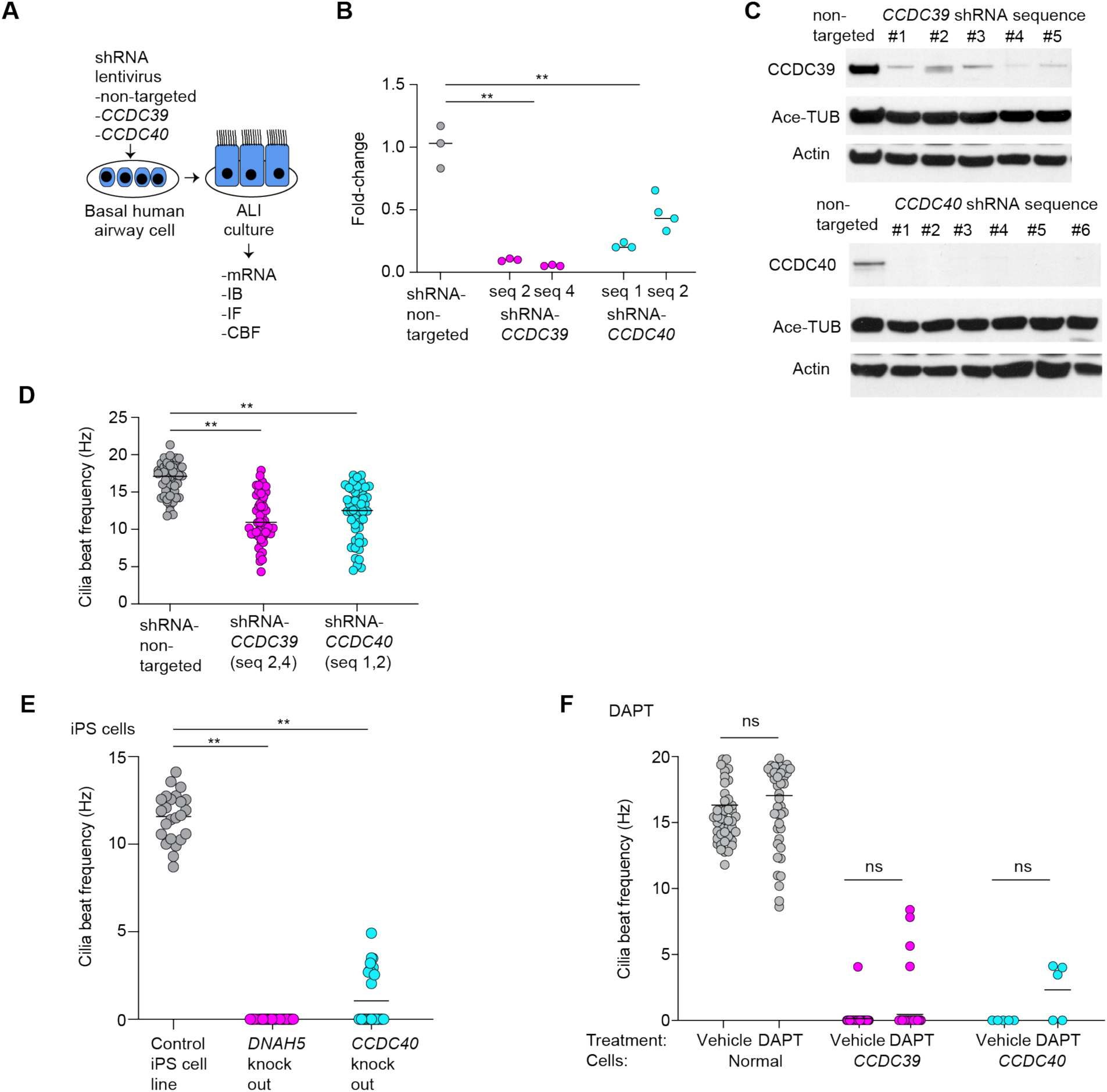
Gene expression and functions in cells with loss of *CCDC39* and *CCDC40* variant expression. **(A)** Scheme for lentivirus mediated *CCDC39* and *CCDC40*-specific shRNA transduction of normal primary culture airway cells with assessment by mRNA, immunoblot (IB), immunofluorescence (IF) and cilia beat frequency (CBF). **(B)** Quantitative reverse transcription PCR to detect changes in *CCDC39* and *CCDC40* in normal airway cells transduced with non-targeted shRNA, *CCDC39*, or *CCDC40*-specific shRNA. Shown are the median of technical replicates of two different shRNA sequences specific for *CCDC39* and *CCDC40*. **(C)** Immunoblot (IB) detection of *CCDC39* and *CCDC40* variant cells transduced with non-targeted shRNA, *CCDC39*, or *CCDC40*-specific shRNAs. **(D)** Cilia beat frequency in cells transduced with the indicated shRNAs sequences. **(E)** Cilia beat frequency in inducible pluripotent stem cells (iPS cells) from the original control line, normal and *DNAH5* knockout cells and *CCDC40* knock out cells differentiated at air-liquid interface. **(F)** Cilia beat frequency in normal, *CCDC39*, and *CCDC40* variant cells treated with vehicle and DAPT. **B**, **E**, **F** Bar indicates median. The p value was determined by: **B**, **E**, Kruskal-Wallis with a Dunn’s Multiple Comparison Test; (B, measures from both sequences for each gene were pooled); **F**, determined by the Mann-Whitney test; ns, non-significant, **p<0.01.

**Supplement Figure S5.**
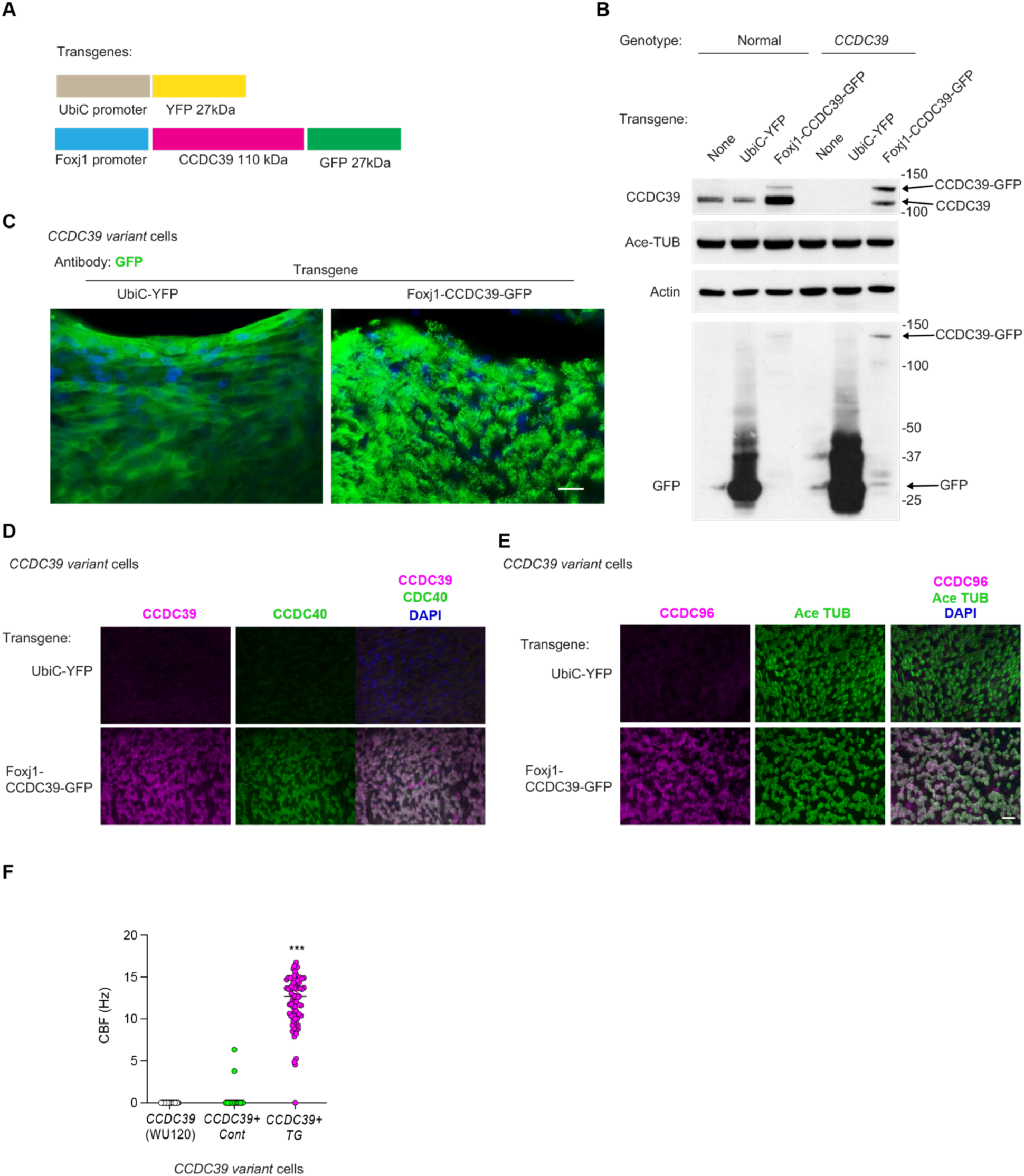
*CCDC39* transgene rescue of the *CCDC39* variant cells. **(A)** Diagram of transgenes in lentivirus vectors. **(B)** Immunoblot detection of transgenes in normal and *CCDC39* variant cells (WU182). Transduction with the *FOXJ1-CCDC39-GFP* plasmid resulted in the delivery of CCDC39 as well as CCDC39-GFP. **(C)** Immunofluorescent (IF) detection of YFP and GFP in *CCDC39* variant cells (WU182) transduced with a lentivirus; cells were differentiated at ALI day 35. Transduction with UbiC-YFP shows fluorescent signal in the cytoplasm compared to *Foxj1-CCDC39-GFP* demonstrating signal in the cilia. **(D)** IF detection of CCDC39 and CCDC40 in *CCDC39* variant cells (WU182) transduced with the *Foxj1-CCDC39-GFP* transgene. **(E)** IF detection of the ciliary address recognition protein (CARP) CCDC96 in the cilia identified by Ace-TUB after transduction of *CCDC39* variant cells (WU182) with the *Foxj1-CCDC39-GFP* transgene. **(F)** Quantitation of cilia beat frequency (CBF) in *CCDC39* variant cells (WU120), and cells transduced with control lentivirus, low or high concentration of the *CCDC39* TG. **C**, **D**, **E**. Bar=25 μm.

**Supplement Table S1.** Primary human airway cells.

| Lab number | Variant gene | Genotype | Age (yrs) | Sex | Race/Ethnicity |
| --- | --- | --- | --- | --- | --- |
| WU-103 | *DNAI1* | c.48+dupT homozygous | 15 | F | White |
| WU-105 | *DNAAF5* | c.2384T>C p.Leu795Pro homozygous | 15 | F | White |
| WU-108 | *DNAAF5* | c.2353-2354 | 3 | M | White |
| WU-111 | *DNAH5* | c.12617 G>A | 5 | F | White |
| WU-120 | *CCDC39* | c.1167+1261A>G, 3q26.33, int9 | 20 | F | Black-African American |
| WU-138 | *DNAH5* | c.12500-14A>G (Intronic)<br>/c.11658_11668del<br>(p.Leu3887Alafs*14) |  |  | Black-White |
| WU-140 | *CCDC40* | c.248DEL P.ALA83VAL FS*84/<br>C.961 C>T P.ARG321* | 15 | M | White |
| WU-145 | *CCDC40* | c.961C>T P.ARG321*;<br>C.3129DELC<br>P.PHE1044SerFS*35 | 18 | M | White |
| WU-146 | *CCDC40* | c.961C>T P.ARG321*;<br>C.3129DELC<br>P.PHE1044SerFS*35 | 15 | M | White |
| WU-150 | *DNAAF5* | c.2384T>C p.Leu795Pro | 0.7 | F | White |
| WU-151 | *CCDC40* | c.248delC(p.Ala83Valfs*84),<br>Int16 splice acceptor (c.2712-1G>T) | 40 | M | White |
| WU-152 | *CCDC39* | c.2601_2607del<br>(p.Pro868LeuFs*48) |  | F | Black-Somali |
| WU-157 | *CCDC39* | NM_181426.1:c.1871_1872del;<br>p.Ile624Lysfs*3 homo | 5-10 | M | Middle Eastern |
| WU-158 | *CCDC39* | NM_181426.1:c.1871_1872del;<br>p.Ile624Lysfs*3 homo | 5-10 | M | Middle Eastern |
| WU-165 | *DNAH5* | c.5281C>T, p.Arg1761*.;<br>c.8167C>T, p.Gln2723* | 16 | M | White |
| WU-182 | *CCDC39* | CCDC39 c.830_831delCA<br>(p.Thr277Argfs*3) heterozygous<br>Pathogenic; CCDC39<br>c.1871_1872delTA<br>(p.Ile624Lysfs*3) heterozygous<br>pathogenic, DNAH11<br>c.2966G>A (p.Arg989Gln)<br>heterozygous Uncertain<br>Significance | 57 | M | White |
| Non-disease donors for<br>scRNA seq, nasal<br>epithelial cells (NEC) |  |  |  |  |  |
| HNEC102 | None<br>detected | Normal donor | 30s | F | White |
| HNEC103 | None<br>detected | Normal donor | 40s | M | White, Middle Eastern |
| HNEC104 | None<br>detected | Normal donor | 60s | M | White |
| HNEC105 | None<br>detected | Normal donor | 60s | M | White |
| HNEC106 | None detected | Normal donor | 20s | M | White |
| HTBEC298 | Not tested | Normal donor | 25 | M | Black, non-Hispanic |
| Excess surgical tissue tracheobronchial (HTBEC) |  |  |  |  |  |
| H#185 |  | Normal donor | 59 | F | White |
| H#256 |  | Normal donor | 21 | M | Black |
| H#203 |  | Normal donor | 19 | M | Black |
| H#255 |  | Normal donor | 34 | M | White |
| H#291 |  | Normal donor | 52 | M | White |
| H#174 |  | Normal donor | 18 | F | White |
| H#299 |  | Normal donor | 36 | F | White |
| H#254 |  | Normal donor | 36 | M | Hispanic (race not available) |
| H#278 |  | Normal donor | 36 | M | White |
| H#310 |  | Normal donor | 52 | M | White |
| H#312 |  | Normal donor | 34 | F | White |
| H#243 |  | Normal donor | 43 | F | Black |
| H#314 |  | Normal donor | 40 | M | White |
| H#243 |  | Normal donor | 43 | F | Black |
| H#295 |  | Normal donor | 31 | M | White |
| H#189 |  | Normal donor | 25 | F | White |
| H#203 |  | Normal donor | 19 | M | Black |
| H#316 |  | Normal donor | 57 | F | White |
| H#276 |  | Normal donor | 35 | M | White |
| H#320 |  | Normal donor | 35 | M | White |
| H#313 |  | Normal donor | 37 | F | White |
| H#104 |  | Normal donor | 38 | F | Native American |
| H#116 |  | Normal donor | 27 | M | White |
| H#284 |  | Normal donor | 43 | M | White |

**Supplement Table S2.** Proteomic analysis of cilia from normal vs. *CCDC39* variant cells.

| **Protein** | **Normal**<br>**(Average peptide numbers)** | ***CCDC39* variant**<br>**(Average normalized**<br>**peptide numbers)** |
| --- | --- | --- |
| **CCDC39/40 proteins** |  |  |
| CCDC39 | 55.00 | 0.00 |
| CCDC40 | 62.00 | 0.00 |
| **Inner dynein arm proteins** |  |  |
| DNAH1 | 182.00 | 0.00 |
| DNAH2 | 170.00 | 0.00 |
| DNAH3 | 110.00 | 0.00 |
| DNAH6 | 195.00 | 0.00 |
| DNAH7 | 230.00 | 0.00 |
| DNAH10 | 219.00 | 0.00 |
| DNAH12 | 134.00 | 0.00 |
| CASC/P97 | 32.00 | 0.00 |
| DNALI1 | 52.00 | 6.00 |
| DNAI3 | 36.00 | 0.00 |
| DNAI4 | 41.00 | 0.00 |
| CFAP57 | 95.00 | 5.00 |
| CCDC146 | 18.00 | 0.00 |
| LRRC74B | 15.00 | 4.20 |
| ZYMND12 | 24.00 | 0.00 |
| TTC29 | 15.00 | 0.00 |
| CFAP44 | 70.00 | 0.00 |
| CFAP43 | 57.00 | 0.00 |
| CFAP58 | 74.00 | 28.00 |
| **Nexin-dynein regulatory**<br>**complex proteins** |  |  |
| DRC1 | 38.00 | 0.00 |
| DRC2/CCDC65 | 32.00 | 0.00 |
| DRC3 | 24.00 | 0.00 |
| DRC4/GAS8 | 46.00 | 0.00 |
| DRC5 | 13.00 | 0.00 |
| DRC7 | 43.00 | 0.00 |
| DRC8 | 9.00 | 0.00 |
| IQCG/DRC9 | 22.00 | 0.00 |
| IQCD/DRC10 | 34.00 | 0.00 |
| IQCA1/DRC11 | 40.00 | 0.00 |
| CFAP100 | 18.00 | 0.00 |
| CFAP73 | 20.00 | 0.00 |
| **Radial spoke proteins** |  |  |
| RSPH1 | 32.00 | 21.70 |
| DYDC2 | 15.50 | 8.80 |
| RSPH4A | 54.50 | 35.70 |
| RSPH10B | 20.50 | 4.20 |
| DNAJB13 | 37.50 | 31.50 |
| NME5 | 37.50 | 36.00 |
| RSPH3 | 60.50 | 53.90 |
| PPIL6 | 18.00 | 11.90 |
| CYB5D1 | 10.00 | 0.00 |
| RSPH9 | 54.00 | 26.00 |
| SPA17 | 18.00 | 16.80 |
| ROPN1L | 33.00 | 19.00 |
| RSPH14 | 35.00 | 19.00 |
| LRRC34 | 18.50 | 6.30 |
| LRRC23 | 34.00 | 9.90 |
| CALM1 | 11.00 | 4.20 |
| DNALI1 | 52.00 | 6.00 |
| CFAP61 | 40.00 | 0.00 |
| AK8 | 52.00 | 43.40 |
| LRRC34 | 18.50 | 6.30 |
| LRRC23 | 34.00 | 9.90 |
| IQUB | 34.00 | 11.00 |
| CFAP91 | 37.00 | 0.00 |
| CFAP251 | 49.00 | 0.00 |
| CCDC96 | 24.00 | 0.00 |
| CCDC113 | 21.00 | 0.00 |
| **Central apparatus proteins** |  |  |
| Hydin | 152.50 | 150.50 |
| SPEF2 | 69.00 | 51.63 |
| SPAG6 | 46.00 | 47.25 |
| CFAP47 | 92.00 | 54.25 |
| CFAP221 | 14.00 | 13.13 |
| CFAP69 | 31.00 | 14.88 |
| CFAP74 | 25.00 | 16.63 |
| CFAP54 | 48.50 | 29.75 |
| CFAP46 | 39.50 | 25.38 |
| LRGUK | 19.50 | 11.38 |
| CFAP74 | 25.00 | 16.60 |
| CCDC180 | 29.00 | 18.38 |
| DLEC1 | 26.00 | 16.60 |
| CFAP99 | 15.50 | 12.30 |
| MYCBP | 16.00 | 18.38 |
| SAXO2 | 85.00 | 123.38 |
| RSPH10B2 | 20.50 | 5.25 |
| WDR93 | 6.00 | 0.00 |
| SPATA17 | 15.00 | 12.25 |
| CFAP65 | 55.00 | 43.75 |
| CFAP70 | 46.00 | 42.00 |
| FAP20 | 32.50 | 34.13 |
| CFAP52 | 66.00 | 107.60 |
| **Microtubule inner proteins** |  |  |
| CFAP20 | 32.50 | 33.15 |
| CFAP21 | 60.50 | 11.90 |
| CFAP45 | 55.00 | 71.00 |
| CFAP52 | 66.00 | 87.50 |
| CFAP53 | 34.00 | 33.00 |
| NME7 | 41.50 | 51.80 |
| CFAP68 | 8.00 | 11.90 |
| CFAP77 | 52.50 | 56.75 |
| CFAP90 | 8.00 | 0.00 |
| CFAP95 | 14.00 | 23.80 |
| CFAP106/ENKUR | 33.50 | 45.00 |
| CFAP107 | 19.00 | 13.60 |
| CFAP126 | 14.50 | 36.12 |
| MNS | 47.50 | 52.70 |
| CFAP141 | 7.00 | 0.00 |
| SPAG8 | 23.50 | 12.75 |
| FAM183A | 12.00 | 22.10 |
| CFAP161 | 24.00 | 23.80 |
| PIERCE1 | 18.00 | 14.45 |
| CFAP210/CCDC173 | 45.00 | 45.00 |
| CFAP276 | 22.00 | 30.60 |
| EFHC1 | 77.50 | 86.70 |
| EFHC2 | 85.50 | 82.45 |
| RIBC2 | 36.00 | 33.15 |
| PACRG | 29.50 | 36.55 |
| TEKTIN1 | 55.50 | 51.00 |
| TEKTIN2 | 57.00 | 36.50 |
| TEKTIN3 | 33.00 | 0.00 |
| TEKTIN4 | 51.50 | 24.65 |
| FAM166B | 33.50 | 34.00 |
| SPACA9 | 36.00 | 46.75 |
| **Unknown ciliary proteins** |  |  |
| AK9 | 82.00 | 0.00 |
| ANKEF1 | 45.00 | 0.00 |
| UBXN11 | 31.00 | 0.00 |
| MDH1B | 32.50 | 0.00 |
| ICK | 17.00 | 0.00 |
| LRRC18 | 16.00 | 0.00 |
| PRKACA | 15.00 | 0.00 |
| NQO1 | 12.00 | 0.00 |
| ELONGINB | 11.00 | 0.00 |
| ELONGINC | 6.00 | 0.00 |
| CFAP299 | 10.00 | 0.00 |
| CFAP157 | 35.00 | 0.00 |
| KIAA2012 | 15.50 | 0.00 |
| RGS22 | 25.00 | 0.00 |
| CATIP | 13.00 | 0.00 |
| FAM47E | 13.00 | 0.00 |
| ARMH1 | 13.00 | 0.00 |
| ENO4 | 11.00 | 0.00 |
| ANKMY1 | 11.50 | 0.00 |
| TMC4 | 11.50 | 0.00 |
| PMSA7 | 7.50 | 0.00 |
| VCP | 67.00 | 13.00 |
| MLF 1 | 77.00 | 36.00 |
| CFAP58 | 34.00 | 28.00 |
| AK7 | 51.00 | 25.20 |
| EFCAB6 | 41.50 | 24.50 |
| PRKAR2A | 32.50 | 9.10 |
| CFAP96 | 15.50 | 4.20 |
| MYCBPAP | 21.50 | 8.40 |
| PPP1R32 | 27.00 | 4.20 |
| SAMD15 | 25.00 | 3.50 |
| STYXL1 | 11.00 | 0.00 |

## Methods

### Human studies

The Institutional Review Board of Washington University in Saint Louis approved these studies (IRB# 201705095). Consent was obtained from adults. For minors, a parent or guardian provided consent, and assent was obtained from older children. Diagnostic assessment and genetic testing for PCD was performed during clinical evaluation as described^92,93^. Human nasal epithelial cells (HNEC) were obtained by brush of the inferior nasal turbinate (Medical Packaging CytoSoft Cytology brush, #CYB-1), other cells were isolated from lungs surgically removed during lung transplantation, from surgical excess of lungs donated for lung transplantation (human tracheobronchial epithelial cells, HTBEC), or those donated for research, as previously described^75,89,92^.

### Airway epithelial cell culture

Human epithelial cells were cultured on Transwell membranes (Transwell, Corning, #3460 and #3470) and differentiated at using air-liquid interface (ALI) conditions using media as previously described^92,94,95^. Human nasal epithelial cells were isolated from biopsy brushes and the basal epithelial cells were expanded in culture using dual SMAD inhibition prior to establishing ALI conditions, as described^96^. All media included penicillin (100U/mL)/streptomycin (100 μg/mL), Primocin (100 μg/ml; InVivoGen #ant-pm-1) and amphotericin (0.25 μg/mL). Additional antibiotics were added to the media during the expansion of cells obtained from patients were with active infections according to clinical susceptibility testing. Additional antibiotics included ciprofloxacin, 20 μg/mL, tobramycin 80 μg/mL and vancomycin, 100 μg/mL.

### *Chlamydomonas* strains, culture conditions and cilia isolation

Strains were backcrossed to wild-type to verify phenotypes. The CC-125, All mutants and epitope tags were verified by PCR. Cilia were detached from cell bodies by pH shock and isolated as previously described^97^. Isolated cilia were resuspended in HMDEK buffer^98^. The double mutant *ccdc39; fap253* was constructed after backcrossing the *fap253* mutant twice to remove phototaxis and clumping phenotypes in the background^99^ using the described protocol^100^.

### RNA isolation and quantitative reverse transcription PCR

mRNA was isolated from cells using the RNeasy Plus Mini Kit Qiagen, #74134) using the manufacturer’s protocol. RNA was quantified by Nano-Drop microvolume spectrophometer (Thermo Scientific). RNA (2 μg) was reverse transcribed using the Applied Bioscience High-Capacity Reverse Transcription Kit (Thermo Fisher, #4368814). mRNA levels were determined using gene-specific primers by SYBR green nucleic acid labeling (Selleckchem PCR Mastermix, #B21202) in a Lightcycler 480 (Roche, Indianapolis, IN). Fold change was calculated using the delta delta C(t) analysis method and normalized to *OAZ1* levels^101^.

### Immunofluorescent detection and microscopy

Cultured airway cells were processed *in situ* on Transwell membranes that were cut from plastic supports and each membrane was cut into multiple pieces as described^95,102,103^. For the antibodies used in these studies, cells on membranes were fixed in methanol/acetone (50%/50% volume) for 15 min at -20 °C, followed by fixation with 4% PFA for 10 min at RT. Non-specific antibody binding was blocked in buffer containing 3% bovine serum albumin (BSA) and 0.1% TritonX100 in PBS for 45-60 min at RT. Cells were wash three times with a solution of 0.1% Tween20 in phosphate buffer saline (PBS). Membranes were incubated in primary antibodies (see Key Resource Table) diluted in blocking buffer, at 4°C overnight. The membranes were then wash 3 times and incubated with species-specific, fluorescent-labeled, donkey, secondary antibodies that were diluted in PBS for 45-60 h at RT. Membranes were wash 3 times and placed on glasses slides in medium containing 4’, 6-diamidino-2-phenylindole (DAPI) to stain DNA (Fluoroshield, Sigma Aldrich, Cat # F6057), under a coverslip. For immunostaining with CCDC39 and CCDC40 antibodies, cells underwent antigen retrieval by incubating membranes in a 1.5 mL plastic, within a capped tube containing citric acid-based antigen retrieval buffer (diluted 1:100; Antigen unmaking solution, Vector, H-3300). The tube was wrapped in flexible, waxed film (Parafilm, Amcor), placed in pressure cooker (decloaking chamber, BioCare Medical Model #109053), heated for 5 min, then allowed to cool. The membranes were washed in PBS then moved to blocking buffer, followed by the immunostaining procedure. For immunostaining with the MUC4 antibody (anti-rabbit, #69, kindly provided by Mehmet Kesimer, University of North Carolina, Chapel Hill), cultured cells were fixed with 4% PFA and processed with all detergent-free solutions.

Sections of paraffin embedded formalin fixed tissues on glass slides were rehydrated and subjected to antigen retrieval. Tissues were incubated in antibody blocking solution with a solution containing 2% fish gelatin (Sigma Aldrich #7765), 5% donkey serum (Sigma-Aldrich, #9663), and 0.1% TritonX100 in PBS for an hour prior to processing using the same protocol as used for cells on membranes. Cells were imaged by wide-field fluorescent microscopy using an upright Leica 6000 microscope equipped with a cooled digital camera interfaced with imaging software (LAS X, Leica) and a Zeiss Axiophot microscope for image capture with an UltraVIEW VoX laser spinning disk confocal system acquired by Volocity software (PerkinElmer) Images were pseudo-colored and globally adjusted for brightness and contrast using ImageJ/FIJI tools or Photoshop (Adobe) software and assembled in figures using Photoshop or Illustrator (Adobe) software.

Images of expression of CCDC39 during centriole amplification were obtained on a Nikon Eclipse Ti-E inverted confocal microscope using a 100x (1.45NA) Plan Fluor oil immersion objective lens (Nikon). Digital optical sections (Z-stacks) were captured at 0.2 - 0.3 µm intervals using a Hamamatsu ORCA-Fusion Digital CMOS camera. Image volumes were deconvolved using Nikon Elements software. For improved resolution, images were captured on a Zeiss LSM880 Airyscan confocal microscope using 60x oil immersion objective. Z-stack images were obtained at 0.16 to 0.30 µm intervals and were analyzed using ImageJ2 or Volocity (Quorum Technologies Inc.) software.

### Imaging human cilia function

Cilia beat frequency was of cells on Transwell membranes was measured using a Nikon Ti inverted microscope enclosed in a temperature controlled environmental chamber held at 37 °C, using phase contrast and Hoffman modulation contrast lenses 20x (Plan Fluoro, Nikon) or 40x (NAMC3, Nikon) as described^92,104^. Images were recorded from at least 5 fields of each Transwell insert (24-well) using a high-speed video CMOS camera and processed using Sisson-Ammons Video Analysis software (Ammons Engineering).

Transport of beads on the surface of cultured airway cells was performed as previously described^104^. Fluorescent beads (Fluoresbrite, 2 mm diameter, Polysciences, #18338-5) were diluted 1:500 in PBS. The solution was added to the apical surface of the cultures using a 10 μL pipet tip. Bead movement was recorded using a 10x objective lens of a Nikon Ti inverted microscope. Images were capture at 10 frames per sec for 5 sec per image, for a total of 50 frames per image. Data were imported to ImageJ/FIJI software for analysis using Trackmate (v. 5.2.0) and the Linear Motion LAP Tracker and Track Statistics applications in ImageJ as described^104^.

### Cilia length and splaying quantification

To maintain cilia membranes, cilia were treated with PHEM buffer (PIPES 240 mM, HEPES 100 mM, EGTA 40 mM, MgCl_2_ 8 mM, pH 7) for 1 min at room temperature. To remove cilia membranes, cells were treated with PHEM buffer containing freshly added TritonX100 0.5% and PMSF for 1 mm. Cilia were removed from the surface of cultured cells by detergent or by gently scraping with a 200 μL pipet tip. The cell surface was washed with PBS to collect cilia. Cilia in solution were partially dried on glass microscope slides. Five min after the cilia settle, excess liquid was removed with lab wiping tissue (Kimwipe) then cilia are fixed by adding 4% PFA. The cilia are immunostained with antibody to acetylated α-tubulin. About 200 cilia were assayed in over 10 fields/sample using 600X. 100-200 cilia were counted in each condition from 11-21 photographed fields. The length was measured using ImageJ.

### Periciliary barrier imaging

To image the entry of particles into the periciliary barrier, the cilia of cultured multiciliated cells were live-labeled using SiR-tubulin (Cytoskeleton, #CY-SC002) or Spy555-tubulin (Cytoskeleton, #SC203). The apical surface was washed with 2% NS culture medium (2% NuSerum, Corning, #355100) twice. SiR tubulin was diluted in the medium at 1:200, 30 μL per 24-well size while pipetting the solution a few times. The cells were returned to the incubator for 60 min. 30 μL of Rhodamine-labeled polystyrene 100 nm beads (Nanocs, #PS200-RB-1) or FITC-labeled gold nanoparticles 40 nm (Nanocs, #GP40) were added. After 10 min (in the incubator), the culture medium was gently removed from the edge of the Transwell insert. Cells were fixed by adding PFA 4% to the apical and basolateral chambers for 10 min at room temperature to stop cilia beating. The surface was not washed further. The membrane was cut from the Transwell support, placed on a slide, and mounting medium with DAPI added to the membrane, followed by a coverslip. The cells were imaged with a Zeiss Axiophot microscope and a cooled EM digital camera Zeiss AxioPhot using a 63x oil immersion lens (Carl Zeiss AG). Clusters of particles of fluorescent particles were quantified within the periciliary space between the apical surface of the cell and lower two-thirds of the cilia.

### Immunoblot analysis and Immunoprecipitation

Human cells were lysed in radioimmunoprecipitation assay (RIPA) buffer (Pierce, #89900) for 30 min at 4 °C. Lysates (35-40 µg) were separated by SDS-PAGE on NuPage 4-12% gels (Bis-Tris Gel; Invitrogen, #NP0335BOX), transferred to polyvinylidene fluoride membranes and blocked in 5% (w/v) milk powder (Nestle instant non-fat dry milk) in TBS-T buffer (0.2% Tween-20 in TBS), for 1 h at room temperature. Membranes were incubated with primary antibodies overnight at 4°C, washed in TBS-T three times, 15 min each, then incubated in rabbit or mouse horseradish peroxidase-conjugated secondary antibodies (see Key Resources Table), diluted 1:5,000, for 1 h at room temperature (see Key Resources Table). The RSP16 antibody was provided by Pinfen Yang^105^. Antibodies were visualized using Pierce ECL western blotting substrate (Thermo Fisher Scientific, #32106.) Signal was detected on X-ray film (Amersham Hyperfilm MP, Cytiva, #28906845). For *Chlamydomonas*, SDS-PAGE and immunoblotting were performed as previously described^97^. Antibodies were anti-α-tubulin antibody (Sigma-Aldrich, T6199, lot number 091M4813, 1:2000 dilution). Secondary antibodies include horseradish peroxidase (HRP)-conjugated goat anti-mouse-IgG antibody (BioRad, #1721011) and HRP-conjugated goat anti-rabbit-IgG antibody, using 1:5000 dilution (Sigma-Aldrich, #A6154).

Proteins were immunoprecipitated using the CCDC40 antibody (see STAR Methods) and analyzed by immunoblot. Cells were lysed in NP-40 Buffer (50 mM Tris HCl, pH 7.4; 150 mM NaCl, 5 mM EDTA, 1% NP-40). CCDC40 antibody, 5 µg was added to 750-1000 µg of cell lysate and rotated overnight at 4°C. Antibodies were captured with magnetic beads (Pierce Classic Magnetic IP Kit, Thermo Scientific, #88804) following the manufacturer’s protocol. Proteins were eluted using the Low pH Elution Buffer and neutralized using the Neutralization Buffer from the kit. Beads were removed and the proteins were separated on SDS-PAGE gels as described for immunoblot analysis.

### Cilia isolation

For mass spectroscopy, ciliary axonemes were isolated from cultured airway cells by application of cilia buffer with minor modifications^104^. Cells cultured in 6-12 inserts of 12-well size Transwell, were washed with PBS then incubated for 1 min with 100 μL/well of deciliation buffer (20 mM Tris, 30 mM NaCl, 10 mM CaCl_2_, 1 mM EDTA, pH 7.5) with 0.1% Triton-X100 (Surfact-Amps X-100, Thermo Scientific #28314), and freshly made 1 mM dithiothreitol, 0.05 2-mercaptoethanol, and proteinase inhibitor cocktail, Sigma #P2714) for 1 min on a vigorously moving rocker with orientation of the plate changed every 15 secs at RT. The supernatant containing cilia was collected from the wells, fresh buffer was added (100 μL) and rocking was repeated at least once and up to 5 times determined by inspection of cells for cilia. All remaining steps were performed in the cold room and samples kept on wet ice. The supernatant from the wells was pooled, then collected at 1020g for 2 min at 4°C to remove debris. A sample (5-10 μL) was reserved to confirm the presence of cilia by immunostaining with acetylated α-tubulin. The remaining supernatant was transferred to fresh tubes, and a cilia pellet was collected at 12,200g for 5 min at 4°C. The cilia pellets were combined in resuspension buffer (30 mM HEPES, 25 mM NaCl_2_, 5 mM MgSO_4_, 1 mM EGTA, 0.1 mM EDTA, 0.1 mM DTT, pH 7.3) using 100-200 mL.

The tubes were rinsed with resuspension buffer and the solution pooled into a single tube. The cilia were pelleted at 12,200 g for 5 min, at 4°C and rinsed once in 400 mL of resuspension buffer. The sample was finally collected at 20,000 g for 1 min at 4°C. The remaining supernatant was carefully removed using a gel loading tip. The tube containing the pellet was snap frozen on dry ice. The presence of isolated cilia was confirmed using immunostaining with antibody to acetylated-α-tubulin.

### Proteomics and LCMS analysis of cilia proteins

For proteomic analysis, the cilia were collected in buffer (30 mM HEPES, 25 mM NaCl_2_, 5 mM MgSO_4_, 1 mM EGTA, 0.1 mM EDTA, 0.1 mM DTT, pH 7.3) and then the pellet was snap frozen on dry ice. Protein preparation and LC-MS analysis was performed as previously described^104^. MS/MS samples were normalized and analyzed using Proteome Discoverer 2.4 (Thermo Fisher Scientific). MS/MS based peptide and protein identifications and quantification results were exported a Scaffold v5.

### Analysis of cilia proteomics data

Although equal amounts of protein were used, there were not equal amounts of cilia due to shorter cilia and cytoplasmic proteins. The peptides for α-tubulin and β-tubulin were averaged for the four normal samples and independently for the three *CCDC39* variant samples. The normal samples had 1.75 times more tubulin than the variant samples. The number of peptides for each protein in the variant was multiplied by 1.75 and compared to the normal sample number. Proteins that were missing from the normal samples were excluded from consideration as they were considered contribution from the cytoplasm. These included cytokeratins and many myosin heavy chains. Numbers of peptides in normal and variants samples were compared using a chi-squared test. Values greater than 12 were considered significant.

### Lentivirus production for shRNA and gene expression

Lentiviruses based gene specific shRNA constructs were selected from RNAi Consortium shRNA Library at the Broad Institute Genetic Perturbation Platform (https://portals.broadinstitute.org/gpp/public/). Non-targeted shRNA (Millipore Sigma Cat#SHC016) or 5-6 gene specific shRNA using 5-6 lentivirus transfer plasmids per gene target (see Key Resources Table) were used. Lentivirus was produced in HEK293T (ATCC, #11268) by cotransfection of the transfer plasmid for the gene of interest, the lentiviral psPAX packaging (Addgene, #12260) and pMD2.VSVG (Addgene, #12259) envelope plasmid (Addgene plasmids were a gift of D. Trono, EPLF, Swiss Federal Institute of Technology in Lausanne) using the calcium phosphate coprecipitation method. After approximately 18 h, transfection medium was exchanged for airway cell medium for an additional 48 h. Lentivirus gene transfer vectors UbiC-YFP (a gift of G. Longmore, Washington University)^106^ were used to express YFP in all cell types or *Foxj1-CCDC39-GFP*, expressing *CCDC39* cDNA under control of the Foxj1 promoter^107^ and each containing a puromycin selection cassette. Virus was concentrated with polyethylene glycol (Lenti-X, Takara, #631232). Lentivirus was produced in airway epithelial cell expansion medium supplemented with Y27632 and retinoic acid as described^95^. Concentrated virus was diluted at different concentrations with medium and added to the apical surface of airway basal cells seeded on Transwells at 2.2 x 10^5^ cells/cm^2^ for 3 d. Cells from at least 2 unique donors were transduced with each shRNA sequence. Transduced cells were selected with puromycin (10 μg/mL), initiated just prior to ALI culture and continued for 3-5 days. Media was changed every two or three d, for three times per week. Gene expression was confirmed using qRT-PCR or immunoblot analysis.

### Induced pluripotent stem cells

The induced pluripotent stem cell line BU3NGT and method for differentiation to induced pluripotent stem cell-derived basal cells (iBCs) was previously described^78^. For gene knockout, Cas9 enzyme (IDT, #1081058) and sgRNAs (see Key Resources Table) were delivered by nucleofection to a single cell suspension of 10^5^ iBCs in Primary Cell P3 solution (Lonza, #V4XP-3032). Cells were re-cultured and the resulting clonally-derived spheroids were isolated, expanded, and screened for targeted deletion. Screening was performed by gel electrophoresis band discrimination to identify truncated amplicons; deletions were confirmed by next generation sequencing. Knockout cells were expanded and cultured using air-liquid interface conditions as described previously^78^.

### Electron microscopy

Cultured airway cells were processed for transmission EM (TEM) as previously described^92^. Briefly, cells were scraped from Transwell membranes or fixed directly on the membranes using 2% paraformaldehyde and 2% glutaraldehyde in 100 mM sodium cacodylate buffer for 1 h at room temperature, then overnight at 4°C. Following post-fixation in 1% osmium tetroxide (Ted Pella, #18459), cells were stained in 1% aqueous uranyl acetate (Electron Microscopy Sciences), then embedded in Eponate 12 resin (Ted Pella, #18005). Sections, 95 nm thick, were post-stained with uranyl acetate and lead citrate. Sections were place on grids and imaged on a JEOL 1200 EX TEM (JEOL USA). For TEM of *Chlamydomonas*, cell pellets were prepared by high-pressure freezing followed by freeze substitution as described^108,109^. Serial thin sections, of 50−60 nm were viewed in a Philips CM10 electron microscope operating at 80 kV or 100 kV. Using the microscope’s goniometer stage, the sections were tilted to an angle that permitted the microtubules to be viewed in cross section^110^.

For scanning EM (SEM), cells cultured on Transwell membranes were fixed overnight in 2% glutaraldehyde in 0.15 M cacodylate buffer, pH 7.4 at 4°C. Membranes were then transferred into 0.1% tannic acid in ultrapure water for 20 min at room temperature. Samples were then rinsed 4 times for 5 min each in ultrapure water and incubated in 0.2% aqueous uranyl acetate for 20 min. Samples were rinsed 3 times for 10 min each in ultrapure water and dehydrated in a graded ethanol series (10%, 20%, 30%, 50%, 70%, 90%, 100%) for 5 min in each step, followed by two additional rinses in 100% ethanol. Once dehydrated, samples were loaded into a critical point drier (Leica EM CPD 300, Vienna, Austria) which was set to perform 12 CO_2_ exchanges at the slowest speed. Samples were mounted on aluminum stubs with carbon adhesive tabs and coated with 5 nm of carbon and 5 nm of iridium (Leica ACE 600, Vienna, Austria). SEM images were acquired on a FIB-SEM platform (Helios 5 UX DualBeam Fisher Scientific, Brno, Czech Republic) using SEM imaging mode at 1.8 kV and 0.1 nA using TLD detector.

### Cryo EM and image reconstruction

Cryo-EM data used to calculate the structure of doublet microtubules (DMTs) from wild-type *Chlamydomonas* was reported in a previous study^10^. Cryo-EM data of DMTs from *Chlamydomonas ccdc39* mutant were collected using a 300 kV Titan Krios microscope (Thermo Fisher Scientific) equipped with a K3 direct electron detector (Gatan) and a BioQuantum K3 Imaging Filter (slit width 20 eV) at Case Western Reserve University. Images were recorded at a defocus range of -0.5 μm to -2.5 μm with a nominal magnification of 64,000x, resulting in a pixel size of 1.34 A. Each exposure was dose-fractionated into 60 movie frames with a total exposure time of 3 sec, resulting in a total dose of 34 electrons per Å^2^. The SerialEM program was used for data collection^111^.

A total of 2,805 movies of the DMTs from *ccdc39* mutant were drift corrected and dose weighted using ‘patch motion correction’ in cryoSPARC v3.3^112^. The parameters of contrast transfer function (CTF) for different regions within the micrographs were estimated using ’patch CTF estimation’ in cryoSPARC. DMTs were automatically picked using ’filament tracer’ in cryoSPARC. DMT particles were extracted from the drift-corrected micrographs using overlapping boxes with 82.5 Å step size, and then subject to two rounds of 2D classification in cryoSPARC to remove junk and off-centered particles. Next, we performed structural refinement of the good DMT particles using ’homogeneous refinement (new)’ in cryoSPARC. The particles and their associated alignment parameters were exported to FREALIGN v9.11^113^ and subject to local refinement. In the next step, improved alignment parameters for the particles set were imported back to cryoSPARC and subjected to one round of local refinement, followed by tubulin signal subtraction.

As a routine procedure to sort out the particles with 48-nm periodicity (48-nm particles) from 8-nm particles^10^, we performed 3D classification of the tubulin-subtracted particles in Relion v3.1, using a soft-edged mask covering a local region containing large globular 48-nm MIPs in the wild type (protofilaments A08-A13). However, this 3D classification strategy failed to identify a subset of 48-nm particles with defined MIP features, indicating the original 48-nm periodicity of MIPs were significantly disrupted in the *ccdc39* mutant. We then look for the presence of 16-nm periodicity. We performed 3D classification using a soft-edged mask covering the FAP52 densities (a MIP with 16-nm periodicity) and could successfully obtain a subset of 16-nm particles. We then re-imported the particle coordinates and alignment parameters back to cryoSPARC, re-extracted the particles using 512-pixel box size and performed one round of local refinement followed by ’local CTF refinement’ and another round of local refinement, resulting in a final DMT structure with 16-nm periodicity (Fig. 4A). In this structure, the weak density of 48-nm MIP densities (compared to the wild type) is expected due to incoherent averaging and rather than the loss of 48-nm MIPs.

### Single cell RNA sequencing

Airway epithelial cells cultured in 24-well Transwell inserts for approximately 5 weeks ALI were used. One day prior to harvest, the surface of cells was washed with pre-warmed medium. Warmed trypsin solution, 0.05%, was added to the apical and basal compartments and incubated at 37 °C for 5-7 min. Cells were detached by repeatedly scraping the membrane and suspending using a 200 μL pipet tip. Cells were again incubated at 37 °C for 5-7 min in trypsin, then suspended in 0.04% bovine serum albumin in PBS. Cell viability was above 80% after processing.

Single cell RNASeq was performed through the Genome Technology Access Center at Washington University in Saint Louis using the 10x Genomics platform (3’v3.1 chemistry). Following co-encapsulation by a Chromium controller (10,000 cells targeted per sample), cDNA libraries were prepared for sequencing per manufacturer protocol, including GEM-RT reaction, cDNA amplification, and cleanup with SPRI select beads. Library size and concentration was assessed using a BioAnalyzer (Agilent) and final gene expression libraries prepared with appropriate PCR cycle modifications as recommended in the 10x Genomics Chromium Single Cell 3’ Reagent Kits User Guide (v3.1 Chemistry Dual Index). Libraries were sequenced on a NovaSeq6000 S4 Flow Cell using XP workflow and a 50×10×16×150 sequencing recipe, with a median depth target of 50,000 reads/cell. Paired-end sequencing reads were processed by CellRanger (10x Genomics software, version 7.0.0) with read alignment to the GRCh38 genome and exclusion of intronic reads.

### Single cell RNA analysis

Filtered count matrices from CellRanger were input to Seurat (version 5.0) for further analysis. Quality control cutoffs for cells (minimum genes per cell and maximum percent mitochondrial genes) were set on a per-sample basis, while genes were removed if not found in at least 3 cells. Doublets for each sample were identified by DoubletFinder (expected doublet rate set to 7.5%) and removed prior to integration. Samples were normalized by *SCTransform* (v2 regularization) and integrated according to the corresponding Seurat protocol for SCTransform normalized datasets. After Leiden clustering and UMAP visualization, *FindAllMarkers* was used to identify cluster markers (log-fold change set to 0.25). We used known markers to define broad cell categories (basal, secretory, ciliated) and defined subgroups within these categories numerically. Ciliated cells were defined by expression of *Foxj1* with the subcluster highest in genes encoding dynein motor components considered mature ciliated cells.

To identify differentially expressed genes within the mature ciliated subcluster (corresponding to Cil3 in the UMAP), we used PrepSCTFindMarkers to control for differences in signaling depth followed by *FindMarkers* with log-fold change set to 0.20. For gene ontology and pathway analyses a subset of marker genes (those with log-fold change >0.25 and adjusted p value > 0.05) were input to Enrichr (mayyanlab.cloud/enrichr). Dotplot visualizations were performed with plot1cell (https://github.com/TheHumphreysLab/plot1cell).

### Statistical analysis

Analysis was performed using GraphPad Prism (Version 10). Heatmaps were generated using the group data function in GraphPad. Differences between genotypes were compared using chi square. Differences between two groups were compared using the Mann-Whitney U test. Multiple medians were compared using the Kruskal-Wallis test, followed by Dunn’s Multiple Comparison. Paired comparisons were analyzed using the Wilcoxon Signed Rank test. For cilia beat and cilia transport studies, unpaired t tests were performed to compare each condition. P <0.05 indicated a statistically significant difference.

### Key Resources Table

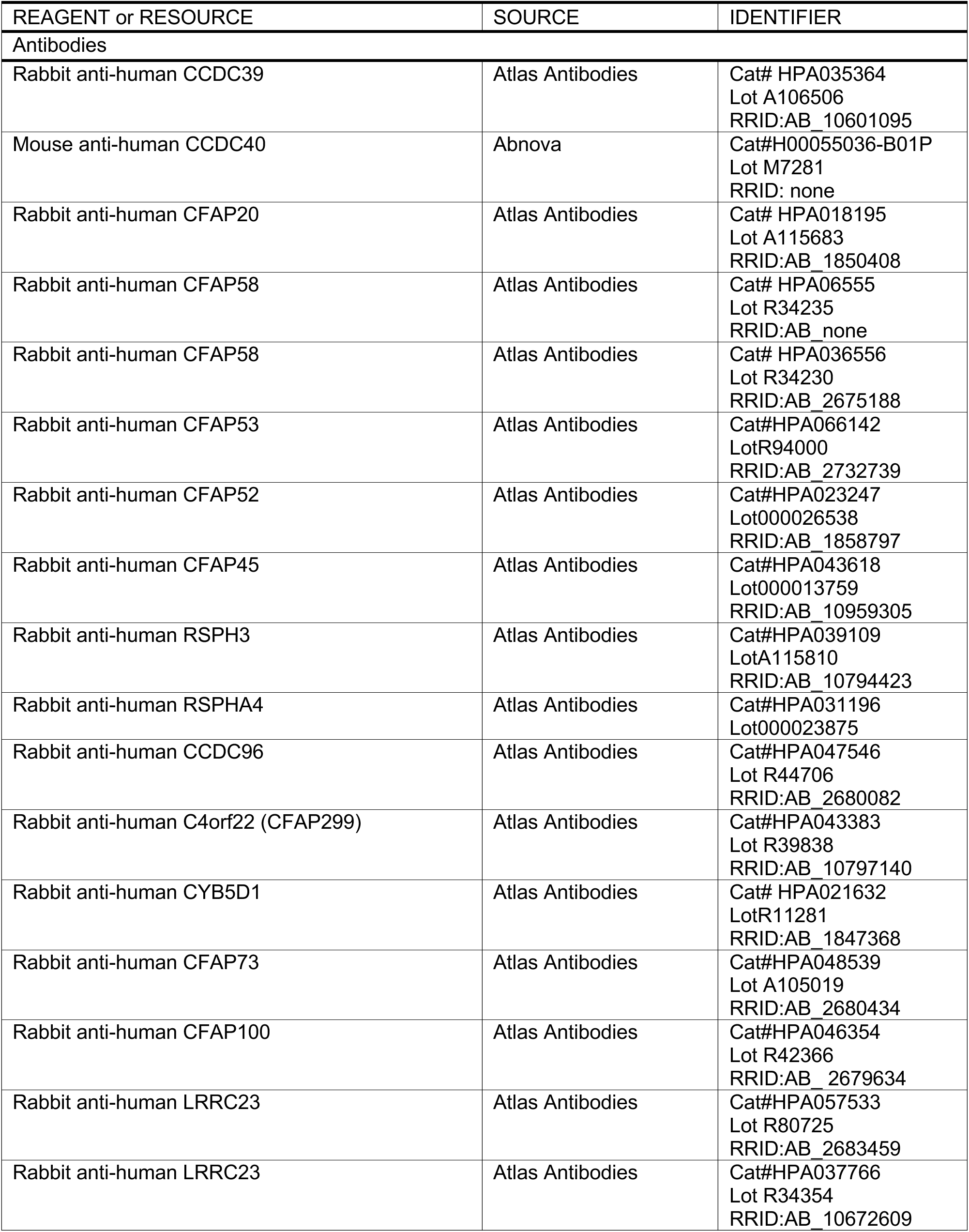
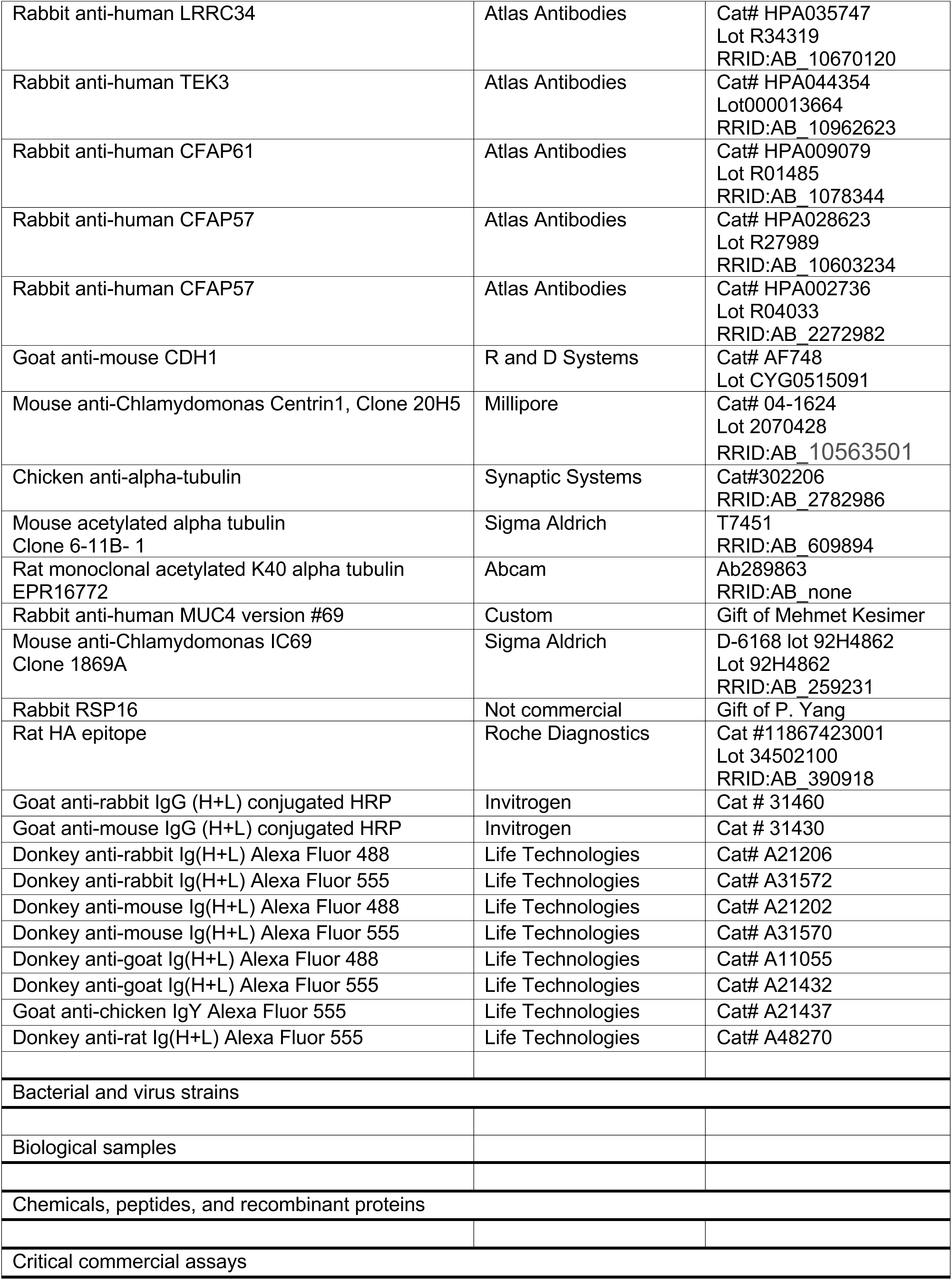
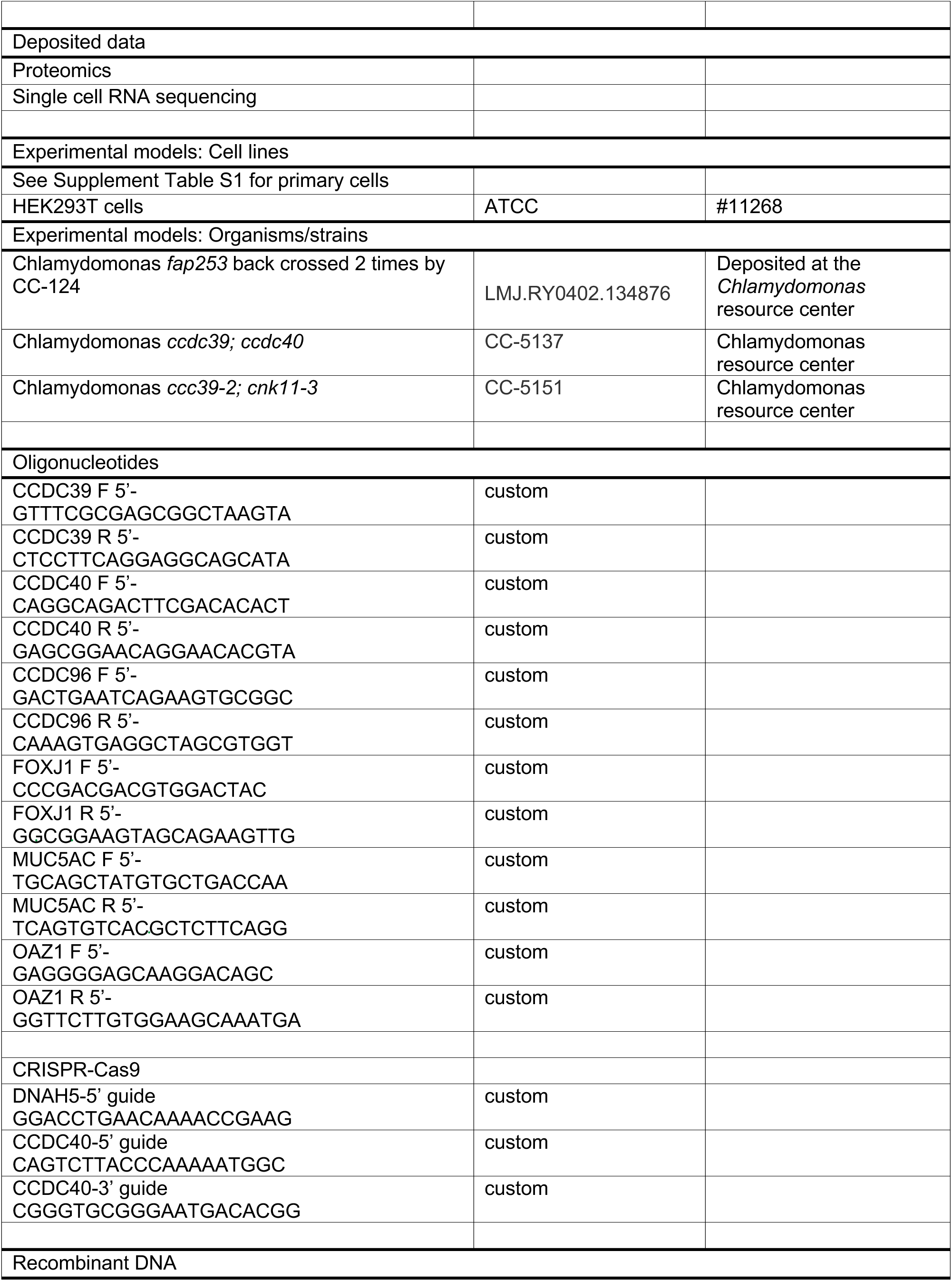
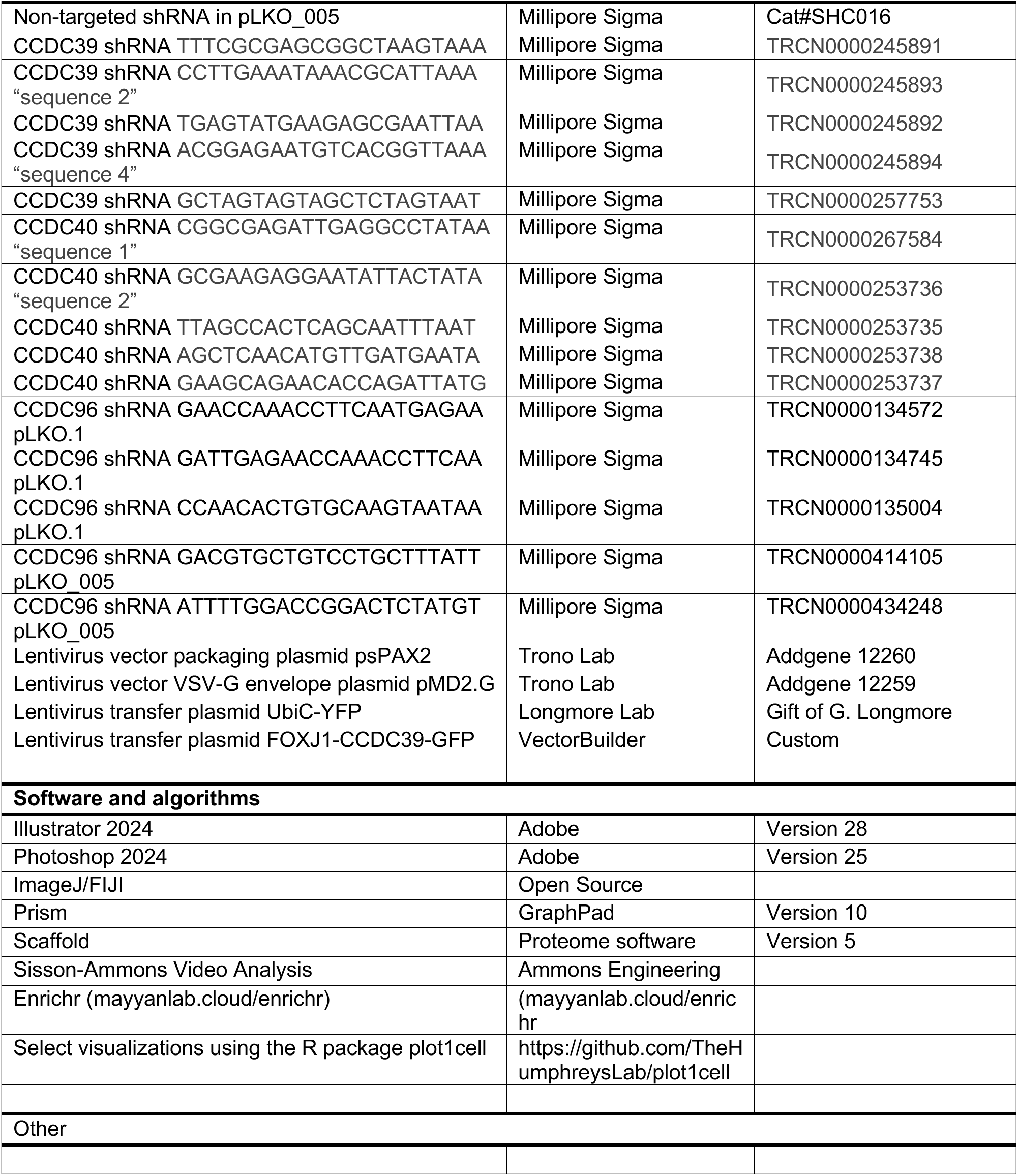

