## Supplementary material for "Loss of an extensive ciliary connectome induces proteostasis and cell fate switching in a severe motile ciliopathy": Brody et al. scRNAseq cil3 cluster

**Supplement Information. Table Single cell RNA; differentially expressed genes in the mature ciliated cell cluster (Cil3); CCDC39/40 variant vs. normal.**

| | p_val | avg_log2FC | pct.1 (CCDC39/40) | pct.2 (normal) | p_val_adj |
|---|---|---|---|---|---|
| **SAA1** | 1.33E-21 | 2.161329324 | 0.387 | 0.267 | 3.21E-17 |
| **SAA2** | 4E-19 | 1.935117333 | 0.466 | 0.365 | 9.67E-15 |
| **C15orf48** | 0.133370974 | 1.24308228 | 0.309 | 0.309 | 1 |
| **TMEM190** | 5.39E-235 | 1.23849169 | 0.997 | 0.907 | 1.3E-230 |
| **HES1** | 8.65E-104 | 1.210643034 | 0.769 | 0.536 | 2.09E-99 |
| **DYDC2** | 1.12E-254 | 1.195071333 | 0.988 | 0.91 | 2.71E-250 |
| **ROPN1L** | 5.94E-260 | 1.163169411 | 0.998 | 0.986 | 1.43E-255 |
| **RIIAD1** | 2.03E-253 | 1.157762194 | 0.979 | 0.79 | 4.89E-249 |
| **RSPH9** | 2.93E-230 | 1.100854153 | 0.987 | 0.81 | 7.07E-226 |
| **TEX26** | 6.66E-115 | 1.079734072 | 0.793 | 0.627 | 1.61E-110 |
| **SAA4** | 1.73E-48 | 1.073092669 | 0.359 | 0.173 | 4.18E-44 |
| **BPIFA1** | 1.97E-14 | 1.069928722 | 0.21 | 0.379 | 4.75E-10 |
| **LCN2** | 0.13962597 | 1.066956377 | 0.571 | 0.732 | 1 |
| **C11orf97** | 2.11E-219 | 1.058770105 | 0.981 | 0.895 | 5.11E-215 |
| **DAW1** | 9.65E-269 | 1.048150955 | 0.963 | 0.687 | 2.33E-264 |
| **C20orf85** | 2.96E-261 | 1.048019999 | 0.998 | 0.996 | 7.14E-257 |
| **FAM183A** | 0 | 1.038498464 | 0.998 | 0.986 | 0 |
| **AC013264.1** | 3.19E-203 | 1.025937593 | 0.987 | 0.834 | 7.7E-199 |
| **CCDC173** | 5.61E-239 | 1.022047504 | 0.976 | 0.736 | 1.36E-234 |
| **LRRC23** | 1.65E-241 | 1.018251931 | 1 | 0.952 | 3.99E-237 |
| **IFI27** | 1.03E-07 | 0.988526393 | 0.684 | 0.645 | 0.002484522 |
| **MNS1** | 8.24E-287 | 0.976120047 | 0.999 | 0.938 | 1.99E-282 |
| **FAM81B** | 2.9E-288 | 0.973615265 | 0.999 | 0.942 | 7E-284 |
| **SPA17** | 1.06E-247 | 0.96968982 | 0.997 | 0.951 | 2.56E-243 |
| **TCTEX1D4** | 8.58E-105 | 0.964752733 | 0.949 | 0.855 | 2.07E-100 |
| **LXN** | 1.59E-154 | 0.96425518 | 0.928 | 0.741 | 3.84E-150 |
| **TPPP3** | 0 | 0.952253801 | 1 | 0.999 | 0 |
| **MRLN** | 1.02E-202 | 0.950419627 | 0.796 | 0.353 | 2.46E-198 |
| **DNAAF4** | 2.21E-237 | 0.945244329 | 0.994 | 0.89 | 5.35E-233 |
| **AC108134.4** | 3E-170 | 0.937559098 | 0.866 | 0.539 | 7.24E-166 |
| **LAP3** | 3.78E-125 | 0.93556841 | 0.961 | 0.799 | 9.14E-121 |
| **RRAD** | 1.03E-114 | 0.935081673 | 0.978 | 0.945 | 2.49E-110 |
| **C1orf189** | 1E-119 | 0.927297733 | 0.838 | 0.561 | 2.43E-115 |
| **CFAP126** | 1.59E-241 | 0.927269037 | 0.998 | 0.954 | 3.84E-237 |

| | | | | | |
|---|---|---|---|---|---|
| **ENKUR** | 2.94E-282 | 0.918001536 | 0.997 | 0.953 | 7.11E-278 |
| **CFAP53** | 7.04E-222 | 0.908310718 | 0.996 | 0.936 | 1.7E-217 |
| **HIST1H4C** | 5.36E-153 | 0.905576912 | 0.731 | 0.373 | 1.29E-148 |
| **CAPSL** | 9.44E-237 | 0.898852385 | 0.998 | 0.976 | 2.28E-232 |
| **DNALI1** | 2.36E-253 | 0.896267431 | 1 | 0.98 | 5.71E-249 |
| **STOML3** | 8.26E-221 | 0.879704561 | 0.988 | 0.825 | 1.99E-216 |
| **CCDC65** | 6.84E-219 | 0.879482184 | 0.993 | 0.887 | 1.65E-214 |
| **AL121899.1** | 7.63E-182 | 0.874431664 | 0.83 | 0.444 | 1.84E-177 |
| **C5orf49** | 4.1E-224 | 0.874211405 | 0.996 | 0.985 | 9.9E-220 |
| **CFAP36** | 3.06E-174 | 0.873800488 | 0.996 | 0.88 | 7.4E-170 |
| **LRRC10B** | 6.97E-191 | 0.87251011 | 0.992 | 0.881 | 1.68E-186 |
| **AC105446.1** | 9.93E-129 | 0.871272539 | 0.741 | 0.394 | 2.4E-124 |
| **ECRG4** | 3.05E-85 | 0.868438592 | 0.753 | 0.473 | 7.37E-81 |
| **IDH2** | 4.66E-152 | 0.862646526 | 0.97 | 0.798 | 1.13E-147 |
| **CFAP298** | 6.68E-283 | 0.860497446 | 0.999 | 0.99 | 1.61E-278 |
| **CFAP300** | 3.44E-177 | 0.860064923 | 0.996 | 0.929 | 8.3E-173 |
| **RSPH1** | 2.35E-248 | 0.858409894 | 1 | 0.998 | 5.67E-244 |
| **OMG** | 1.24E-106 | 0.85662424 | 0.984 | 0.956 | 2.99E-102 |
| **ANKRD37** | 1.09E-154 | 0.852138235 | 0.958 | 0.776 | 2.63E-150 |
| **IQCG** | 2.88E-259 | 0.844218772 | 0.999 | 0.945 | 6.95E-255 |
| **FOXJ1** | 8.02E-176 | 0.842848296 | 0.999 | 0.95 | 1.94E-171 |
| **C11orf88** | 8.2E-120 | 0.840246315 | 0.994 | 0.949 | 1.98E-115 |
| **KIF9** | 1.04E-239 | 0.835854794 | 0.994 | 0.965 | 2.51E-235 |
| **IFI6** | 3.81E-09 | 0.835696713 | 0.362 | 0.286 | 9.2E-05 |
| **WDR54** | 2.23E-181 | 0.834778321 | 1 | 0.937 | 5.38E-177 |
| **CCDC74B** | 7.38E-147 | 0.831775913 | 0.922 | 0.639 | 1.78E-142 |
| **SCGB3A1** | 0.000414585 | 0.831685272 | 0.128 | 0.186 | 1 |
| **CALML4** | 1.06E-183 | 0.83087677 | 0.997 | 0.924 | 2.57E-179 |
| **ODF3B** | 3.14E-249 | 0.82805988 | 1 | 0.988 | 7.58E-245 |
| **PPIL6** | 5.57E-231 | 0.827502843 | 0.994 | 0.925 | 1.35E-226 |
| **CFAP299** | 1.71E-207 | 0.826200796 | 0.953 | 0.626 | 4.12E-203 |
| **CTXN1** | 5.44E-123 | 0.824573241 | 0.978 | 0.851 | 1.31E-118 |
| **GDF15** | 7.57E-33 | 0.8232529 | 0.747 | 0.69 | 1.83E-28 |
| **SMIM22** | 1.3E-240 | 0.816696353 | 0.996 | 0.985 | 3.15E-236 |
| **C9orf116** | 1.27E-208 | 0.816162455 | 0.998 | 0.989 | 3.08E-204 |
| **PCSK1N** | 2.88E-94 | 0.815446127 | 0.868 | 0.699 | 6.97E-90 |
| **PSENEN** | 7E-233 | 0.811404559 | 1 | 0.986 | 1.69E-228 |
| **C9orf135** | 3.2E-160 | 0.810254237 | 0.992 | 0.866 | 7.73E-156 |

| | | | | | |
|---|---|---|---|---|---|
| **APOO** | 6.2E-166 | 0.807696135 | 0.946 | 0.693 | 1.5E-161 |
| **CABCOCO1** | 4.86E-162 | 0.803425241 | 0.982 | 0.861 | 1.17E-157 |
| **AC007906.2** | 2.28E-202 | 0.801024292 | 1 | 0.987 | 5.51E-198 |
| **IFT57** | 1.66E-215 | 0.800677371 | 0.999 | 0.99 | 4.02E-211 |
| **IGFBP7** | 1.49E-109 | 0.796621237 | 0.994 | 0.995 | 3.6E-105 |
| **FAM229B** | 1.23E-161 | 0.793168774 | 0.998 | 0.977 | 2.97E-157 |
| **SPAG1** | 7.28E-211 | 0.79126594 | 0.998 | 0.931 | 1.76E-206 |
| **FGGY** | 6.01E-141 | 0.790850642 | 0.909 | 0.671 | 1.45E-136 |
| **SNTN** | 4.11E-242 | 0.790791525 | 1 | 0.997 | 9.93E-238 |
| **CCDC74A** | 2.76E-126 | 0.789550185 | 0.986 | 0.843 | 6.66E-122 |
| **HSPA1A** | 1.85E-101 | 0.789089939 | 0.841 | 0.599 | 4.48E-97 |
| **IFT74** | 7.19E-194 | 0.784221798 | 0.983 | 0.804 | 1.74E-189 |
| **RUVBL1** | 2.31E-190 | 0.781535413 | 0.999 | 0.939 | 5.58E-186 |
| **C17orf97** | 2.02E-187 | 0.777039242 | 0.977 | 0.783 | 4.88E-183 |
| **MORN5** | 3.47E-178 | 0.776264659 | 0.994 | 0.96 | 8.38E-174 |
| **RSPH4A** | 1.24E-178 | 0.775476272 | 0.998 | 0.967 | 2.99E-174 |
| **TCTEX1D2** | 7.51E-176 | 0.772536571 | 0.992 | 0.965 | 1.81E-171 |
| **HIST1H1C** | 2.05E-79 | 0.771874786 | 0.861 | 0.638 | 4.96E-75 |
| **C9orf24** | 8.52E-156 | 0.762960413 | 0.996 | 0.985 | 2.06E-151 |
| **ZMYND10** | 3.03E-132 | 0.760435497 | 0.999 | 0.957 | 7.32E-128 |
| **ATF7IP2** | 8.56E-87 | 0.759944127 | 0.857 | 0.646 | 2.07E-82 |
| **SLAIN2** | 1.46E-87 | 0.756502884 | 0.874 | 0.677 | 3.53E-83 |
| **MAP1A** | 3.42E-121 | 0.756205833 | 0.988 | 0.886 | 8.26E-117 |
| **TTC29** | 2.93E-181 | 0.754784619 | 0.967 | 0.768 | 7.07E-177 |
| **MORN2** | 5.42E-157 | 0.753421742 | 0.998 | 0.993 | 1.31E-152 |
| **RAMP1** | 9E-44 | 0.753235891 | 0.502 | 0.316 | 2.17E-39 |
| **SPATA17** | 3.59E-189 | 0.752025425 | 0.992 | 0.863 | 8.67E-185 |
| **CCDC146** | 2.04E-243 | 0.75188307 | 1 | 0.988 | 4.93E-239 |
| **STMND1** | 4.93E-144 | 0.749306024 | 0.934 | 0.7 | 1.19E-139 |
| **C2orf73** | 8.79E-106 | 0.748951571 | 0.882 | 0.639 | 2.12E-101 |
| **SCGB2A1** | 4.79E-31 | 0.744605944 | 0.736 | 0.623 | 1.16E-26 |
| **TUBA4B** | 9.99E-178 | 0.743883171 | 0.996 | 0.858 | 2.41E-173 |
| **IQCD** | 2.71E-154 | 0.740850126 | 0.964 | 0.772 | 6.54E-150 |
| **C1orf194** | 2.86E-171 | 0.739523461 | 0.996 | 0.99 | 6.9E-167 |
| **TEX9** | 3.07E-183 | 0.738514861 | 0.989 | 0.84 | 7.41E-179 |
| **PIH1D2** | 1.67E-176 | 0.737735522 | 0.972 | 0.767 | 4.03E-172 |
| **LINC02166** | 1.67E-146 | 0.736083961 | 0.761 | 0.384 | 4.03E-142 |
| **FAM92B** | 5.77E-141 | 0.735130898 | 0.997 | 0.972 | 1.39E-136 |

| | | | | | |
|---|---|---|---|---|---|
| **EFCAB1** | 8.43E-127 | 0.733438434 | 0.998 | 0.952 | 2.04E-122 |
| **ACBD3-AS1** | 3.69E-109 | 0.731888863 | 0.94 | 0.76 | 8.92E-105 |
| **CES1** | 5.12E-21 | 0.729223807 | 0.586 | 0.53 | 1.24E-16 |
| **ARL 3.00** | 9.19E-193 | 0.727679891 | 0.997 | 0.987 | 2.22E-188 |
| **BBOF1** | 1.05E-165 | 0.727510654 | 0.992 | 0.911 | 2.55E-161 |
| **AGR3** | 7.48E-187 | 0.726433316 | 0.999 | 0.99 | 1.81E-182 |
| **CETN2** | 1.7E-249 | 0.726117556 | 0.999 | 0.995 | 4.1E-245 |
| **SPAG6** | 4.1E-179 | 0.723016082 | 0.986 | 0.936 | 9.91E-175 |
| **LINC01765** | 1.51E-121 | 0.719375635 | 0.837 | 0.541 | 3.64E-117 |
| **EFCAB10** | 5.87E-130 | 0.717745277 | 0.966 | 0.833 | 1.42E-125 |
| **TUBA1A** | 1.42E-133 | 0.716130532 | 1 | 0.997 | 3.44E-129 |
| **CCNA1** | 3.33E-96 | 0.7125046 | 0.272 | 0.063 | 8.04E-92 |
| **GON7** | 9.03E-150 | 0.708609618 | 0.989 | 0.916 | 2.18E-145 |
| **C22orf15** | 6.34E-69 | 0.708412177 | 0.914 | 0.747 | 1.53E-64 |
| **DYNLRB2** | 2.66E-168 | 0.707789918 | 0.992 | 0.974 | 6.43E-164 |
| **AKAP14** | 1.6E-125 | 0.707788247 | 0.984 | 0.887 | 3.87E-121 |
| **NDUFB1** | 2.67E-179 | 0.706423894 | 0.992 | 0.927 | 6.44E-175 |
| **GSTA1** | 7.2E-39 | 0.70411103 | 0.906 | 0.919 | 1.74E-34 |
| **TTC12** | 5.39E-147 | 0.701581294 | 0.87 | 0.527 | 1.3E-142 |
| **ARMC4** | 2.28E-142 | 0.700142296 | 0.948 | 0.755 | 5.5E-138 |
| **METTL27** | 7.09E-135 | 0.698875477 | 0.75 | 0.359 | 1.71E-130 |
| **DPCD** | 1.51E-157 | 0.697251879 | 0.994 | 0.897 | 3.65E-153 |
| **SRGAP3-AS2** | 1.74E-67 | 0.695133029 | 0.858 | 0.737 | 4.21E-63 |
| **OXTR** | 1.68E-85 | 0.694949843 | 0.397 | 0.145 | 4.06E-81 |
| **COPRS** | 1.29E-140 | 0.694280007 | 0.987 | 0.844 | 3.11E-136 |
| **GIHCG** | 3.05E-108 | 0.693063261 | 0.961 | 0.814 | 7.37E-104 |
| **TTC25** | 5.43E-142 | 0.692731882 | 0.963 | 0.8 | 1.31E-137 |
| **AL357093.2** | 2.91E-66 | 0.688964861 | 0.996 | 0.966 | 7.02E-62 |
| **NME5** | 5.67E-153 | 0.680559891 | 0.99 | 0.954 | 1.37E-148 |
| **IFT22** | 1.07E-145 | 0.678315864 | 0.997 | 0.939 | 2.59E-141 |
| **FAIM** | 2.68E-149 | 0.675797434 | 0.92 | 0.62 | 6.47E-145 |
| **PPP1R42** | 4.62E-132 | 0.675549255 | 0.942 | 0.719 | 1.11E-127 |
| **EFCAB2** | 1.94E-147 | 0.673185973 | 0.983 | 0.785 | 4.7E-143 |
| **FANK1** | 1.76E-182 | 0.673031008 | 0.998 | 0.927 | 4.25E-178 |
| **CCL15** | 1.51E-96 | 0.672983801 | 0.847 | 0.589 | 3.66E-92 |
| **HSPBP1** | 2.52E-110 | 0.672354125 | 0.973 | 0.805 | 6.09E-106 |
| **RIBC2** | 1.59E-122 | 0.669571483 | 0.93 | 0.704 | 3.84E-118 |
| **LDLRAD1** | 1.76E-108 | 0.669084047 | 0.98 | 0.897 | 4.25E-104 |

| | | | | | |
|---|---|---|---|---|---|
| **TCTEX1D1** | 4.56E-135 | 0.668893857 | 0.966 | 0.795 | 1.1E-130 |
| **PACRG** | 4.65E-150 | 0.667843534 | 0.983 | 0.834 | 1.12E-145 |
| **MT-CO1** | 1.06E-158 | 0.667641397 | 1 | 1 | 2.56E-154 |
| **SPAG16** | 1.84E-160 | 0.662811668 | 0.999 | 0.932 | 4.44E-156 |
| **KIAA1211L** | 4.29E-140 | 0.662122042 | 0.978 | 0.801 | 1.04E-135 |
| **CCN2** | 4.24E-51 | 0.660821936 | 0.931 | 0.917 | 1.03E-46 |
| **NPHP1** | 1.65E-154 | 0.653904235 | 0.993 | 0.923 | 3.98E-150 |
| **TEKT1** | 5.34E-165 | 0.653732888 | 1 | 0.946 | 1.29E-160 |
| **ST3GAL6** | 1.3E-74 | 0.652397591 | 0.582 | 0.316 | 3.15E-70 |
| **CFAP45** | 1.33E-150 | 0.651663468 | 0.997 | 0.947 | 3.2E-146 |
| **ARMC3** | 2.04E-167 | 0.650781864 | 1 | 0.962 | 4.92E-163 |
| **OSCP1** | 1.4E-130 | 0.645339235 | 0.982 | 0.836 | 3.38E-126 |
| **CCDC78** | 3.62E-128 | 0.644519671 | 0.997 | 0.94 | 8.75E-124 |
| **STK33** | 3.12E-123 | 0.643045173 | 0.978 | 0.888 | 7.55E-119 |
| **REC8** | 1.21E-89 | 0.640630183 | 0.721 | 0.428 | 2.91E-85 |
| **FAM216B** | 1.71E-136 | 0.638746285 | 0.994 | 0.955 | 4.12E-132 |
| **C11orf16** | 1.04E-98 | 0.634799435 | 0.76 | 0.466 | 2.51E-94 |
| **PIFO** | 7.43E-154 | 0.634355959 | 1 | 0.996 | 1.79E-149 |
| **LRRC34** | 6.26E-122 | 0.633418339 | 0.961 | 0.761 | 1.51E-117 |
| **FAM166B** | 1.44E-104 | 0.630961265 | 0.883 | 0.634 | 3.48E-100 |
| **CLDN4** | 2.11E-59 | 0.628989771 | 0.959 | 0.918 | 5.1E-55 |
| **CASC1** | 1.03E-114 | 0.628836815 | 0.981 | 0.894 | 2.49E-110 |
| **WDR38** | 2.33E-94 | 0.62762323 | 0.934 | 0.787 | 5.63E-90 |
| **LRRC46** | 1.94E-127 | 0.626274042 | 0.991 | 0.929 | 4.68E-123 |
| **AK7** | 1.62E-107 | 0.622274278 | 0.984 | 0.888 | 3.91E-103 |
| **LRRIQ1** | 2.32E-186 | 0.622263209 | 1 | 0.994 | 5.6E-182 |
| **WDR78** | 3.21E-137 | 0.621782205 | 1 | 0.962 | 7.75E-133 |
| **CFAP221** | 8.04E-131 | 0.620589406 | 0.987 | 0.833 | 1.94E-126 |
| **DZIP3** | 3.5E-120 | 0.617133022 | 0.991 | 0.937 | 8.45E-116 |
| **CC2D2A** | 1.6E-115 | 0.615023187 | 0.99 | 0.918 | 3.87E-111 |
| **MRPL23** | 2.46E-127 | 0.614206001 | 0.771 | 0.402 | 5.94E-123 |
| **ZFP36** | 1.04E-49 | 0.613809108 | 0.791 | 0.647 | 2.52E-45 |
| **MSH3** | 3.44E-81 | 0.613471792 | 0.854 | 0.655 | 8.31E-77 |
| **ANKRD66** | 2.49E-114 | 0.612963145 | 0.97 | 0.855 | 6E-110 |
| **MUC12** | 9E-74 | 0.612358688 | 0.607 | 0.329 | 2.17E-69 |
| **LRRC6** | 5.4E-131 | 0.611702407 | 0.957 | 0.796 | 1.3E-126 |
| **CCDC96** | 1.95E-121 | 0.610837886 | 0.954 | 0.818 | 4.71E-117 |
| **RUVBL2** | 3.94E-134 | 0.610279481 | 0.986 | 0.866 | 9.51E-130 |

| | | | | | |
|---|---|---|---|---|---|
| **CCDC153** | 2.28E-120 | 0.608818211 | 0.988 | 0.911 | 5.51E-116 |
| **LRWD1** | 5.04E-110 | 0.608621941 | 0.877 | 0.614 | 1.22E-105 |
| **MYCBP** | 4.75E-148 | 0.608527108 | 0.998 | 0.918 | 1.15E-143 |
| **C11orf74** | 4.35E-108 | 0.607983802 | 0.976 | 0.867 | 1.05E-103 |
| **ATP5ME** | 8.55E-155 | 0.607702701 | 0.991 | 0.941 | 2.06E-150 |
| **CRIP1** | 2.99E-75 | 0.606858368 | 0.998 | 0.996 | 7.23E-71 |
| **CFAP73** | 2.23E-114 | 0.606394512 | 0.971 | 0.838 | 5.39E-110 |
| **AP001207.3** | 1.55E-107 | 0.605123579 | 0.72 | 0.396 | 3.75E-103 |
| **CENPM** | 1.41E-91 | 0.604188932 | 0.719 | 0.409 | 3.42E-87 |
| **STX2** | 1.32E-97 | 0.600962728 | 0.84 | 0.571 | 3.18E-93 |
| **WDR86-AS1** | 7.82E-29 | 0.599473618 | 0.852 | 0.818 | 1.89E-24 |
| **SAMD15** | 5.39E-68 | 0.599269535 | 0.819 | 0.583 | 1.3E-63 |
| **SAXO2** | 4.19E-121 | 0.59881923 | 0.996 | 0.961 | 1.01E-116 |
| **MAP9** | 3.95E-133 | 0.595600935 | 0.996 | 0.931 | 9.55E-129 |
| **SSB** | 2.39E-153 | 0.595355937 | 0.997 | 0.976 | 5.76E-149 |
| **LRTOMT** | 3.66E-119 | 0.595200181 | 0.977 | 0.861 | 8.84E-115 |
| **EFHB** | 1.63E-51 | 0.594658642 | 0.784 | 0.634 | 3.94E-47 |
| **SOD2** | 2.54E-33 | 0.594352734 | 0.828 | 0.652 | 6.13E-29 |
| **DNAH12** | 5.22E-104 | 0.592513214 | 0.997 | 0.954 | 1.26E-99 |
| **BSCL2** | 1.93E-105 | 0.590401357 | 1 | 0.937 | 4.66E-101 |
| **AHSA1** | 1.94E-124 | 0.587302944 | 0.976 | 0.847 | 4.69E-120 |
| **CCDC113** | 2.69E-101 | 0.587151454 | 0.997 | 0.955 | 6.5E-97 |
| **RFX2** | 1.65E-111 | 0.585573573 | 0.956 | 0.798 | 3.99E-107 |
| **KLHDC9** | 3.79E-109 | 0.584291843 | 0.858 | 0.572 | 9.15E-105 |
| **MIR200CHG** | 9.97E-84 | 0.5839404 | 0.833 | 0.603 | 2.41E-79 |
| **TMEM107** | 1.98E-104 | 0.583913283 | 0.962 | 0.794 | 4.79E-100 |
| **C20orf96** | 2.31E-113 | 0.583376494 | 0.927 | 0.691 | 5.59E-109 |
| **ENDOG** | 8.45E-102 | 0.583163662 | 0.977 | 0.862 | 2.04E-97 |
| **C1orf87** | 9.16E-86 | 0.582684694 | 0.661 | 0.359 | 2.21E-81 |
| **CNIH2** | 1.65E-114 | 0.582313708 | 0.487 | 0.169 | 3.99E-110 |
| **FDXR** | 4.49E-115 | 0.582137622 | 0.896 | 0.6 | 1.09E-110 |
| **CLIC6** | 2.77E-67 | 0.580686092 | 0.856 | 0.632 | 6.68E-63 |
| **TRIP13** | 5.44E-67 | 0.579113327 | 0.69 | 0.464 | 1.31E-62 |
| **LCA5L** | 9.53E-96 | 0.578236041 | 0.877 | 0.652 | 2.3E-91 |
| **UBAC1** | 5.35E-119 | 0.576293329 | 0.986 | 0.871 | 1.29E-114 |
| **MRPS31** | 3.56E-68 | 0.575118225 | 0.928 | 0.766 | 8.61E-64 |
| **HSPH1** | 9.87E-115 | 0.574558206 | 0.996 | 0.982 | 2.38E-110 |
| **EPPIN** | 2.08E-41 | 0.573775466 | 0.673 | 0.468 | 5.03E-37 |

| | | | | | |
|---|---|---|---|---|---|
| **ROMO1** | 2.14E-133 | 0.573454798 | 0.99 | 0.912 | 5.16E-129 |
| **IFT43** | 7.83E-115 | 0.571857511 | 0.996 | 0.905 | 1.89E-110 |
| **SPEF1** | 7.46E-106 | 0.571836942 | 0.963 | 0.755 | 1.8E-101 |
| **VPS13D** | 1.35E-05 | 0.571786209 | 0.471 | 0.466 | 0.325559625 |
| **NQO1** | 1.63E-85 | 0.570040601 | 0.994 | 0.936 | 3.95E-81 |
| **NR2F1** | 1.24E-131 | 0.569929954 | 0.381 | 0.092 | 2.99E-127 |
| **SPATA4** | 1.71E-108 | 0.568729805 | 0.798 | 0.479 | 4.14E-104 |
| **DRC1** | 6.09E-76 | 0.567350532 | 0.986 | 0.92 | 1.47E-71 |
| **C12orf75** | 1.2E-57 | 0.565672441 | 0.974 | 0.929 | 2.9E-53 |
| **CBY1** | 4.47E-88 | 0.565028478 | 0.922 | 0.706 | 1.08E-83 |
| **PRR7** | 1.8E-97 | 0.564820615 | 0.856 | 0.58 | 4.34E-93 |
| **LCA5** | 9.92E-101 | 0.56307167 | 0.964 | 0.794 | 2.4E-96 |
| **RARRES1** | 3.66E-10 | 0.562338344 | 0.777 | 0.791 | 8.84E-06 |
| **AC007405.3** | 7.74E-86 | 0.560434549 | 0.782 | 0.488 | 1.87E-81 |
| **CCDC34** | 3.35E-98 | 0.557609928 | 0.918 | 0.708 | 8.1E-94 |
| **ZNF688** | 2.55E-74 | 0.555799683 | 0.816 | 0.6 | 6.16E-70 |
| **FBXW9** | 1.88E-82 | 0.554236523 | 0.827 | 0.584 | 4.54E-78 |
| **NME7** | 2.65E-112 | 0.549797261 | 0.963 | 0.788 | 6.4E-108 |
| **PPOX** | 7.24E-67 | 0.549792347 | 0.87 | 0.689 | 1.75E-62 |
| **LINC00240** | 5.38E-97 | 0.549216984 | 0.399 | 0.132 | 1.3E-92 |
| **CCDC160** | 8.23E-74 | 0.548509111 | 0.758 | 0.528 | 1.99E-69 |
| **DNAL4** | 3.13E-83 | 0.547837724 | 0.797 | 0.541 | 7.56E-79 |
| **CFAP206** | 2.36E-111 | 0.546820418 | 0.992 | 0.891 | 5.7E-107 |
| **CIR1** | 1.85E-123 | 0.546482437 | 0.99 | 0.883 | 4.48E-119 |
| **CSPP1** | 4.98E-109 | 0.545768709 | 0.986 | 0.882 | 1.2E-104 |
| **APOBEC4** | 3.51E-87 | 0.544838763 | 0.849 | 0.6 | 8.47E-83 |
| **HACD4** | 5.85E-97 | 0.543673267 | 0.897 | 0.653 | 1.41E-92 |
| **DRC3** | 2.77E-108 | 0.543333671 | 0.992 | 0.934 | 6.69E-104 |
| **CATIP** | 7.73E-93 | 0.540686684 | 0.819 | 0.533 | 1.87E-88 |
| **NUDC** | 7.42E-163 | 0.53924434 | 0.999 | 0.991 | 1.79E-158 |
| **PIH1D3** | 1.24E-94 | 0.539080668 | 0.932 | 0.743 | 2.98E-90 |
| **CCDC33** | 4.39E-92 | 0.538669686 | 0.949 | 0.794 | 1.06E-87 |
| **IQUB** | 4.81E-92 | 0.538589165 | 0.968 | 0.844 | 1.16E-87 |
| **FILIP1** | 7.41E-82 | 0.536777308 | 0.966 | 0.859 | 1.79E-77 |
| **NDUFA3** | 1.85E-110 | 0.536688422 | 0.968 | 0.809 | 4.46E-106 |
| **DNAJA4** | 6.63E-96 | 0.536350465 | 0.998 | 0.913 | 1.6E-91 |
| **B9D1** | 2.1E-99 | 0.535056543 | 0.986 | 0.848 | 5.08E-95 |
| **SNRNP25** | 5.77E-70 | 0.534629572 | 0.818 | 0.578 | 1.39E-65 |

| | | | | | |
|---|---|---|---|---|---|
| **TTC26** | 1.46E-89 | 0.533425625 | 0.921 | 0.729 | 3.52E-85 |
| **UFC1** | 8.77E-133 | 0.532914495 | 1 | 0.997 | 2.12E-128 |
| **CD164L2** | 4.4E-39 | 0.5320215 | 0.847 | 0.689 | 1.06E-34 |
| **HLA-B** | 8.11E-41 | 0.530470575 | 0.928 | 0.842 | 1.96E-36 |
| **CFB** | 0.000158558 | 0.52864708 | 0.549 | 0.52 | 1 |
| **IFT81** | 1.44E-94 | 0.527406148 | 0.963 | 0.849 | 3.47E-90 |
| **ZBED5-AS1** | 5.82E-83 | 0.526748614 | 0.917 | 0.736 | 1.41E-78 |
| **DPY30** | 1.67E-85 | 0.526531559 | 0.991 | 0.948 | 4.04E-81 |
| **FBXO15** | 6.19E-94 | 0.524897909 | 0.817 | 0.527 | 1.5E-89 |
| **GSTA2** | 7.39E-06 | 0.524876061 | 0.738 | 0.72 | 0.178530691 |
| **KIFAP3** | 1.54E-96 | 0.524748653 | 0.877 | 0.626 | 3.72E-92 |
| **CFAP44** | 4.02E-67 | 0.52438605 | 0.983 | 0.923 | 9.72E-63 |
| **IK** | 3.92E-154 | 0.522716606 | 1 | 0.997 | 9.47E-150 |
| **FBXO2** | 2.2E-101 | 0.522681479 | 0.529 | 0.211 | 5.31E-97 |
| **POLR2I** | 3.92E-118 | 0.521442341 | 0.992 | 0.983 | 9.46E-114 |
| **ABHD2** | 2.17E-29 | 0.520599127 | 0.914 | 0.863 | 5.23E-25 |
| **WDR63** | 1.57E-94 | 0.517766198 | 0.974 | 0.855 | 3.78E-90 |
| **AK1** | 3.47E-72 | 0.517416003 | 0.939 | 0.764 | 8.39E-68 |
| **REPIN1** | 2.21E-81 | 0.517252545 | 0.883 | 0.666 | 5.35E-77 |
| **SPATA6** | 2.99E-91 | 0.516964485 | 0.953 | 0.811 | 7.21E-87 |
| **DNAI1** | 2.1E-91 | 0.516596591 | 0.897 | 0.674 | 5.07E-87 |
| **RIBC1** | 2.11E-92 | 0.516322001 | 0.859 | 0.589 | 5.1E-88 |
| **CAPS** | 2.85E-158 | 0.515203423 | 1 | 1 | 6.89E-154 |
| **MAP6** | 6.06E-79 | 0.514610593 | 0.913 | 0.714 | 1.46E-74 |
| **RPA3** | 1.95E-101 | 0.514369232 | 0.986 | 0.867 | 4.71E-97 |
| **B9D2** | 1.02E-71 | 0.514097026 | 0.868 | 0.663 | 2.47E-67 |
| **CKB** | 1.94E-60 | 0.513521247 | 0.992 | 0.968 | 4.69E-56 |
| **CLDN3** | 4.19E-44 | 0.513504586 | 0.668 | 0.48 | 1.01E-39 |
| **IFT27** | 2.04E-93 | 0.511103167 | 0.989 | 0.875 | 4.92E-89 |
| **ACYP1** | 6.69E-85 | 0.51075419 | 0.903 | 0.673 | 1.62E-80 |
| **STOX1** | 8.68E-90 | 0.510285484 | 0.957 | 0.813 | 2.1E-85 |
| **MORN3** | 2.07E-75 | 0.509857297 | 0.873 | 0.678 | 4.99E-71 |
| **NDUFA1** | 6.22E-127 | 0.506417122 | 0.992 | 0.965 | 1.5E-122 |
| **AKR7A2** | 6.11E-77 | 0.50622069 | 0.884 | 0.68 | 1.48E-72 |
| **MS4A8** | 3.61E-82 | 0.505214428 | 0.976 | 0.851 | 8.72E-78 |
| **UCP2** | 6.56E-95 | 0.504883964 | 0.994 | 0.968 | 1.58E-90 |
| **LRRC61** | 5.58E-46 | 0.5036165 | 0.542 | 0.343 | 1.35E-41 |
| **GCC2** | 1.5E-104 | 0.503525335 | 0.994 | 0.965 | 3.62E-100 |

| | | | | | |
|---|---|---|---|---|---|
| **PKIG** | 3.02E-81 | 0.503495786 | 0.919 | 0.721 | 7.28E-77 |
| **SYT5** | 4.65E-59 | 0.503249124 | 0.266 | 0.088 | 1.12E-54 |
| **LRRC49** | 1.53E-76 | 0.503174394 | 0.873 | 0.678 | 3.69E-72 |
| **IFT46** | 8.81E-76 | 0.502659474 | 0.839 | 0.625 | 2.13E-71 |
| **KATNAL2** | 5.48E-89 | 0.502373119 | 0.719 | 0.413 | 1.32E-84 |
| **C4orf48** | 1.64E-83 | 0.499173687 | 0.984 | 0.898 | 3.97E-79 |
| **LINC01513** | 8.85E-81 | 0.498930113 | 0.802 | 0.544 | 2.14E-76 |
| **ATP5IF1** | 9.34E-105 | 0.497888205 | 0.996 | 0.999 | 2.26E-100 |
| **TMA7** | 3.95E-135 | 0.497306725 | 0.988 | 0.989 | 9.54E-131 |
| **CFAP43** | 9.04E-78 | 0.49692009 | 0.994 | 0.96 | 2.18E-73 |
| **RPP38** | 3.48E-83 | 0.496866643 | 0.88 | 0.648 | 8.41E-79 |
| **COX17** | 4.21E-97 | 0.495746643 | 0.988 | 0.909 | 1.02E-92 |
| **CFAP77** | 2.1E-80 | 0.494993841 | 0.892 | 0.686 | 5.08E-76 |
| **PPP1R15A** | 6.39E-78 | 0.494795824 | 0.928 | 0.743 | 1.54E-73 |
| **AL022068.1** | 1.12E-76 | 0.493843883 | 0.897 | 0.735 | 2.71E-72 |
| **TTLL10** | 3.26E-64 | 0.492621146 | 0.763 | 0.527 | 7.87E-60 |
| **ARMH4** | 3.24E-65 | 0.492432589 | 0.551 | 0.295 | 7.83E-61 |
| **DIXDC1** | 9E-70 | 0.490892272 | 0.848 | 0.628 | 2.17E-65 |
| **GPR162** | 2.25E-83 | 0.490167621 | 0.869 | 0.629 | 5.44E-79 |
| **SPACA9** | 1.39E-66 | 0.489493948 | 0.976 | 0.876 | 3.35E-62 |
| **CFAP52** | 2.62E-96 | 0.487189496 | 0.986 | 0.878 | 6.34E-92 |
| **SAP30** | 4.26E-75 | 0.486553358 | 0.733 | 0.461 | 1.03E-70 |
| **ZMAT2** | 1.26E-77 | 0.485990065 | 0.924 | 0.751 | 3.05E-73 |
| **LINC02265** | 8.76E-88 | 0.48453571 | 0.638 | 0.318 | 2.12E-83 |
| **MEIG1** | 6.46E-92 | 0.484240219 | 0.7 | 0.371 | 1.56E-87 |
| **DCDC2B** | 1.21E-64 | 0.482850495 | 0.733 | 0.477 | 2.92E-60 |
| **NUCB2** | 6.16E-61 | 0.482803428 | 0.998 | 0.994 | 1.49E-56 |
| **FAM104B** | 4.19E-67 | 0.482311268 | 0.922 | 0.757 | 1.01E-62 |
| **PLPPR3** | 7.74E-53 | 0.481821341 | 0.776 | 0.549 | 1.87E-48 |
| **SMYD2** | 1.03E-77 | 0.479826209 | 0.89 | 0.694 | 2.49E-73 |
| **DNPH1** | 1.99E-83 | 0.479426778 | 0.986 | 0.917 | 4.82E-79 |
| **SMKR1** | 2.52E-94 | 0.479231979 | 0.704 | 0.368 | 6.08E-90 |
| **PLAC8** | 9.36E-72 | 0.478522859 | 0.996 | 0.984 | 2.26E-67 |
| **TMEM212** | 2.54E-66 | 0.476757729 | 0.766 | 0.53 | 6.14E-62 |
| **LRRC73** | 5.27E-71 | 0.476239987 | 0.78 | 0.499 | 1.27E-66 |
| **BAIAP3** | 8.44E-65 | 0.474483845 | 0.649 | 0.379 | 2.04E-60 |
| **MPC2** | 1.92E-78 | 0.474358842 | 0.981 | 0.905 | 4.64E-74 |
| **UBXN10** | 1.19E-85 | 0.474320837 | 0.97 | 0.853 | 2.88E-81 |

| | | | | | |
|---|---|---|---|---|---|
| **LINC00467** | 1.12E-70 | 0.473811766 | 0.821 | 0.594 | 2.71E-66 |
| **ARFIP2** | 8.56E-42 | 0.472582021 | 0.653 | 0.468 | 2.07E-37 |
| **MAPK8IP1** | 4.43E-39 | 0.471751946 | 0.74 | 0.552 | 1.07E-34 |
| **TMEM231** | 2.53E-79 | 0.47107111 | 0.987 | 0.928 | 6.11E-75 |
| **PET100** | 9.17E-82 | 0.470611645 | 0.954 | 0.801 | 2.21E-77 |
| **KCNRG** | 1.32E-69 | 0.470254853 | 0.856 | 0.637 | 3.2E-65 |
| **TCTN1** | 3.21E-97 | 0.468699547 | 0.993 | 0.921 | 7.75E-93 |
| **KIAA2012** | 1.09E-61 | 0.466995099 | 0.796 | 0.556 | 2.63E-57 |
| **CFAP161** | 2.41E-72 | 0.466764993 | 0.784 | 0.513 | 5.83E-68 |
| **DNAJA1** | 2.93E-101 | 0.466120975 | 0.997 | 0.98 | 7.07E-97 |
| **CARS** | 8.09E-31 | 0.464021672 | 0.881 | 0.771 | 1.95E-26 |
| **BUD23** | 8.32E-59 | 0.463227357 | 0.939 | 0.796 | 2.01E-54 |
| **JOSD2** | 3.28E-57 | 0.461782654 | 0.922 | 0.778 | 7.91E-53 |
| **ERCC1** | 8.25E-32 | 0.46152185 | 0.827 | 0.697 | 1.99E-27 |
| **C6orf118** | 1.03E-62 | 0.458868622 | 0.979 | 0.88 | 2.5E-58 |
| **COQ4** | 9.02E-67 | 0.458641434 | 0.961 | 0.811 | 2.18E-62 |
| **C7orf57** | 5.24E-70 | 0.458347317 | 0.943 | 0.759 | 1.27E-65 |
| **WDR34** | 7.87E-71 | 0.458284464 | 0.973 | 0.866 | 1.9E-66 |
| **MIPEP** | 1.77E-72 | 0.457641178 | 0.861 | 0.657 | 4.27E-68 |
| **CENPS** | 2.75E-67 | 0.457197843 | 0.818 | 0.595 | 6.64E-63 |
| **CEP290** | 2.32E-62 | 0.456654966 | 0.924 | 0.779 | 5.6E-58 |
| **EFHC2** | 6.17E-67 | 0.45647247 | 0.91 | 0.728 | 1.49E-62 |
| **SRD5A2** | 2.98E-77 | 0.454690198 | 0.694 | 0.389 | 7.21E-73 |
| **PPP1R14C** | 4.47E-74 | 0.454560784 | 0.883 | 0.65 | 1.08E-69 |
| **ZFHX2** | 1.68E-64 | 0.454380176 | 0.79 | 0.547 | 4.07E-60 |
| **HLA-DRA** | 2.25E-22 | 0.453827712 | 0.68 | 0.541 | 5.44E-18 |
| **MSMB** | 4.39E-16 | 0.453731255 | 0.108 | 0.238 | 1.06E-11 |
| **DNAI2** | 8.07E-68 | 0.452928099 | 0.848 | 0.615 | 1.95E-63 |
| **HMGN2** | 1.7E-69 | 0.452155322 | 0.964 | 0.883 | 4.11E-65 |
| **ANKMY1** | 3.05E-55 | 0.451879347 | 0.821 | 0.656 | 7.36E-51 |
| **CNTRL** | 6.62E-68 | 0.451821931 | 0.938 | 0.779 | 1.6E-63 |
| **GLIPR2** | 2.49E-70 | 0.451762618 | 0.746 | 0.48 | 6.03E-66 |
| **CALM2** | 4.23E-119 | 0.450734335 | 0.999 | 0.999 | 1.02E-114 |
| **VIM-AS1** | 3.18E-42 | 0.450530444 | 0.641 | 0.437 | 7.69E-38 |
| **PRR34-AS1** | 5.08E-71 | 0.449711079 | 0.728 | 0.452 | 1.23E-66 |
| **TSNAXIP1** | 5.84E-66 | 0.449459037 | 0.849 | 0.621 | 1.41E-61 |
| **STAU1** | 2.67E-54 | 0.44917167 | 0.907 | 0.774 | 6.44E-50 |
| **LRRC71** | 3.07E-68 | 0.449056735 | 0.842 | 0.613 | 7.42E-64 |

| | | | | | |
|---|---|---|---|---|---|
| **NDUFAF3** | 3.63E-74 | 0.447564504 | 0.952 | 0.816 | 8.76E-70 |
| **IQCB1** | 3.11E-67 | 0.447190349 | 0.837 | 0.616 | 7.51E-63 |
| **ACAT1** | 5.81E-69 | 0.446894032 | 0.704 | 0.429 | 1.4E-64 |
| **DYNC2LI1** | 6.01E-71 | 0.445847575 | 0.946 | 0.782 | 1.45E-66 |
| **SNORC** | 3.78E-48 | 0.445836819 | 0.284 | 0.112 | 9.13E-44 |
| **RSPH14** | 7.12E-76 | 0.445433133 | 0.709 | 0.412 | 1.72E-71 |
| **BASP1** | 4.51E-81 | 0.44410242 | 1 | 0.994 | 1.09E-76 |
| **ERICH3** | 5.43E-64 | 0.443887998 | 1 | 0.981 | 1.31E-59 |
| **CCDC181** | 2.98E-42 | 0.443585835 | 0.749 | 0.587 | 7.19E-38 |
| **RBKS** | 7.47E-78 | 0.44298978 | 0.748 | 0.456 | 1.81E-73 |
| **MLF 1.00** | 1.18E-92 | 0.442413258 | 0.996 | 0.969 | 2.85E-88 |
| **EFCAB11** | 4.16E-70 | 0.442393344 | 0.787 | 0.53 | 1E-65 |
| **AC023300.2** | 1.14E-35 | 0.442336707 | 0.733 | 0.56 | 2.76E-31 |
| **AL035701.1** | 1.84E-55 | 0.441879192 | 0.729 | 0.504 | 4.44E-51 |
| **SPATA33** | 6.32E-70 | 0.441595071 | 0.932 | 0.779 | 1.53E-65 |
| **SLC25A4** | 3.85E-67 | 0.44125653 | 0.93 | 0.779 | 9.3E-63 |
| **C6orf52** | 2.84E-70 | 0.441053581 | 0.672 | 0.391 | 6.85E-66 |
| **FUZ** | 2.34E-61 | 0.440521567 | 0.772 | 0.55 | 5.66E-57 |
| **TSGA10** | 7.84E-69 | 0.439764078 | 0.967 | 0.86 | 1.89E-64 |
| **ISG15** | 2.65E-12 | 0.439267716 | 0.447 | 0.343 | 6.4E-08 |
| **SMIM41** | 8.33E-72 | 0.438319213 | 0.607 | 0.314 | 2.01E-67 |
| **IFT88** | 2.95E-80 | 0.438293251 | 0.982 | 0.883 | 7.14E-76 |
| **MT-ATP8** | 1.08E-33 | 0.436986983 | 0.62 | 0.466 | 2.6E-29 |
| **RIPOR2** | 5.04E-51 | 0.436511765 | 0.899 | 0.774 | 1.22E-46 |
| **PSMB10** | 5.96E-68 | 0.436349097 | 0.892 | 0.676 | 1.44E-63 |
| **WDR60** | 5.51E-76 | 0.436159347 | 0.987 | 0.909 | 1.33E-71 |
| **C22orf23** | 2.43E-85 | 0.436129615 | 0.471 | 0.189 | 5.87E-81 |
| **UGDH** | 6.04E-55 | 0.435260158 | 0.91 | 0.735 | 1.46E-50 |
| **FABP6** | 2E-27 | 0.433450916 | 0.369 | 0.217 | 4.84E-23 |
| **FAM47E** | 1.27E-64 | 0.433294454 | 0.857 | 0.638 | 3.07E-60 |
| **PSME2** | 6.66E-53 | 0.433288944 | 0.889 | 0.735 | 1.61E-48 |
| **CCDC189** | 7.8E-60 | 0.433285056 | 0.843 | 0.625 | 1.88E-55 |
| **ATP12A** | 1.42E-08 | 0.432514594 | 0.313 | 0.242 | 0.000342828 |
| **AC004832.1** | 3.96E-57 | 0.431234714 | 0.846 | 0.637 | 9.58E-53 |
| **C2orf81** | 2.53E-68 | 0.430997206 | 0.807 | 0.551 | 6.12E-64 |
| **RPGRIP1L** | 1.18E-60 | 0.430889993 | 0.91 | 0.759 | 2.85E-56 |
| **BUD31** | 4.39E-74 | 0.430752215 | 0.973 | 0.847 | 1.06E-69 |
| **DERL3** | 7.43E-50 | 0.430116463 | 0.447 | 0.222 | 1.8E-45 |

| | | | | | |
|---|---|---|---|---|---|
| **DRC7** | 1.61E-71 | 0.429499563 | 0.854 | 0.605 | 3.89E-67 |
| **IRF9** | 9.62E-67 | 0.428362545 | 0.847 | 0.601 | 2.32E-62 |
| **MRVI1-AS1** | 1E-73 | 0.427643189 | 0.582 | 0.289 | 2.42E-69 |
| **AC244090.1** | 6.65E-40 | 0.427385193 | 0.857 | 0.725 | 1.61E-35 |
| **ENKD1** | 7.28E-59 | 0.426532089 | 0.849 | 0.652 | 1.76E-54 |
| **LPAR3** | 3.85E-43 | 0.42582533 | 0.663 | 0.477 | 9.31E-39 |
| **HLA-DRB1** | 1.88E-19 | 0.425813246 | 0.527 | 0.396 | 4.53E-15 |
| **GLB1L** | 8.74E-69 | 0.42547302 | 0.669 | 0.382 | 2.11E-64 |
| **MNAT1** | 1.89E-72 | 0.424851098 | 0.922 | 0.709 | 4.56E-68 |
| **C1orf158** | 7.85E-70 | 0.424221652 | 0.844 | 0.585 | 1.9E-65 |
| **TEKT2** | 7.02E-61 | 0.422693245 | 0.937 | 0.796 | 1.7E-56 |
| **TNFAIP8L1** | 2.95E-72 | 0.422683732 | 0.982 | 0.894 | 7.12E-68 |
| **DMKN** | 3.14E-62 | 0.421101529 | 0.994 | 0.926 | 7.57E-58 |
| **FOCAD** | 1.47E-62 | 0.420373404 | 0.763 | 0.524 | 3.55E-58 |
| **TMEM232** | 2.69E-57 | 0.420347834 | 0.942 | 0.799 | 6.49E-53 |
| **PHTF1** | 7.02E-57 | 0.419722146 | 0.914 | 0.743 | 1.7E-52 |
| **MX1** | 6.66E-25 | 0.419168382 | 0.639 | 0.482 | 1.61E-20 |
| **SPATS2L** | 1.13E-55 | 0.419164406 | 0.922 | 0.813 | 2.73E-51 |
| **MUC15** | 5.52E-63 | 0.419065343 | 0.948 | 0.822 | 1.33E-58 |
| **NELFE** | 5.55E-69 | 0.418501036 | 0.907 | 0.705 | 1.34E-64 |
| **CCDC81** | 5.49E-53 | 0.418058011 | 0.418 | 0.198 | 1.33E-48 |
| **MRPS33** | 8.41E-49 | 0.41573918 | 0.916 | 0.766 | 2.03E-44 |
| **RARRES2** | 4.62E-31 | 0.415694259 | 0.386 | 0.222 | 1.12E-26 |
| **ITGB3BP** | 2.54E-68 | 0.414015399 | 0.623 | 0.347 | 6.13E-64 |
| **LINC01571** | 1.68E-38 | 0.413303801 | 0.523 | 0.329 | 4.05E-34 |
| **C4orf47** | 2.92E-66 | 0.41203465 | 0.811 | 0.568 | 7.04E-62 |
| **RGS22** | 4.25E-50 | 0.411520572 | 0.631 | 0.394 | 1.03E-45 |
| **CIB1** | 1.41E-73 | 0.410635315 | 0.999 | 1 | 3.4E-69 |
| **SPAG8** | 3.32E-58 | 0.41061003 | 0.928 | 0.778 | 8.01E-54 |
| **MUC13** | 0.107521752 | 0.41029343 | 0.196 | 0.181 | 1 |
| **RPS29** | 8.99E-89 | 0.409753028 | 0.989 | 0.995 | 2.17E-84 |
| **ZMYND12** | 1.32E-81 | 0.409748988 | 0.626 | 0.308 | 3.19E-77 |
| **NDUFC1** | 1.32E-76 | 0.408772555 | 0.982 | 0.931 | 3.18E-72 |
| **H2AFJ** | 3.21E-65 | 0.408437083 | 0.996 | 0.975 | 7.74E-61 |
| **ZNF487** | 3.54E-49 | 0.408424545 | 0.913 | 0.741 | 8.54E-45 |
| **DNAH2** | 5.61E-54 | 0.407951251 | 0.818 | 0.603 | 1.35E-49 |
| **ADGB** | 2.56E-52 | 0.407536715 | 0.883 | 0.718 | 6.18E-48 |
| **EIPR1** | 5.95E-50 | 0.406728319 | 0.644 | 0.419 | 1.44E-45 |

| | | | | | |
|---|---|---|---|---|---|
| **TP53TG1** | 7.96E-42 | 0.40580056 | 0.892 | 0.748 | 1.92E-37 |
| **PLSCR1** | 1.15E-46 | 0.405519533 | 0.843 | 0.643 | 2.78E-42 |
| **LKAAEAR1** | 6.21E-60 | 0.405327912 | 0.274 | 0.092 | 1.5E-55 |
| **MINDY4** | 4.63E-72 | 0.40513407 | 0.671 | 0.375 | 1.12E-67 |
| **HAGHL** | 5.7E-58 | 0.404738674 | 0.926 | 0.746 | 1.38E-53 |
| **MAP1B** | 6.73E-41 | 0.404408294 | 0.899 | 0.743 | 1.63E-36 |
| **CCDC170** | 3.69E-25 | 0.403899981 | 1 | 0.993 | 8.91E-21 |
| **DALRD3** | 6.55E-61 | 0.402900191 | 0.88 | 0.667 | 1.58E-56 |
| **SSBP4** | 4.38E-61 | 0.402830277 | 0.976 | 0.859 | 1.06E-56 |
| **COL28A1** | 1.17E-29 | 0.402552697 | 0.5 | 0.338 | 2.83E-25 |
| **MT-CO2** | 1.14E-104 | 0.402319998 | 1 | 1 | 2.74E-100 |
| **DUSP14** | 1.8E-51 | 0.401731296 | 0.821 | 0.627 | 4.34E-47 |
| **P4HTM** | 3.11E-61 | 0.401465057 | 0.974 | 0.889 | 7.51E-57 |
| **DTNA** | 6.7E-87 | 0.400583247 | 0.466 | 0.176 | 1.62E-82 |
| **SLIRP** | 1.21E-69 | 0.400507423 | 0.941 | 0.758 | 2.92E-65 |
| **CFAP58** | 2.27E-58 | 0.399097776 | 0.782 | 0.534 | 5.49E-54 |
| **CCDC17** | 4.18E-63 | 0.398679631 | 0.99 | 0.944 | 1.01E-58 |
| **RPGR** | 4.92E-59 | 0.398505427 | 0.942 | 0.793 | 1.19E-54 |
| **ARMC2** | 2.53E-66 | 0.398409527 | 0.818 | 0.559 | 6.11E-62 |
| **KIF19** | 2.7E-61 | 0.39606122 | 0.881 | 0.659 | 6.53E-57 |
| **KRT7** | 6.59E-07 | 0.395616405 | 0.206 | 0.317 | 0.015912032 |
| **POLR3K** | 1.08E-58 | 0.393078705 | 0.731 | 0.484 | 2.62E-54 |
| **BTC** | 3.89E-58 | 0.392795394 | 0.744 | 0.488 | 9.4E-54 |
| **HINT2** | 1.54E-49 | 0.39214531 | 0.907 | 0.724 | 3.72E-45 |
| **NEK11** | 1.62E-61 | 0.391944421 | 0.941 | 0.778 | 3.92E-57 |
| **NAT1** | 4.4E-52 | 0.391725981 | 0.697 | 0.464 | 1.06E-47 |
| **CYP4B1** | 3.39E-42 | 0.391508022 | 0.992 | 0.997 | 8.2E-38 |
| **SMIM5** | 1.15E-62 | 0.391487154 | 0.69 | 0.413 | 2.79E-58 |
| **SMPD2** | 1.08E-52 | 0.391471315 | 0.814 | 0.599 | 2.61E-48 |
| **PIP** | 2.31E-45 | 0.391195537 | 0.223 | 0.077 | 5.57E-41 |
| **CDKN2A** | 4.56E-25 | 0.390785166 | 0.689 | 0.527 | 1.1E-20 |
| **DYDC1** | 9.82E-69 | 0.390351821 | 0.614 | 0.324 | 2.37E-64 |
| **KIF6** | 2.87E-67 | 0.389974345 | 0.651 | 0.364 | 6.92E-63 |
| **FAM92A** | 2.75E-51 | 0.389897561 | 0.69 | 0.463 | 6.64E-47 |
| **GSTO1** | 7.14E-38 | 0.389839268 | 0.718 | 0.534 | 1.72E-33 |
| **NMI** | 6.52E-59 | 0.389761667 | 0.679 | 0.413 | 1.58E-54 |
| **TM9SF1** | 4.08E-59 | 0.388358609 | 0.933 | 0.771 | 9.86E-55 |
| **AL451165.2** | 8.33E-60 | 0.387351521 | 0.612 | 0.355 | 2.01E-55 |

| | | | | | |
|---|---|---|---|---|---|
| **LINC01091** | 2.45E-58 | 0.386913979 | 0.559 | 0.302 | 5.91E-54 |
| **TOMM7** | 3.69E-94 | 0.386617775 | 0.991 | 0.993 | 8.91E-90 |
| **DNAH10** | 8.06E-49 | 0.386370796 | 0.952 | 0.848 | 1.95E-44 |
| **CCDC82** | 1.02E-55 | 0.386218313 | 0.869 | 0.647 | 2.47E-51 |
| **BBS5** | 3.29E-56 | 0.38616585 | 0.772 | 0.555 | 7.94E-52 |
| **PLAAT3** | 2.46E-49 | 0.384665712 | 0.889 | 0.718 | 5.93E-45 |
| **PALMD** | 2.85E-38 | 0.384310545 | 0.831 | 0.661 | 6.87E-34 |
| **SPEF2** | 1.63E-65 | 0.38395954 | 0.996 | 0.971 | 3.93E-61 |
| **CENPK** | 6.44E-75 | 0.383702225 | 0.478 | 0.205 | 1.55E-70 |
| **IQCK** | 9.51E-56 | 0.382959499 | 0.937 | 0.789 | 2.3E-51 |
| **CD74** | 5.41E-29 | 0.382907317 | 0.96 | 0.903 | 1.31E-24 |
| **WRAP53** | 7.77E-57 | 0.382820064 | 0.769 | 0.52 | 1.88E-52 |
| **ANKRD42** | 7.16E-52 | 0.382685251 | 0.894 | 0.735 | 1.73E-47 |
| **CCDC148** | 1.02E-59 | 0.382009789 | 0.751 | 0.497 | 2.46E-55 |
| **CYGB** | 2.38E-38 | 0.381893242 | 0.294 | 0.135 | 5.75E-34 |
| **RPL38** | 6.79E-100 | 0.380968127 | 0.993 | 0.993 | 1.64E-95 |
| **CFAP46** | 5.76E-54 | 0.380953614 | 0.939 | 0.785 | 1.39E-49 |
| **TMEM14B** | 1.32E-68 | 0.380775224 | 0.977 | 0.934 | 3.18E-64 |
| **PLCH1** | 3.63E-57 | 0.38050407 | 0.752 | 0.503 | 8.76E-53 |
| **DCDC2** | 7.09E-72 | 0.380214254 | 0.528 | 0.246 | 1.71E-67 |
| **RAB36** | 3.29E-51 | 0.380126022 | 0.918 | 0.772 | 7.95E-47 |
| **HSD11B1L** | 7.73E-41 | 0.379803734 | 0.761 | 0.569 | 1.87E-36 |
| **SMIM27** | 1.29E-56 | 0.379472663 | 0.686 | 0.437 | 3.1E-52 |
| **CFAP65** | 2.01E-52 | 0.379225163 | 0.891 | 0.727 | 4.85E-48 |
| **CCDC66** | 4.33E-44 | 0.379020013 | 0.774 | 0.597 | 1.05E-39 |
| **MAP3K19** | 2.18E-53 | 0.378714506 | 0.991 | 0.934 | 5.26E-49 |
| **TSTD1** | 2.22E-69 | 0.37857421 | 0.992 | 0.983 | 5.35E-65 |
| **MAATS1** | 2.52E-54 | 0.378541182 | 0.961 | 0.838 | 6.1E-50 |
| **SPATA7** | 9.35E-60 | 0.378400819 | 0.688 | 0.43 | 2.26E-55 |
| **ANKRD26** | 1.17E-45 | 0.377730544 | 0.773 | 0.578 | 2.83E-41 |
| **HOXB-AS1** | 6.45E-24 | 0.377118435 | 0.256 | 0.135 | 1.56E-19 |
| **TTC30B** | 2.36E-40 | 0.376833669 | 0.721 | 0.526 | 5.7E-36 |
| **CCDC138** | 4.05E-49 | 0.376764675 | 0.713 | 0.487 | 9.79E-45 |
| **RBIS** | 4.72E-56 | 0.376676928 | 0.884 | 0.677 | 1.14E-51 |
| **DNAL1** | 1.09E-56 | 0.37666702 | 0.984 | 0.909 | 2.63E-52 |
| **S100A6** | 2.27E-53 | 0.376586091 | 0.998 | 0.999 | 5.49E-49 |
| **MICOS10** | 1.4E-59 | 0.376264121 | 0.993 | 0.977 | 3.37E-55 |
| **DZIP1L** | 9.32E-45 | 0.375430977 | 0.843 | 0.658 | 2.25E-40 |

| | | | | | |
|---|---|---|---|---|---|
| **CEP162** | 4.44E-50 | 0.375242358 | 0.894 | 0.723 | 1.07E-45 |
| **C21orf58** | 1.49E-37 | 0.374693292 | 0.968 | 0.889 | 3.6E-33 |
| **UCKL1-AS1** | 1.14E-46 | 0.373409846 | 0.713 | 0.489 | 2.75E-42 |
| **TCTE1** | 9.18E-73 | 0.373302951 | 0.612 | 0.311 | 2.22E-68 |
| **TSPAN1** | 2.76E-67 | 0.372743948 | 1 | 1 | 6.66E-63 |
| **PLEKHB1** | 1.54E-41 | 0.372445118 | 0.796 | 0.622 | 3.71E-37 |
| **CFAP100** | 3.37E-54 | 0.37233377 | 0.868 | 0.668 | 8.14E-50 |
| **ZNF106** | 3.14E-32 | 0.372186742 | 0.956 | 0.877 | 7.57E-28 |
| **10-Mar** | 6.57E-62 | 0.371987889 | 0.64 | 0.36 | 1.59E-57 |
| **SLC51B** | 3.35E-70 | 0.371927077 | 0.503 | 0.23 | 8.09E-66 |
| **CHCHD5** | 9.55E-49 | 0.370964311 | 0.927 | 0.776 | 2.31E-44 |
| **CTSS** | 1.93E-69 | 0.370884077 | 0.998 | 0.989 | 4.66E-65 |
| **SULT1A1** | 2.16E-49 | 0.370865672 | 0.791 | 0.558 | 5.23E-45 |
| **CEP83** | 1.24E-51 | 0.370801699 | 0.85 | 0.659 | 3.01E-47 |
| **CCDC151** | 8.38E-66 | 0.370669339 | 0.699 | 0.402 | 2.02E-61 |
| **GAS2L2** | 1.16E-51 | 0.369957166 | 0.676 | 0.423 | 2.81E-47 |
| **STOML2** | 1.04E-42 | 0.369564595 | 0.868 | 0.706 | 2.51E-38 |
| **MOK** | 1.49E-56 | 0.36929915 | 0.959 | 0.822 | 3.59E-52 |
| **KIF3A** | 4.1E-55 | 0.368062525 | 0.956 | 0.838 | 9.91E-51 |
| **PPP1R7** | 3.44E-47 | 0.368056782 | 0.979 | 0.925 | 8.32E-43 |
| **DZIP1** | 1.58E-46 | 0.367970393 | 0.653 | 0.42 | 3.83E-42 |
| **ZDHHC1** | 1.34E-42 | 0.367406544 | 0.799 | 0.618 | 3.23E-38 |
| **PIN1** | 7.74E-43 | 0.367381785 | 0.884 | 0.702 | 1.87E-38 |
| **PLA2G10** | 5.39E-35 | 0.366652613 | 0.762 | 0.605 | 1.3E-30 |
| **GTF2F1** | 1.02E-54 | 0.366491071 | 0.791 | 0.537 | 2.47E-50 |
| **C2orf74** | 1.19E-25 | 0.365990874 | 0.389 | 0.241 | 2.87E-21 |
| **SLC22A4** | 2.38E-49 | 0.365622186 | 0.866 | 0.686 | 5.74E-45 |
| **TCEAL3** | 3.52E-40 | 0.365335663 | 0.762 | 0.59 | 8.51E-36 |
| **STPG1** | 2.4E-54 | 0.36435856 | 0.727 | 0.484 | 5.81E-50 |
| **AL035420.3** | 1.6E-62 | 0.364262459 | 0.473 | 0.222 | 3.87E-58 |
| **HSBP1** | 2.25E-83 | 0.36394041 | 0.997 | 0.996 | 5.43E-79 |
| **SYTL3** | 1.59E-44 | 0.363649654 | 0.746 | 0.532 | 3.85E-40 |
| **GAS8** | 1.16E-51 | 0.363353147 | 0.793 | 0.582 | 2.81E-47 |
| **TMEM256** | 8.8E-49 | 0.363094685 | 0.874 | 0.683 | 2.13E-44 |
| **HEBP2** | 2.71E-39 | 0.362777076 | 0.916 | 0.816 | 6.55E-35 |
| **CFAP20** | 5.94E-46 | 0.362468185 | 0.831 | 0.641 | 1.44E-41 |
| **KIAA1257** | 4.51E-59 | 0.362425829 | 0.591 | 0.326 | 1.09E-54 |
| **TRAF3IP1** | 4.36E-40 | 0.362323839 | 0.986 | 0.935 | 1.05E-35 |

| | | | | | |
|---|---|---|---|---|---|
| **UBXN11** | 2.54E-50 | 0.361892086 | 0.877 | 0.712 | 6.15E-46 |
| **C10orf95** | 4.63E-57 | 0.360595695 | 0.648 | 0.38 | 1.12E-52 |
| **ZNHIT2** | 1.21E-43 | 0.359991653 | 0.804 | 0.623 | 2.93E-39 |
| **MRPL18** | 7.59E-55 | 0.35982945 | 0.916 | 0.746 | 1.83E-50 |
| **ZCRB1** | 6.46E-45 | 0.359762968 | 0.94 | 0.808 | 1.56E-40 |
| **SURF2** | 1.21E-36 | 0.359365255 | 0.802 | 0.629 | 2.92E-32 |
| **IL5RA** | 2.51E-51 | 0.358886616 | 0.718 | 0.471 | 6.07E-47 |
| **CCDC13** | 9.76E-52 | 0.35817108 | 0.687 | 0.44 | 2.36E-47 |
| **DNAJB13** | 2.7E-58 | 0.357599033 | 0.824 | 0.559 | 6.52E-54 |
| **ZNF474** | 8.65E-53 | 0.35720911 | 0.713 | 0.465 | 2.09E-48 |
| **SELENOW** | 5.53E-47 | 0.356916588 | 0.958 | 0.847 | 1.34E-42 |
| **PAIP2** | 5.3E-52 | 0.356903107 | 0.951 | 0.819 | 1.28E-47 |
| **SNAPC1** | 1.36E-50 | 0.356807879 | 0.694 | 0.451 | 3.29E-46 |
| **RHPN2** | 4.82E-40 | 0.356459441 | 0.807 | 0.638 | 1.16E-35 |
| **FAM174A** | 4.83E-59 | 0.356438448 | 0.984 | 0.878 | 1.17E-54 |
| **MAPRE3** | 1.19E-49 | 0.355928895 | 0.917 | 0.749 | 2.88E-45 |
| **VDAC3** | 3.33E-53 | 0.355779919 | 0.963 | 0.832 | 8.04E-49 |
| **PTGES3** | 2.71E-69 | 0.355715186 | 0.996 | 0.988 | 6.54E-65 |
| **TUBB4B** | 4.29E-45 | 0.355514391 | 1 | 1 | 1.04E-40 |
| **YWHAH** | 5.9E-42 | 0.355352216 | 0.888 | 0.719 | 1.42E-37 |
| **ARL13B** | 1.7E-45 | 0.355197029 | 0.747 | 0.542 | 4.1E-41 |
| **CYSTM1** | 2.64E-69 | 0.355112177 | 0.991 | 0.985 | 6.37E-65 |
| **PPP1R36** | 1.36E-49 | 0.354983837 | 0.532 | 0.294 | 3.29E-45 |
| **BOLA3** | 6.06E-45 | 0.354898737 | 0.801 | 0.604 | 1.46E-40 |
| **MAP1LC3A** | 3.57E-39 | 0.354734786 | 0.531 | 0.335 | 8.62E-35 |
| **PSME1** | 9.43E-51 | 0.354501726 | 0.916 | 0.777 | 2.28E-46 |
| **MRPS21** | 2.18E-55 | 0.35416526 | 0.976 | 0.894 | 5.26E-51 |
| **NGRN** | 3.18E-57 | 0.354034712 | 0.984 | 0.905 | 7.67E-53 |
| **PCYT2** | 1.81E-45 | 0.353723861 | 0.652 | 0.427 | 4.37E-41 |
| **PRR13** | 2.87E-51 | 0.353519967 | 0.954 | 0.807 | 6.93E-47 |
| **CHORDC1** | 6.81E-49 | 0.353333598 | 0.728 | 0.498 | 1.64E-44 |
| **TP53AIP1** | 1.92E-48 | 0.353152711 | 0.393 | 0.184 | 4.63E-44 |
| **TOGARAM2** | 2.14E-46 | 0.352132284 | 0.802 | 0.61 | 5.16E-42 |
| **FAM86B1** | 2.44E-116 | 0.352043664 | 0.378 | 0.097 | 5.88E-112 |
| **SLC44A4** | 3.55E-48 | 0.351966306 | 0.996 | 0.96 | 8.56E-44 |
| **DNAAF3** | 2.46E-38 | 0.35188929 | 0.881 | 0.723 | 5.94E-34 |
| **MRPS18C** | 3.36E-46 | 0.351608481 | 0.88 | 0.696 | 8.12E-42 |
| **ALOX15** | 3.53E-19 | 0.351084824 | 0.911 | 0.891 | 8.53E-15 |

| | | | | | |
|---|---|---|---|---|---|
| **NAA20** | 9.88E-48 | 0.35071249 | 0.953 | 0.831 | 2.39E-43 |
| **PPID** | 2.71E-41 | 0.350555721 | 0.748 | 0.565 | 6.55E-37 |
| **ADGRE5** | 5.97E-36 | 0.350329053 | 0.572 | 0.387 | 1.44E-31 |
| **DHRS7B** | 2.5E-49 | 0.350062766 | 0.751 | 0.514 | 6.03E-45 |
| **SPATS1** | 4.18E-87 | 0.350000378 | 0.402 | 0.136 | 1.01E-82 |
| **NOL7** | 4.02E-51 | 0.349044767 | 0.921 | 0.749 | 9.71E-47 |
| **TCF7** | 1.12E-48 | 0.348852974 | 0.316 | 0.132 | 2.7E-44 |
| **PPP1R16A** | 1.01E-41 | 0.348353354 | 0.884 | 0.731 | 2.45E-37 |
| **GLT8D1** | 7.41E-40 | 0.348302118 | 0.791 | 0.617 | 1.79E-35 |
| **WDR92** | 6.73E-44 | 0.347841324 | 0.799 | 0.605 | 1.62E-39 |
| **IFT172** | 1.39E-43 | 0.347551547 | 0.944 | 0.844 | 3.36E-39 |
| **ARF5** | 3.2E-40 | 0.347523406 | 0.453 | 0.257 | 7.74E-36 |
| **CLMN** | 9.61E-35 | 0.347104055 | 0.979 | 0.938 | 2.32E-30 |
| **TFPT** | 3.06E-24 | 0.346886113 | 0.538 | 0.389 | 7.39E-20 |
| **NAA38** | 6.65E-57 | 0.346884702 | 0.98 | 0.936 | 1.61E-52 |
| **KRT23** | 0.007802746 | 0.346428643 | 0.499 | 0.46 | 1 |
| **MYB** | 4.37E-42 | 0.346419964 | 0.768 | 0.571 | 1.05E-37 |
| **ATP6V1D** | 4.1E-53 | 0.345731422 | 0.978 | 0.89 | 9.91E-49 |
| **GABRB3** | 7.2E-96 | 0.3449855 | 0.239 | 0.046 | 1.74E-91 |
| **DNAH6** | 2.02E-46 | 0.344831171 | 0.982 | 0.924 | 4.87E-42 |
| **BX284668.5** | 7.25E-42 | 0.344764507 | 0.429 | 0.233 | 1.75E-37 |
| **CCDC69** | 3.4E-41 | 0.34474881 | 0.881 | 0.692 | 8.22E-37 |
| **MRPL47** | 1.11E-38 | 0.344599484 | 0.784 | 0.593 | 2.68E-34 |
| **DTHD1** | 4.76E-42 | 0.344572609 | 0.987 | 0.941 | 1.15E-37 |
| **EBNA1BP2** | 8.82E-46 | 0.344476352 | 0.899 | 0.715 | 2.13E-41 |
| **TMEM154** | 1.22E-37 | 0.344354862 | 0.888 | 0.752 | 2.94E-33 |
| **PLEKHA4** | 6.8E-51 | 0.344249066 | 0.359 | 0.158 | 1.64E-46 |
| **DNER** | 2.21E-46 | 0.344076522 | 0.707 | 0.466 | 5.33E-42 |
| **TEKT4** | 8.04E-40 | 0.343327387 | 0.594 | 0.383 | 1.94E-35 |
| **DYNLT1** | 1.13E-56 | 0.343115095 | 0.997 | 0.999 | 2.73E-52 |
| **AC025181.2** | 1.13E-38 | 0.341846253 | 0.658 | 0.461 | 2.73E-34 |
| **CLDN9** | 1.94E-72 | 0.341799214 | 0.263 | 0.073 | 4.68E-68 |
| **ANP32E** | 1.62E-40 | 0.341395288 | 0.707 | 0.494 | 3.91E-36 |
| **TRNAU1AP** | 7.35E-35 | 0.341112658 | 0.646 | 0.467 | 1.78E-30 |
| **AGBL2** | 1.43E-41 | 0.340923315 | 0.74 | 0.537 | 3.46E-37 |
| **IRF1** | 4.95E-31 | 0.340388618 | 0.834 | 0.681 | 1.2E-26 |
| **ATP5F1E** | 2.58E-89 | 0.339924121 | 0.994 | 0.999 | 6.22E-85 |
| **MED25** | 1.12E-44 | 0.339676736 | 0.87 | 0.72 | 2.69E-40 |

| | | | | | |
|---|---|---|---|---|---|
| **C6** | 4.8E-23 | 0.339436157 | 0.298 | 0.168 | 1.16E-18 |
| **ENO4** | 1.99E-51 | 0.339204826 | 0.63 | 0.375 | 4.82E-47 |
| **LYSMD2** | 1.95E-43 | 0.339202215 | 0.581 | 0.358 | 4.7E-39 |
| **AC096637.2** | 3.24E-48 | 0.338809168 | 0.638 | 0.389 | 7.82E-44 |
| **C16orf46** | 3.93E-48 | 0.338530425 | 0.762 | 0.54 | 9.5E-44 |
| **SOCS3** | 1.79E-18 | 0.338227882 | 0.434 | 0.304 | 4.33E-14 |
| **NBL1** | 3.83E-27 | 0.337404862 | 0.536 | 0.376 | 9.26E-23 |
| **AC008035.1** | 2.52E-85 | 0.337238486 | 0.4 | 0.136 | 6.09E-81 |
| **NDUFB5** | 1.56E-39 | 0.336795617 | 0.931 | 0.798 | 3.76E-35 |
| **HSPB11** | 5.12E-51 | 0.336424714 | 0.992 | 0.97 | 1.24E-46 |
| **KDM1B** | 6.58E-36 | 0.335991923 | 0.669 | 0.48 | 1.59E-31 |
| **ARHGAP18** | 1.83E-50 | 0.335778762 | 0.996 | 0.98 | 4.41E-46 |
| **SAT2** | 7.78E-34 | 0.335735565 | 0.856 | 0.678 | 1.88E-29 |
| **AC046134.2** | 5.75E-43 | 0.335283614 | 0.579 | 0.358 | 1.39E-38 |
| **MTIF3** | 3.96E-44 | 0.33527848 | 0.871 | 0.704 | 9.56E-40 |
| **FLACC1** | 5.82E-50 | 0.33454091 | 0.594 | 0.349 | 1.4E-45 |
| **SMIM19** | 8.2E-45 | 0.334175741 | 0.933 | 0.799 | 1.98E-40 |
| **EIF1B** | 2.63E-37 | 0.333976332 | 0.791 | 0.603 | 6.35E-33 |
| **CFAP410** | 2.31E-34 | 0.333943484 | 0.574 | 0.392 | 5.57E-30 |
| **WFDC6** | 5.52E-78 | 0.332807473 | 0.372 | 0.127 | 1.33E-73 |
| **C16orf71** | 7.32E-50 | 0.331470557 | 0.659 | 0.411 | 1.77E-45 |
| **MYO1D** | 1.43E-33 | 0.331458798 | 0.732 | 0.56 | 3.45E-29 |
| **ISCA2** | 7.31E-46 | 0.331300169 | 0.804 | 0.59 | 1.77E-41 |
| **LINC02345** | 1.78E-37 | 0.331242598 | 0.806 | 0.634 | 4.3E-33 |
| **PTPMT1** | 1.18E-40 | 0.330911665 | 0.794 | 0.606 | 2.85E-36 |
| **DDX3Y** | 4.33E-43 | 0.330688938 | 0.814 | 0.576 | 1.05E-38 |
| **MMP24OS** | 2.27E-30 | 0.330357342 | 0.9 | 0.775 | 5.47E-26 |
| **VMO1** | 2.33E-15 | 0.330235226 | 0.193 | 0.345 | 5.62E-11 |
| **CACNG6** | 1.72E-52 | 0.329501184 | 0.497 | 0.252 | 4.15E-48 |
| **PPP1R32** | 9.25E-57 | 0.329229972 | 0.462 | 0.219 | 2.23E-52 |
| **PHGDH** | 1.52E-39 | 0.32778487 | 0.412 | 0.22 | 3.67E-35 |
| **VRK3** | 2.47E-38 | 0.327407291 | 0.826 | 0.649 | 5.97E-34 |
| **AC007325.4** | 9.54E-38 | 0.327318974 | 0.551 | 0.346 | 2.31E-33 |
| **CDHR4** | 7.94E-45 | 0.326978673 | 0.949 | 0.796 | 1.92E-40 |
| **CLUAP1** | 4.34E-47 | 0.326685667 | 0.986 | 0.942 | 1.05E-42 |
| **MCAT** | 7.56E-48 | 0.326450344 | 0.632 | 0.392 | 1.83E-43 |
| **CROCC2** | 2.43E-102 | 0.325829906 | 0.27 | 0.056 | 5.86E-98 |
| **SYBU** | 2.36E-43 | 0.325776646 | 0.419 | 0.215 | 5.71E-39 |

| | | | | | |
|---|---|---|---|---|---|
| **CHIC2** | 5.19E-44 | 0.325507989 | 0.756 | 0.532 | 1.25E-39 |
| **EIF2S2** | 8.44E-50 | 0.325374308 | 0.98 | 0.911 | 2.04E-45 |
| **NUDCD2** | 2.36E-44 | 0.324741218 | 0.856 | 0.659 | 5.7E-40 |
| **SCG3** | 9.09E-54 | 0.324398471 | 0.194 | 0.052 | 2.2E-49 |
| **ADH1C** | 2.78E-11 | 0.324088778 | 0.381 | 0.288 | 6.7E-07 |
| **IFIT1** | 1.87E-33 | 0.324059613 | 0.348 | 0.182 | 4.52E-29 |
| **DENND6B** | 1.1E-35 | 0.323255419 | 0.807 | 0.627 | 2.65E-31 |
| **FAM166C** | 1.98E-59 | 0.322933791 | 0.407 | 0.175 | 4.78E-55 |
| **OCEL1** | 6.25E-39 | 0.322813136 | 0.788 | 0.598 | 1.51E-34 |
| **KIF3B** | 1.56E-31 | 0.322370275 | 0.942 | 0.855 | 3.78E-27 |
| **SCOC-AS1** | 7.98E-90 | 0.322313662 | 0.31 | 0.083 | 1.93E-85 |
| **INO80B** | 5.33E-46 | 0.322022529 | 0.557 | 0.327 | 1.29E-41 |
| **IDNK** | 1.78E-41 | 0.321169766 | 0.606 | 0.384 | 4.31E-37 |
| **HMGN5** | 2.72E-30 | 0.320716035 | 0.668 | 0.512 | 6.56E-26 |
| **MEAF6** | 2.76E-43 | 0.3201395 | 0.948 | 0.814 | 6.68E-39 |
| **GBP6** | 1E-35 | 0.3200846 | 0.422 | 0.24 | 2.42E-31 |
| **MGMT** | 4.2E-33 | 0.319730186 | 0.888 | 0.715 | 1.01E-28 |
| **ORAI2** | 9.26E-33 | 0.319630973 | 0.908 | 0.791 | 2.24E-28 |
| **SUGT1** | 1.57E-42 | 0.31928731 | 0.906 | 0.722 | 3.8E-38 |
| **NDUFA8** | 2.01E-31 | 0.317977939 | 0.92 | 0.803 | 4.85E-27 |
| **RCAN3AS** | 1.89E-61 | 0.317945225 | 0.468 | 0.211 | 4.58E-57 |
| **EFHC1** | 1.15E-47 | 0.317902018 | 0.999 | 0.991 | 2.78E-43 |
| **COX7A1** | 0.000516167 | 0.317760062 | 0.277 | 0.247 | 1 |
| **SNRPG** | 7.83E-53 | 0.317260406 | 0.981 | 0.921 | 1.89E-48 |
| **MRPS24** | 1.22E-42 | 0.317002915 | 0.944 | 0.799 | 2.94E-38 |
| **CRNDE** | 2.76E-51 | 0.316643104 | 0.994 | 0.971 | 6.67E-47 |
| **ABHD11** | 3.47E-37 | 0.316540288 | 0.716 | 0.517 | 8.37E-33 |
| **CFAP47** | 4.16E-35 | 0.316387803 | 0.847 | 0.7 | 1E-30 |
| **AGR2** | 0.000111725 | 0.316174766 | 0.844 | 0.885 | 1 |
| **TUSC2** | 1.01E-41 | 0.315826923 | 0.896 | 0.699 | 2.44E-37 |
| **HHLA2** | 9.84E-34 | 0.315495916 | 0.603 | 0.397 | 2.38E-29 |
| **AC130456.2** | 1.36E-45 | 0.315462156 | 0.562 | 0.323 | 3.27E-41 |
| **STYXL1** | 1.68E-42 | 0.315424575 | 0.851 | 0.632 | 4.06E-38 |
| **STIMATE** | 3.01E-31 | 0.315161894 | 0.511 | 0.337 | 7.26E-27 |
| **HIST1H2BD** | 1.99E-26 | 0.314926805 | 0.573 | 0.416 | 4.8E-22 |
| **TRPT1** | 3.9E-28 | 0.314686812 | 0.647 | 0.485 | 9.43E-24 |
| **NUDT7** | 4.19E-36 | 0.314182625 | 0.441 | 0.25 | 1.01E-31 |
| **AC105052.5** | 9.27E-39 | 0.31414668 | 0.606 | 0.387 | 2.24E-34 |

| | | | | | |
|---|---|---|---|---|---|
| **NHLRC4** | 1.13E-46 | 0.313095664 | 0.716 | 0.471 | 2.72E-42 |
| **TRNP1** | 7.91E-20 | 0.312978202 | 0.661 | 0.539 | 1.91E-15 |
| **CFAP57** | 2.52E-40 | 0.312448673 | 0.872 | 0.715 | 6.08E-36 |
| **MFSD2A** | 9.27E-26 | 0.311959761 | 0.671 | 0.515 | 2.24E-21 |
| **CALM1** | 9.28E-83 | 0.311838802 | 1 | 1 | 2.24E-78 |
| **SMIM26** | 5.05E-42 | 0.31178432 | 0.958 | 0.836 | 1.22E-37 |
| **B3GNT7** | 1.08E-26 | 0.311709668 | 0.798 | 0.663 | 2.61E-22 |
| **BBIP1** | 8.68E-39 | 0.311049629 | 0.911 | 0.758 | 2.1E-34 |
| **LEKR1** | 2.11E-40 | 0.310952778 | 0.633 | 0.414 | 5.09E-36 |
| **CST6** | 2.13E-13 | 0.310284021 | 0.642 | 0.495 | 5.15E-09 |
| **SUDS3** | 1.29E-40 | 0.309664297 | 0.909 | 0.759 | 3.1E-36 |
| **NDUFA6** | 9.6E-36 | 0.309483218 | 0.79 | 0.595 | 2.32E-31 |
| **FAM184A** | 2.53E-31 | 0.309063792 | 0.691 | 0.503 | 6.12E-27 |
| **MAK** | 1.02E-42 | 0.308900584 | 0.704 | 0.465 | 2.47E-38 |
| **CCP110** | 5.01E-32 | 0.308484968 | 0.933 | 0.819 | 1.21E-27 |
| **KRT8** | 1.36E-24 | 0.308448214 | 0.987 | 0.979 | 3.29E-20 |
| **GAS7** | 1.74E-37 | 0.308273365 | 0.539 | 0.329 | 4.21E-33 |
| **SPATA18** | 1.23E-37 | 0.30787568 | 0.983 | 0.931 | 2.98E-33 |
| **CDS1** | 1.23E-44 | 0.307740556 | 0.996 | 0.951 | 2.97E-40 |
| **CDKL2** | 1.38E-41 | 0.307684164 | 0.527 | 0.304 | 3.32E-37 |
| **BCAS3** | 2.19E-35 | 0.307395114 | 0.634 | 0.441 | 5.3E-31 |
| **FHAD1** | 2.25E-30 | 0.307235241 | 0.832 | 0.683 | 5.43E-26 |
| **TMEM254** | 1.9E-33 | 0.307183237 | 0.742 | 0.569 | 4.58E-29 |
| **TCTN2** | 4.28E-42 | 0.306733715 | 0.664 | 0.435 | 1.03E-37 |
| **POLD2** | 1.13E-40 | 0.306504753 | 0.884 | 0.709 | 2.73E-36 |
| **AL627171.2** | 6.36E-24 | 0.306229159 | 0.51 | 0.36 | 1.54E-19 |
| **PCBD1** | 3.19E-29 | 0.305993615 | 0.734 | 0.572 | 7.7E-25 |
| **MORN1** | 5.91E-42 | 0.305892906 | 0.714 | 0.482 | 1.43E-37 |
| **ABCC6** | 1.12E-48 | 0.305734092 | 0.563 | 0.32 | 2.7E-44 |
| **GLO1** | 1.73E-35 | 0.305285169 | 0.814 | 0.633 | 4.18E-31 |
| **CCDC121** | 2.77E-50 | 0.305004298 | 0.517 | 0.272 | 6.69E-46 |
| **MDM1** | 7.55E-37 | 0.304967042 | 0.761 | 0.57 | 1.82E-32 |
| **CDC26** | 1.07E-33 | 0.304266117 | 0.869 | 0.719 | 2.6E-29 |
| **TJP3** | 8.61E-38 | 0.303867545 | 0.902 | 0.733 | 2.08E-33 |
| **TMPRSS3** | 4.7E-45 | 0.303852302 | 0.442 | 0.226 | 1.13E-40 |
| **SF3B5** | 5.83E-28 | 0.303750503 | 0.891 | 0.766 | 1.41E-23 |
| **WDR13** | 2.18E-36 | 0.303567546 | 0.873 | 0.7 | 5.27E-32 |
| **F5** | 2.59E-45 | 0.303322898 | 0.283 | 0.113 | 6.26E-41 |

| | | | | | |
|---|---|---|---|---|---|
| **ADH6** | 2.5E-52 | 0.303313376 | 0.402 | 0.181 | 6.05E-48 |
| **THOC7** | 5.64E-40 | 0.302666541 | 0.878 | 0.717 | 1.36E-35 |
| **AC027237.3** | 3.81E-40 | 0.30261541 | 0.669 | 0.458 | 9.2E-36 |
| **JHY** | 2.18E-33 | 0.302355366 | 0.781 | 0.61 | 5.27E-29 |
| **NDUFAB1** | 2.72E-48 | 0.302352916 | 0.978 | 0.947 | 6.58E-44 |
| **LRRC36** | 8.46E-34 | 0.3019356 | 0.414 | 0.234 | 2.04E-29 |
| **DHX40** | 3.03E-33 | 0.301798607 | 0.864 | 0.727 | 7.31E-29 |
| **PITPNM1** | 3.5E-30 | 0.301540195 | 0.757 | 0.59 | 8.44E-26 |
| **EHD4** | 1.93E-26 | 0.301373742 | 0.636 | 0.467 | 4.66E-22 |
| **AC092802.1** | 1.38E-42 | 0.301111786 | 0.554 | 0.323 | 3.33E-38 |
| **C17orf82** | 5.06E-56 | 0.30072139 | 0.429 | 0.194 | 1.22E-51 |
| **ZNF584** | 4.47E-52 | 0.300705244 | 0.463 | 0.225 | 1.08E-47 |
| **DEUP1** | 1.01E-48 | 0.300704506 | 0.432 | 0.21 | 2.45E-44 |
| **ADIPOR1** | 1.18E-37 | 0.30061459 | 0.817 | 0.628 | 2.84E-33 |
| **ZC2HC1C** | 1.06E-47 | 0.300314424 | 0.638 | 0.383 | 2.56E-43 |
| **ARL 6.00** | 3.76E-40 | 0.299498361 | 0.706 | 0.488 | 9.08E-36 |
| **ID4** | 1.99E-06 | 0.29931225 | 0.549 | 0.51 | 0.048114609 |
| **ACAA2** | 4.58E-49 | 0.299275103 | 0.374 | 0.169 | 1.11E-44 |
| **ST6GALNAC6** | 1.13E-18 | 0.299078317 | 0.54 | 0.419 | 2.72E-14 |
| **RPS27** | 1.34E-64 | 0.299053287 | 0.993 | 0.999 | 3.24E-60 |
| **IRF2** | 2.26E-38 | 0.298920366 | 0.792 | 0.591 | 5.46E-34 |
| **IFT20** | 1.68E-35 | 0.298851651 | 0.878 | 0.723 | 4.05E-31 |
| **AC009119.1** | 3.77E-60 | 0.298615818 | 0.289 | 0.098 | 9.11E-56 |
| **COX7C** | 1.8E-58 | 0.298491857 | 0.994 | 0.998 | 4.35E-54 |
| **CCDC125** | 8.37E-35 | 0.298482762 | 0.86 | 0.684 | 2.02E-30 |
| **AP002008.4** | 6.98E-34 | 0.298404653 | 0.609 | 0.413 | 1.69E-29 |
| **SLF1** | 5.05E-38 | 0.298314796 | 0.538 | 0.328 | 1.22E-33 |
| **KATNB1** | 9.93E-35 | 0.298285198 | 0.678 | 0.48 | 2.4E-30 |
| **S100A9** | 5.97E-47 | 0.298268808 | 0.149 | 0.441 | 1.44E-42 |
| **SNHG9** | 5.11E-46 | 0.298077488 | 0.408 | 0.199 | 1.23E-41 |
| **TM7SF2** | 3.14E-26 | 0.29800625 | 0.82 | 0.681 | 7.58E-22 |
| **IFT52** | 6.58E-37 | 0.297973779 | 0.729 | 0.528 | 1.59E-32 |
| **TBCA** | 1.99E-58 | 0.297790219 | 0.992 | 0.988 | 4.81E-54 |
| **BANF1** | 1.11E-36 | 0.297576609 | 0.917 | 0.75 | 2.68E-32 |
| **IFT122** | 5.83E-40 | 0.297349209 | 0.777 | 0.574 | 1.41E-35 |
| **MRPL52** | 7.96E-35 | 0.297283846 | 0.849 | 0.673 | 1.92E-30 |
| **C10orf67** | 3.84E-41 | 0.297112814 | 0.57 | 0.342 | 9.29E-37 |
| **TSEN34** | 2.85E-32 | 0.297060596 | 0.714 | 0.53 | 6.88E-28 |

| | | | | | |
|---|---|---|---|---|---|
| **DZANK1** | 9.44E-40 | 0.296575835 | 0.671 | 0.448 | 2.28E-35 |
| **AZIN1** | 6.43E-35 | 0.296555357 | 0.968 | 0.905 | 1.55E-30 |
| **SCPEP1** | 6.21E-18 | 0.296491834 | 0.864 | 0.835 | 1.5E-13 |
| **TRMT10A** | 3.02E-34 | 0.296300876 | 0.533 | 0.342 | 7.3E-30 |
| **HSPA8** | 3.88E-47 | 0.296223023 | 0.997 | 0.998 | 9.38E-43 |
| **AC044849.1** | 3.02E-49 | 0.296020003 | 0.526 | 0.278 | 7.31E-45 |
| **BAD** | 2.76E-35 | 0.296015078 | 0.753 | 0.538 | 6.67E-31 |
| **SEC14L4** | 1.39E-38 | 0.295906996 | 0.526 | 0.313 | 3.35E-34 |
| **MYLK3** | 1.54E-40 | 0.29586639 | 0.153 | 0.043 | 3.73E-36 |
| **ZBBX** | 6.38E-38 | 0.295733161 | 0.981 | 0.894 | 1.54E-33 |
| **ATP5PD** | 1.49E-40 | 0.295670722 | 0.939 | 0.846 | 3.61E-36 |
| **ZCCHC10** | 7.06E-38 | 0.295642294 | 0.618 | 0.408 | 1.71E-33 |
| **C11orf49** | 2.29E-37 | 0.295311742 | 0.779 | 0.581 | 5.53E-33 |
| **PSMB9** | 8.28E-35 | 0.295218883 | 0.683 | 0.478 | 2E-30 |
| **AC010255.3** | 6.29E-51 | 0.295108477 | 0.351 | 0.15 | 1.52E-46 |
| **SMAP2** | 4.8E-32 | 0.295097576 | 0.817 | 0.658 | 1.16E-27 |
| **ERICH6-AS1** | 6.63E-36 | 0.294918107 | 0.713 | 0.499 | 1.6E-31 |
| **FAM3D** | 1.61E-08 | 0.294785244 | 0.244 | 0.175 | 0.000387906 |
| **RBM38** | 1.79E-24 | 0.294646727 | 0.636 | 0.489 | 4.32E-20 |
| **AKNA** | 1.47E-29 | 0.29423283 | 0.772 | 0.6 | 3.55E-25 |
| **METRN** | 9.93E-09 | 0.293952445 | 0.526 | 0.456 | 0.000239892 |
| **RRAGA** | 4.23E-36 | 0.293810892 | 0.899 | 0.743 | 1.02E-31 |
| **PDLIM4** | 3.71E-20 | 0.293777401 | 0.753 | 0.634 | 8.95E-16 |
| **DYNLRB1** | 2.02E-42 | 0.293770969 | 0.988 | 0.948 | 4.88E-38 |
| **PYCARD** | 2.97E-24 | 0.293412555 | 0.834 | 0.688 | 7.17E-20 |
| **AK9** | 6.66E-33 | 0.293331424 | 0.961 | 0.872 | 1.61E-28 |
| **SLC4A8** | 8.03E-24 | 0.293215921 | 0.681 | 0.52 | 1.94E-19 |
| **LYPLA2** | 1.9E-31 | 0.292987707 | 0.818 | 0.637 | 4.59E-27 |
| **ODF2L** | 1.29E-34 | 0.292711991 | 0.994 | 0.975 | 3.12E-30 |
| **MAGIX** | 4.39E-37 | 0.292663667 | 0.611 | 0.403 | 1.06E-32 |
| **ARMC9** | 4.63E-35 | 0.29216228 | 0.767 | 0.586 | 1.12E-30 |
| **HAGH** | 1.22E-37 | 0.292027568 | 0.908 | 0.741 | 2.95E-33 |
| **SNAPC5** | 2.49E-32 | 0.29179218 | 0.629 | 0.442 | 6.01E-28 |
| **BBS12** | 2.54E-48 | 0.291624886 | 0.53 | 0.288 | 6.12E-44 |
| **DGCR6L** | 3.91E-33 | 0.291548347 | 0.631 | 0.432 | 9.44E-29 |
| **PARD6G-AS1** | 2.2E-40 | 0.291529757 | 0.407 | 0.209 | 5.32E-36 |
| **CKLF** | 1.5E-20 | 0.291316932 | 0.617 | 0.486 | 3.63E-16 |
| **FAM161A** | 2.26E-28 | 0.291266914 | 0.757 | 0.584 | 5.45E-24 |

| | | | | | |
|---|---|---|---|---|---|
| **DPM3** | 5.82E-31 | 0.291066149 | 0.82 | 0.658 | 1.41E-26 |
| **LZTFL1** | 6.02E-37 | 0.291004195 | 0.946 | 0.835 | 1.45E-32 |
| **UBE2V1** | 2.75E-40 | 0.290900493 | 0.924 | 0.776 | 6.64E-36 |
| **ERCC3** | 3.99E-22 | 0.290872081 | 0.573 | 0.429 | 9.64E-18 |
| **EML1** | 5.55E-35 | 0.290688713 | 0.684 | 0.484 | 1.34E-30 |
| **USP2** | 5.51E-33 | 0.290468828 | 0.77 | 0.579 | 1.33E-28 |
| **SNHG10** | 1.36E-36 | 0.290420597 | 0.538 | 0.33 | 3.29E-32 |
| **PRXL2C** | 1.48E-22 | 0.290294993 | 0.607 | 0.469 | 3.57E-18 |
| **IRX3** | 4.58E-20 | 0.290291755 | 0.818 | 0.733 | 1.11E-15 |
| **MAFG-DT** | 1.88E-40 | 0.290153517 | 0.546 | 0.322 | 4.54E-36 |
| **ENOSF1** | 5.43E-28 | 0.290065106 | 0.726 | 0.569 | 1.31E-23 |
| **SLC16A12** | 3.38E-43 | 0.289993068 | 0.348 | 0.159 | 8.16E-39 |
| **IPO11** | 4.13E-35 | 0.289906736 | 0.443 | 0.251 | 9.97E-31 |
| **CEP19** | 9.26E-40 | 0.289798779 | 0.637 | 0.401 | 2.24E-35 |
| **RFXANK** | 1.37E-26 | 0.289470189 | 0.552 | 0.386 | 3.32E-22 |
| **PPP1R9A** | 2.37E-43 | 0.289340906 | 0.356 | 0.166 | 5.73E-39 |
| **LACTB2** | 4.66E-32 | 0.289206556 | 0.474 | 0.286 | 1.12E-27 |
| **CCDC28A** | 2.49E-27 | 0.289174573 | 0.841 | 0.705 | 6E-23 |
| **PRR29** | 1.95E-33 | 0.288328136 | 0.849 | 0.703 | 4.7E-29 |
| **ST8SIA1** | 2.59E-34 | 0.288101598 | 0.417 | 0.233 | 6.26E-30 |
| **NSRP1** | 1.96E-31 | 0.288011963 | 0.726 | 0.542 | 4.74E-27 |
| **SEC14L1** | 1.23E-35 | 0.287961644 | 0.94 | 0.803 | 2.97E-31 |
| **VNN 2.00** | 1.3E-70 | 0.28772803 | 0.277 | 0.081 | 3.14E-66 |
| **HMGB3** | 8.14E-08 | 0.287273917 | 0.396 | 0.329 | 0.001966837 |
| **JUNB** | 6.83E-07 | 0.287255742 | 0.793 | 0.863 | 0.01648626 |
| **CATSPERD** | 7.89E-32 | 0.287010852 | 0.331 | 0.169 | 1.91E-27 |
| **UNC119B** | 2.29E-31 | 0.286552155 | 0.806 | 0.656 | 5.54E-27 |
| **HMGB2** | 1.39E-29 | 0.286442485 | 0.557 | 0.376 | 3.35E-25 |
| **SLC34A2** | 0.000184576 | 0.286352489 | 0.428 | 0.385 | 1 |
| **MGLL** | 9.69E-22 | 0.286192445 | 0.618 | 0.466 | 2.34E-17 |
| **RPL37** | 2.06E-51 | 0.285755264 | 0.996 | 0.999 | 4.97E-47 |
| **AC036176.1** | 4.67E-41 | 0.28515501 | 0.484 | 0.272 | 1.13E-36 |
| **NME9** | 3E-38 | 0.284962621 | 0.599 | 0.377 | 7.24E-34 |
| **WDR66** | 7.64E-33 | 0.284913938 | 0.992 | 0.942 | 1.85E-28 |
| **GBP1** | 2.49E-14 | 0.284756166 | 0.409 | 0.295 | 6.01E-10 |
| **ATP5MF** | 1.32E-45 | 0.284742059 | 0.984 | 0.966 | 3.19E-41 |
| **GRAMD2B** | 3.85E-06 | 0.284401405 | 0.53 | 0.478 | 0.093101459 |
| **CGN** | 6.67E-31 | 0.284124828 | 0.813 | 0.665 | 1.61E-26 |

| | | | | | |
|---|---|---|---|---|---|
| **NOA1** | 2.37E-27 | 0.284044642 | 0.541 | 0.373 | 5.72E-23 |
| **MDH1B** | 1.95E-33 | 0.2838052 | 0.812 | 0.631 | 4.71E-29 |
| **DBNDD2** | 1.51E-31 | 0.283583534 | 0.663 | 0.469 | 3.64E-27 |
| **TRMT1L** | 1.12E-27 | 0.283487808 | 0.768 | 0.627 | 2.71E-23 |
| **RNF32** | 4.98E-46 | 0.283327258 | 0.49 | 0.26 | 1.2E-41 |
| **LMO7-AS1** | 3.65E-38 | 0.28317724 | 0.458 | 0.255 | 8.81E-34 |
| **PYCR2** | 7.69E-34 | 0.282946885 | 0.704 | 0.516 | 1.86E-29 |
| **UQCRQ** | 1.01E-43 | 0.282939304 | 0.992 | 0.972 | 2.43E-39 |
| **SMIM4** | 5.91E-35 | 0.282916984 | 0.603 | 0.393 | 1.43E-30 |
| **RCAN3** | 5.22E-36 | 0.282808504 | 0.959 | 0.846 | 1.26E-31 |
| **C11orf52** | 5.17E-29 | 0.28268412 | 0.626 | 0.457 | 1.25E-24 |
| **INTS10** | 2.58E-29 | 0.282557851 | 0.63 | 0.445 | 6.23E-25 |
| **CFI** | 2.37E-15 | 0.282255962 | 0.288 | 0.189 | 5.72E-11 |
| **PRR15** | 1.49E-27 | 0.282124904 | 0.933 | 0.863 | 3.59E-23 |
| **CWH43** | 1.06E-26 | 0.282081776 | 0.466 | 0.295 | 2.57E-22 |
| **GOLPH3L** | 7.48E-33 | 0.281925688 | 0.822 | 0.652 | 1.81E-28 |
| **SIX3-AS1** | 8.88E-24 | 0.281579026 | 0.449 | 0.294 | 2.15E-19 |
| **AC133552.5** | 1.85E-30 | 0.281312234 | 0.473 | 0.296 | 4.48E-26 |
| **AKR1A1** | 2.19E-10 | 0.281043841 | 0.811 | 0.784 | 5.3E-06 |
| **MRPL57** | 5.74E-38 | 0.281022078 | 0.973 | 0.899 | 1.39E-33 |
| **TXLNB** | 3.05E-40 | 0.280966383 | 0.552 | 0.32 | 7.36E-36 |
| **CFAP97** | 5.49E-32 | 0.280870659 | 0.842 | 0.661 | 1.33E-27 |
| **ACAP1** | 5.41E-21 | 0.280566296 | 0.54 | 0.398 | 1.31E-16 |
| **NEK10** | 1.09E-27 | 0.280434881 | 0.973 | 0.922 | 2.63E-23 |
| **AZIN1-AS1** | 1.34E-41 | 0.280407299 | 0.552 | 0.32 | 3.24E-37 |
| **AMZ2** | 2.27E-31 | 0.279436522 | 0.673 | 0.49 | 5.49E-27 |
| **MT-CYB** | 3.55E-12 | 0.279146429 | 1 | 1 | 8.58E-08 |
| **CFAP74** | 3.11E-30 | 0.279096244 | 0.877 | 0.709 | 7.51E-26 |
| **PECR** | 3E-33 | 0.279094722 | 0.529 | 0.336 | 7.24E-29 |
| **FBXO36** | 1.3E-35 | 0.279037259 | 0.787 | 0.591 | 3.13E-31 |
| **AGBL5** | 8.93E-24 | 0.279023507 | 0.618 | 0.468 | 2.16E-19 |
| **TXN** | 3.27E-18 | 0.278708045 | 0.997 | 1 | 7.9E-14 |
| **MT-ND3** | 8.39E-12 | 0.278643525 | 0.998 | 1 | 2.03E-07 |
| **MRPL33** | 1.22E-39 | 0.278415353 | 0.946 | 0.811 | 2.96E-35 |
| **ABHD12B** | 1.17E-29 | 0.278310768 | 0.453 | 0.277 | 2.83E-25 |
| **C6orf132** | 1.5E-22 | 0.278276449 | 0.926 | 0.841 | 3.63E-18 |
| **MED31** | 3.47E-27 | 0.277835073 | 0.88 | 0.746 | 8.39E-23 |
| **MKS1** | 1.31E-42 | 0.277441027 | 0.551 | 0.318 | 3.16E-38 |

| | | | | | |
|---|---|---|---|---|---|
| **UQCR11** | 1.13E-55 | 0.277184609 | 0.991 | 0.995 | 2.73E-51 |
| **RTP4** | 7.6E-30 | 0.277166856 | 0.489 | 0.309 | 1.84E-25 |
| **C8orf76** | 1.85E-26 | 0.277073249 | 0.803 | 0.665 | 4.46E-22 |
| **ASL** | 6.01E-20 | 0.276838197 | 0.459 | 0.319 | 1.45E-15 |
| **RPL31** | 4.24E-37 | 0.276625796 | 0.962 | 0.954 | 1.02E-32 |
| **AL163051.1** | 2.86E-38 | 0.276411443 | 0.582 | 0.36 | 6.91E-34 |
| **SH3YL1** | 2.68E-35 | 0.27625228 | 0.793 | 0.575 | 6.48E-31 |
| **JUND** | 3.31E-30 | 0.276097142 | 0.964 | 0.915 | 8E-26 |
| **CCDC157** | 2.96E-34 | 0.276077078 | 0.684 | 0.472 | 7.14E-30 |
| **SERPINB1** | 1.57E-24 | 0.276069222 | 0.958 | 0.921 | 3.78E-20 |
| **STK16** | 2.16E-30 | 0.275903952 | 0.673 | 0.49 | 5.22E-26 |
| **ULK4** | 3.11E-31 | 0.275639192 | 0.738 | 0.575 | 7.5E-27 |
| **EIF4EBP3** | 1.36E-27 | 0.275172294 | 0.694 | 0.513 | 3.29E-23 |
| **DAD1** | 2.43E-36 | 0.275164847 | 0.98 | 0.912 | 5.87E-32 |
| **LRRC27** | 1.3E-35 | 0.2751317 | 0.659 | 0.444 | 3.14E-31 |
| **TRIM31** | 6.14E-34 | 0.274912625 | 0.251 | 0.109 | 1.48E-29 |
| **DIAPH2** | 3.46E-06 | 0.274676171 | 0.594 | 0.588 | 0.083464922 |
| **DNAJB4** | 2.62E-21 | 0.274646538 | 0.583 | 0.44 | 6.33E-17 |
| **ZC3H6** | 3.09E-29 | 0.27462854 | 0.75 | 0.583 | 7.47E-25 |
| **LRRIQ3** | 5.09E-34 | 0.274532045 | 0.628 | 0.421 | 1.23E-29 |
| **PDCD5** | 7.3E-31 | 0.274511899 | 0.773 | 0.582 | 1.76E-26 |
| **CEP89** | 2.43E-27 | 0.274106101 | 0.702 | 0.537 | 5.87E-23 |
| **NAP1L4** | 5.55E-18 | 0.27389471 | 0.85 | 0.744 | 1.34E-13 |
| **ALDH3A1** | 0.391463817 | 0.273832199 | 0.924 | 0.987 | 1 |
| **PSMA5** | 8.61E-34 | 0.273616507 | 0.917 | 0.773 | 2.08E-29 |
| **HIST2H2AA4** | 3.22E-121 | 0.273509563 | 0.194 | 0.021 | 7.78E-117 |
| **NDUFA7** | 9.49E-35 | 0.273418175 | 0.962 | 0.871 | 2.29E-30 |
| **SNW1** | 1.1E-33 | 0.273295438 | 0.827 | 0.634 | 2.66E-29 |
| **SNX2** | 4.48E-31 | 0.273295438 | 0.916 | 0.776 | 1.08E-26 |
| **MRPS6** | 5.7E-22 | 0.27295531 | 0.733 | 0.58 | 1.38E-17 |
| **NKAP** | 9.25E-27 | 0.271991086 | 0.677 | 0.512 | 2.23E-22 |
| **ABO** | 1.98E-22 | 0.271601685 | 0.508 | 0.359 | 4.77E-18 |
| **BAMBI** | 6.63E-20 | 0.271590062 | 0.454 | 0.317 | 1.6E-15 |
| **NARS** | 2.69E-32 | 0.271436056 | 0.951 | 0.847 | 6.5E-28 |
| **RAB11FIP1** | 5.36E-08 | 0.271259823 | 0.981 | 0.944 | 0.001294902 |
| **GEMIN7-AS1** | 1.38E-39 | 0.271003914 | 0.571 | 0.341 | 3.34E-35 |
| **APOL4** | 2.19E-77 | 0.270821937 | 0.284 | 0.078 | 5.3E-73 |
| **SMDT1** | 2.56E-31 | 0.270813176 | 0.968 | 0.873 | 6.19E-27 |

| | | | | | |
|---|---|---|---|---|---|
| **PDCD6** | 4.16E-38 | 0.270527643 | 0.97 | 0.893 | 1E-33 |
| **RWDD4** | 1.94E-28 | 0.270209458 | 0.592 | 0.416 | 4.68E-24 |
| **KIF27** | 1.09E-27 | 0.269782532 | 0.868 | 0.746 | 2.62E-23 |
| **PSMB8** | 3.48E-25 | 0.269546111 | 0.791 | 0.634 | 8.41E-21 |
| **ALDH1L1** | 2.04E-11 | 0.269435032 | 0.288 | 0.199 | 4.93E-07 |
| **KIAA0895** | 1.77E-32 | 0.269398158 | 0.607 | 0.405 | 4.27E-28 |
| **ECH1** | 1.65E-19 | 0.269180098 | 0.768 | 0.641 | 3.98E-15 |
| **TMEM258** | 1.08E-41 | 0.268860114 | 0.983 | 0.954 | 2.6E-37 |
| **KCNE1** | 2.38E-26 | 0.268804692 | 0.731 | 0.57 | 5.74E-22 |
| **CIAO2B** | 2.63E-29 | 0.268639454 | 0.952 | 0.847 | 6.35E-25 |
| **SAPCD1-AS1** | 3.91E-76 | 0.26798001 | 0.231 | 0.054 | 9.44E-72 |
| **KIF2A** | 4.21E-31 | 0.267908008 | 0.88 | 0.715 | 1.02E-26 |
| **NAT14** | 7.86E-30 | 0.267744668 | 0.587 | 0.398 | 1.9E-25 |
| **HDGF** | 3.76E-28 | 0.267571497 | 0.961 | 0.905 | 9.09E-24 |
| **THYN1** | 1.91E-30 | 0.267524492 | 0.651 | 0.46 | 4.62E-26 |
| **TLCD1** | 5.97E-30 | 0.26751173 | 0.468 | 0.285 | 1.44E-25 |
| **CNFN** | 7.37E-34 | 0.267215451 | 0.374 | 0.199 | 1.78E-29 |
| **NCS1** | 7.35E-26 | 0.266999548 | 0.621 | 0.456 | 1.78E-21 |
| **BDH2** | 1.87E-30 | 0.266923949 | 0.712 | 0.524 | 4.51E-26 |
| **DPP7** | 2.32E-29 | 0.266826063 | 0.844 | 0.655 | 5.6E-25 |
| **HSPA1B** | 7.14E-31 | 0.266656131 | 0.671 | 0.465 | 1.73E-26 |
| **ACTR6** | 8.91E-31 | 0.266592879 | 0.767 | 0.585 | 2.15E-26 |
| **ORMDL2** | 1.35E-29 | 0.266503224 | 0.804 | 0.62 | 3.27E-25 |
| **MRPL40** | 2.05E-27 | 0.266415626 | 0.948 | 0.844 | 4.96E-23 |
| **AL354953.1** | 3.05E-43 | 0.266338637 | 0.441 | 0.227 | 7.36E-39 |
| **VCAN** | 3.67E-24 | 0.266118813 | 0.174 | 0.074 | 8.87E-20 |
| **OTUD6B-AS1** | 9.08E-29 | 0.265869915 | 0.901 | 0.749 | 2.19E-24 |
| **VWA3B** | 7.76E-28 | 0.265851059 | 0.93 | 0.824 | 1.87E-23 |
| **RALGPS2** | 5.76E-27 | 0.265825844 | 0.801 | 0.638 | 1.39E-22 |
| **SYS1** | 6.27E-29 | 0.265166198 | 0.738 | 0.56 | 1.51E-24 |
| **FAM149A** | 3.21E-43 | 0.265095935 | 0.497 | 0.269 | 7.76E-39 |
| **CCDC60** | 1.11E-30 | 0.264956738 | 0.74 | 0.575 | 2.68E-26 |
| **TRADD** | 4.01E-29 | 0.264898453 | 0.674 | 0.495 | 9.69E-25 |
| **AC034139.1** | 3.72E-42 | 0.264800833 | 0.359 | 0.169 | 8.98E-38 |
| **AC005962.2** | 2.07E-57 | 0.264789468 | 0.316 | 0.116 | 5.01E-53 |
| **AC099518.4** | 1.88E-60 | 0.264508708 | 0.338 | 0.125 | 4.55E-56 |
| **RNF5** | 1E-30 | 0.26446744 | 0.701 | 0.504 | 2.42E-26 |
| **SNF8** | 4.9E-30 | 0.264310936 | 0.803 | 0.614 | 1.18E-25 |

| | | | | | |
|---|---|---|---|---|---|
| **NANS** | 1.14E-26 | 0.264296043 | 0.88 | 0.755 | 2.74E-22 |
| **TMEM219** | 7.39E-30 | 0.263994424 | 0.962 | 0.863 | 1.79E-25 |
| **MRPL21** | 8.76E-32 | 0.263919589 | 0.757 | 0.551 | 2.12E-27 |
| **ATP5MPL** | 3.97E-41 | 0.263553057 | 0.987 | 0.968 | 9.59E-37 |
| **PAIP2B** | 5.77E-35 | 0.263502597 | 0.517 | 0.312 | 1.39E-30 |
| **CRIP2** | 4.09E-31 | 0.263235692 | 0.882 | 0.725 | 9.89E-27 |
| **CEP112** | 1.83E-41 | 0.263184315 | 0.443 | 0.232 | 4.43E-37 |
| **FXR1** | 1.24E-32 | 0.262891611 | 0.892 | 0.745 | 2.99E-28 |
| **POLD3** | 3.8E-25 | 0.262847343 | 0.679 | 0.53 | 9.18E-21 |
| **BRD3OS** | 2.54E-30 | 0.262753813 | 0.664 | 0.479 | 6.13E-26 |
| **ATP5MD** | 7.18E-41 | 0.262726401 | 0.981 | 0.956 | 1.74E-36 |
| **RILPL2** | 5.95E-27 | 0.262513806 | 0.76 | 0.588 | 1.44E-22 |
| **MCRIP2** | 7.02E-28 | 0.262464037 | 0.543 | 0.363 | 1.7E-23 |
| **AGTRAP** | 8.19E-31 | 0.262228741 | 0.704 | 0.511 | 1.98E-26 |
| **IPO4** | 5.6E-31 | 0.262118006 | 0.554 | 0.362 | 1.35E-26 |
| **CHCHD6** | 1.85E-25 | 0.26168649 | 0.667 | 0.5 | 4.47E-21 |
| **PSMB5** | 3.34E-30 | 0.261649108 | 0.976 | 0.919 | 8.06E-26 |
| **COPS9** | 3.27E-32 | 0.261348147 | 0.948 | 0.844 | 7.9E-28 |
| **CCDC191** | 3.25E-19 | 0.261332073 | 0.849 | 0.756 | 7.85E-15 |
| **UBL5** | 9.84E-53 | 0.260929892 | 0.991 | 0.995 | 2.38E-48 |
| **TRAPPC2L** | 1.9E-23 | 0.260848792 | 0.75 | 0.609 | 4.58E-19 |
| **MTX2** | 1.3E-32 | 0.260663126 | 0.741 | 0.532 | 3.14E-28 |
| **IL20RA** | 9.97E-25 | 0.260545235 | 0.606 | 0.443 | 2.41E-20 |
| **FAM81A** | 6.97E-35 | 0.260235058 | 0.577 | 0.364 | 1.68E-30 |
| **ID2** | 0.092346376 | 0.259853834 | 0.489 | 0.505 | 1 |
| **SVIL** | 9.15E-25 | 0.259644923 | 0.496 | 0.334 | 2.21E-20 |
| **WDR49** | 1.31E-22 | 0.259606332 | 0.85 | 0.712 | 3.17E-18 |
| **KTN1** | 1.47E-47 | 0.259400949 | 0.999 | 0.999 | 3.56E-43 |
| **GET3** | 2.09E-31 | 0.25875932 | 0.723 | 0.505 | 5.05E-27 |
| **CYB5D1** | 6.32E-29 | 0.258615919 | 0.833 | 0.685 | 1.53E-24 |
| **PSIP1** | 6.38E-28 | 0.258217186 | 0.734 | 0.555 | 1.54E-23 |
| **PIBF1** | 7.14E-28 | 0.258003673 | 0.794 | 0.631 | 1.73E-23 |
| **IRAK1BP1** | 6.71E-28 | 0.257961077 | 0.573 | 0.394 | 1.62E-23 |
| **LRRC43** | 1.27E-40 | 0.25770397 | 0.546 | 0.313 | 3.07E-36 |
| **DLG5-AS1** | 1.23E-35 | 0.257676438 | 0.359 | 0.182 | 2.97E-31 |
| **TIMM10B** | 2.82E-19 | 0.257258722 | 0.473 | 0.334 | 6.81E-15 |
| **GMPR2** | 4.84E-28 | 0.257219097 | 0.689 | 0.511 | 1.17E-23 |
| **FAM182A** | 5.34E-39 | 0.256985977 | 0.484 | 0.269 | 1.29E-34 |

| | | | | | |
|---|---|---|---|---|---|
| **MAPK15** | 4.76E-19 | 0.256963211 | 0.829 | 0.698 | 1.15E-14 |
| **TTC21A** | 2.68E-28 | 0.256929071 | 0.848 | 0.674 | 6.46E-24 |
| **TLE4** | 3.49E-23 | 0.256761007 | 0.739 | 0.577 | 8.44E-19 |
| **HRK** | 3.47E-29 | 0.256694378 | 0.43 | 0.256 | 8.39E-25 |
| **CCDC184** | 1.82E-53 | 0.256403659 | 0.361 | 0.149 | 4.4E-49 |
| **LINC00326** | 7.87E-26 | 0.256306765 | 0.411 | 0.251 | 1.9E-21 |
| **DNMT1** | 1.52E-27 | 0.256219291 | 0.694 | 0.503 | 3.66E-23 |
| **NDUFB4** | 9.15E-34 | 0.256140599 | 0.96 | 0.892 | 2.21E-29 |
| **LINC02754** | 3.26E-41 | 0.256131714 | 0.328 | 0.146 | 7.87E-37 |
| **APH1B** | 4E-31 | 0.256112176 | 0.45 | 0.265 | 9.67E-27 |
| **EPCAM** | 1.57E-25 | 0.255836673 | 0.943 | 0.837 | 3.8E-21 |
| **SYAP1** | 2.29E-30 | 0.255833649 | 0.963 | 0.889 | 5.52E-26 |
| **NRAV** | 5.82E-31 | 0.255825442 | 0.832 | 0.655 | 1.41E-26 |
| **NMNAT2** | 2.96E-74 | 0.255382239 | 0.24 | 0.059 | 7.16E-70 |
| **SHROOM3** | 9.11E-19 | 0.255361069 | 0.894 | 0.794 | 2.2E-14 |
| **TSPAN6** | 9.88E-28 | 0.255324274 | 0.986 | 0.945 | 2.39E-23 |
| **REEP1** | 4.5E-60 | 0.255052628 | 0.26 | 0.081 | 1.09E-55 |
| **NSUN7** | 1.09E-25 | 0.255050159 | 0.801 | 0.671 | 2.63E-21 |
| **CYB5R2** | 3.35E-28 | 0.254811648 | 0.37 | 0.211 | 8.08E-24 |
| **PARP14** | 1.82E-18 | 0.254771466 | 0.826 | 0.693 | 4.4E-14 |
| **SEM1** | 1.98E-38 | 0.254752797 | 0.994 | 0.991 | 4.79E-34 |
| **LAMP3** | 8.22E-48 | 0.254660492 | 0.312 | 0.126 | 1.99E-43 |
| **ETFB** | 2.16E-26 | 0.254593549 | 0.876 | 0.725 | 5.21E-22 |
| **HPF1** | 6.97E-28 | 0.254027375 | 0.518 | 0.342 | 1.68E-23 |
| **RPL26L1** | 3.3E-30 | 0.253968273 | 0.827 | 0.639 | 7.97E-26 |
| **CCT2** | 2.18E-32 | 0.253785276 | 0.941 | 0.822 | 5.28E-28 |
| **FCF1** | 2.73E-25 | 0.253751788 | 0.643 | 0.47 | 6.6E-21 |
| **RSPH3** | 3.41E-23 | 0.253107743 | 0.964 | 0.898 | 8.23E-19 |
| **FOS** | 0.101919608 | 0.253014477 | 0.706 | 0.796 | 1 |
| **CLDN7** | 1.91E-23 | 0.252530649 | 0.99 | 0.961 | 4.61E-19 |
| **HERC5** | 1.54E-42 | 0.252251342 | 0.237 | 0.087 | 3.72E-38 |
| **FKBP8** | 4.2E-26 | 0.252138995 | 0.758 | 0.584 | 1.01E-21 |
| **ALG1L** | 4.98E-17 | 0.251945339 | 0.264 | 0.157 | 1.2E-12 |
| **NDUFV2** | 4.45E-34 | 0.251680175 | 0.982 | 0.939 | 1.08E-29 |
| **NUDT14** | 2E-26 | 0.251474835 | 0.871 | 0.747 | 4.83E-22 |
| **ARMH1** | 1.32E-32 | 0.251326923 | 0.592 | 0.38 | 3.19E-28 |
| **ERICH5** | 8.29E-19 | 0.251297351 | 0.797 | 0.687 | 2E-14 |
| **HIPK1** | 2.16E-29 | 0.250985897 | 0.984 | 0.937 | 5.21E-25 |

| | | | | | |
|---|---|---|---|---|---|
| **ICK** | 3.04E-24 | 0.250943599 | 0.687 | 0.516 | 7.34E-20 |
| **GET1** | 2.61E-26 | 0.250836896 | 0.933 | 0.817 | 6.31E-22 |
| **CCDC89** | 2.14E-29 | 0.250685644 | 0.329 | 0.172 | 5.17E-25 |
| **C1QBP** | 9E-25 | 0.250431026 | 0.873 | 0.735 | 2.17E-20 |
| **KCNH3** | 1.19E-31 | 0.250258802 | 0.469 | 0.281 | 2.87E-27 |
| **MT-ND4L** | 1.86E-14 | 0.250250452 | 0.926 | 0.928 | 4.49E-10 |
| **ABRAXAS1** | 1.7E-24 | 0.250186546 | 0.598 | 0.428 | 4.11E-20 |
| **MX2** | 1.37E-29 | 0.250072687 | 0.291 | 0.144 | 3.31E-25 |
| **ERGIC3** | 2.42E-28 | 0.24995265 | 0.97 | 0.909 | 5.85E-24 |
| **TRMT112** | 1.84E-27 | 0.249944908 | 0.948 | 0.867 | 4.44E-23 |
| **NDUFS3** | 6.78E-22 | 0.249944562 | 0.808 | 0.666 | 1.64E-17 |
| **KBTBD4** | 1.19E-22 | 0.249906485 | 0.588 | 0.435 | 2.88E-18 |
| **AC007114.1** | 1.7E-36 | 0.249856173 | 0.413 | 0.221 | 4.11E-32 |
| **PSMD4** | 1.4E-29 | 0.249687723 | 0.914 | 0.798 | 3.37E-25 |
| **UMODL1-AS1** | 3.83E-38 | 0.249675835 | 0.378 | 0.189 | 9.25E-34 |
| **IFITM2** | 1.29E-07 | 0.249376216 | 0.579 | 0.516 | 0.003107271 |
| **TUBGCP2** | 1.59E-33 | 0.2492965 | 0.971 | 0.891 | 3.85E-29 |
| **HBB** | 5.3E-17 | 0.249057777 | 0.153 | 0.071 | 1.28E-12 |
| **C11orf58** | 1.68E-24 | 0.248832219 | 0.954 | 0.895 | 4.06E-20 |
| **DNTTIP1** | 5.48E-29 | 0.248773039 | 0.654 | 0.458 | 1.32E-24 |
| **TGS1** | 4.57E-21 | 0.248690985 | 0.566 | 0.412 | 1.1E-16 |
| **COMT** | 1.4E-24 | 0.248258953 | 0.882 | 0.808 | 3.37E-20 |
| **EIF1AY** | 2.28E-34 | 0.248022433 | 0.646 | 0.412 | 5.51E-30 |
| **ATF3** | 1.82E-14 | 0.247922227 | 0.577 | 0.458 | 4.4E-10 |
| **LINC02832** | 1.13E-33 | 0.247773571 | 0.3 | 0.141 | 2.72E-29 |
| **EPB41L4B** | 5.3E-26 | 0.247747298 | 0.76 | 0.602 | 1.28E-21 |
| **DNAAF5** | 7.86E-27 | 0.24767192 | 0.628 | 0.453 | 1.9E-22 |
| **C4orf3** | 2.58E-37 | 0.247667897 | 0.99 | 0.984 | 6.23E-33 |
| **SCCPDH** | 4.68E-29 | 0.247570187 | 0.828 | 0.658 | 1.13E-24 |
| **RPL41** | 1.71E-60 | 0.247546143 | 0.999 | 1 | 4.12E-56 |
| **KAZN** | 1E-20 | 0.247467472 | 0.689 | 0.544 | 2.43E-16 |
| **AL022345.4** | 6.75E-53 | 0.247379346 | 0.353 | 0.144 | 1.63E-48 |
| **ARL6IP4** | 2.54E-26 | 0.2470513 | 0.951 | 0.879 | 6.12E-22 |
| **NAE1** | 1.58E-24 | 0.246996409 | 0.698 | 0.511 | 3.83E-20 |
| **GMPPB** | 7.12E-24 | 0.246934939 | 0.682 | 0.515 | 1.72E-19 |
| **WDR19** | 8.79E-26 | 0.246862891 | 0.78 | 0.617 | 2.12E-21 |
| **CD99** | 2.21E-19 | 0.246699928 | 0.71 | 0.554 | 5.34E-15 |
| **BLOC1S1** | 6.11E-32 | 0.24647158 | 0.98 | 0.931 | 1.48E-27 |

| | | | | | |
|---|---|---|---|---|---|
| **TMEM121** | 9.76E-47 | 0.24643337 | 0.336 | 0.142 | 2.36E-42 |
| **PNMA1** | 4.65E-21 | 0.246365485 | 0.696 | 0.557 | 1.12E-16 |
| **SNX3** | 6.25E-27 | 0.245778047 | 0.962 | 0.908 | 1.51E-22 |
| **SPATA24** | 5.05E-32 | 0.245756414 | 0.489 | 0.295 | 1.22E-27 |
| **CCL28** | 1.76E-14 | 0.245446201 | 0.431 | 0.318 | 4.25E-10 |
| **AP4M1** | 5.79E-24 | 0.24537581 | 0.609 | 0.443 | 1.4E-19 |
| **HAX1** | 2.14E-25 | 0.245286919 | 0.786 | 0.61 | 5.18E-21 |
| **IGFBP2** | 6.33E-39 | 0.245138052 | 1 | 1 | 1.53E-34 |
| **GNA14** | 1.62E-29 | 0.244878235 | 0.368 | 0.204 | 3.91E-25 |
| **DNAH9** | 1.11E-26 | 0.244858658 | 0.976 | 0.92 | 2.69E-22 |
| **TCEA2** | 1.12E-26 | 0.244458054 | 0.59 | 0.412 | 2.7E-22 |
| **C7orf50** | 4.63E-27 | 0.244429964 | 0.586 | 0.402 | 1.12E-22 |
| **SLC47A2** | 1.36E-39 | 0.244330826 | 0.207 | 0.073 | 3.29E-35 |
| **BBOX1** | 5.71E-35 | 0.244318184 | 0.336 | 0.165 | 1.38E-30 |
| **LRP2BP** | 1.01E-36 | 0.244307305 | 0.449 | 0.247 | 2.44E-32 |
| **EPN2** | 9.89E-27 | 0.24428833 | 0.704 | 0.529 | 2.39E-22 |
| **SEC62** | 3.2E-30 | 0.243891717 | 0.968 | 0.903 | 7.73E-26 |
| **FYB2** | 1.26E-27 | 0.243888427 | 0.311 | 0.164 | 3.05E-23 |
| **CDH26** | 6.46E-16 | 0.243661129 | 0.564 | 0.441 | 1.56E-11 |
| **CLINT1** | 2.33E-21 | 0.243574721 | 0.923 | 0.863 | 5.62E-17 |
| **NUB1** | 2.86E-25 | 0.242699702 | 0.691 | 0.515 | 6.91E-21 |
| **GFM2** | 2.16E-25 | 0.242637762 | 0.852 | 0.699 | 5.21E-21 |
| **LRRC4** | 2.37E-16 | 0.242391422 | 0.717 | 0.591 | 5.71E-12 |
| **C5orf15** | 1.59E-24 | 0.24231733 | 0.862 | 0.715 | 3.84E-20 |
| **GK** | 5.06E-21 | 0.242180759 | 0.637 | 0.489 | 1.22E-16 |
| **FNDC11** | 1.11E-42 | 0.241526703 | 0.344 | 0.156 | 2.69E-38 |
| **SAMHD1** | 7.24E-25 | 0.24145303 | 0.994 | 0.971 | 1.75E-20 |
| **STRBP** | 7.35E-27 | 0.241261575 | 0.95 | 0.87 | 1.78E-22 |
| **PIH1D1** | 2.43E-29 | 0.241172848 | 0.713 | 0.515 | 5.88E-25 |
| **MMAB** | 5.42E-26 | 0.240870663 | 0.696 | 0.513 | 1.31E-21 |
| **SSRP1** | 5.58E-22 | 0.240857615 | 0.754 | 0.592 | 1.35E-17 |
| **C12orf10** | 7.59E-24 | 0.240745246 | 0.712 | 0.543 | 1.83E-19 |
| **PROM1** | 6.18E-17 | 0.240708144 | 0.973 | 0.926 | 1.49E-12 |
| **RBM24** | 7.48E-23 | 0.240701806 | 0.541 | 0.377 | 1.81E-18 |
| **AC004130.2** | 4.57E-24 | 0.240663945 | 0.502 | 0.339 | 1.1E-19 |
| **MGST3** | 1.72E-29 | 0.240621585 | 0.986 | 0.962 | 4.15E-25 |
| **GADD45GIP1** | 3.23E-23 | 0.240551329 | 0.956 | 0.89 | 7.8E-19 |
| **EFCAB12** | 9.49E-29 | 0.240470499 | 0.734 | 0.539 | 2.29E-24 |

| | | | | | |
|---|---|---|---|---|---|
| **NDUFB3** | 4.26E-35 | 0.240367447 | 0.984 | 0.941 | 1.03E-30 |
| **ATG9B** | 3.89E-29 | 0.240228711 | 0.47 | 0.285 | 9.39E-25 |
| **AL162253.2** | 9.08E-39 | 0.23996094 | 0.326 | 0.15 | 2.19E-34 |
| **DTX3** | 5.74E-35 | 0.239928484 | 0.513 | 0.303 | 1.39E-30 |
| **CCDC122** | 3.38E-32 | 0.239840044 | 0.43 | 0.245 | 8.17E-28 |
| **SAT1** | 1.24E-12 | 0.239679717 | 1 | 1 | 3E-08 |
| **EFCAB7** | 2.4E-30 | 0.23965368 | 0.578 | 0.382 | 5.79E-26 |
| **CYTH2** | 4.7E-28 | 0.239277224 | 0.833 | 0.648 | 1.14E-23 |
| **C8orf48** | 7.75E-86 | 0.238824267 | 0.214 | 0.041 | 1.87E-81 |
| **SELENOH** | 3.93E-31 | 0.238445622 | 0.994 | 0.98 | 9.5E-27 |
| **ACOT13** | 1.33E-24 | 0.238297383 | 0.678 | 0.497 | 3.21E-20 |
| **SGSM3** | 2.23E-21 | 0.23824421 | 0.602 | 0.44 | 5.38E-17 |
| **TUSC3** | 1.81E-26 | 0.238072942 | 0.951 | 0.88 | 4.37E-22 |
| **CHMP2A** | 1.33E-35 | 0.237998799 | 0.99 | 0.948 | 3.21E-31 |
| **NUDT4** | 4.1E-10 | 0.23793917 | 0.793 | 0.693 | 9.91E-06 |
| **ZNF599** | 2.06E-26 | 0.237796363 | 0.491 | 0.32 | 4.96E-22 |
| **AL121956.6** | 1.51E-21 | 0.237755273 | 0.464 | 0.308 | 3.65E-17 |
| **CBX5** | 2.66E-23 | 0.237653644 | 0.847 | 0.712 | 6.42E-19 |
| **IQCE** | 2.03E-23 | 0.237602906 | 0.844 | 0.695 | 4.9E-19 |
| **GDA** | 9.45E-23 | 0.237556044 | 0.314 | 0.175 | 2.28E-18 |
| **SHOC2** | 1.49E-29 | 0.237507764 | 0.747 | 0.548 | 3.61E-25 |
| **TP53BP1** | 5.43E-24 | 0.236920342 | 0.914 | 0.804 | 1.31E-19 |
| **TMPRSS2** | 2.51E-20 | 0.236846308 | 0.616 | 0.471 | 6.07E-16 |
| **COTL1** | 1.66E-40 | 0.236798882 | 0.229 | 0.085 | 4.01E-36 |
| **MTLN** | 1.19E-21 | 0.236732678 | 0.607 | 0.442 | 2.86E-17 |
| **SLFN13** | 2.25E-22 | 0.236516762 | 0.87 | 0.734 | 5.43E-18 |
| **TTC21B** | 3.72E-18 | 0.236504101 | 0.483 | 0.339 | 8.98E-14 |
| **DLGAP1-AS1** | 3.4E-25 | 0.236434193 | 0.549 | 0.374 | 8.21E-21 |
| **CDC37L1-DT** | 1.92E-55 | 0.236264297 | 0.308 | 0.111 | 4.65E-51 |
| **USP18** | 1.72E-33 | 0.236264121 | 0.373 | 0.196 | 4.14E-29 |
| **PNLDC1** | 5.43E-64 | 0.236225777 | 0.239 | 0.066 | 1.31E-59 |
| **PKN1** | 7.48E-22 | 0.236086807 | 0.664 | 0.513 | 1.81E-17 |
| **UCN3** | 3.41E-14 | 0.235933926 | 0.152 | 0.077 | 8.23E-10 |
| **COMMD3** | 1.85E-23 | 0.235819326 | 0.786 | 0.634 | 4.46E-19 |
| **DYNC1LI1** | 5.87E-23 | 0.235774561 | 0.604 | 0.439 | 1.42E-18 |
| **ING2** | 1.04E-23 | 0.235754691 | 0.568 | 0.4 | 2.52E-19 |
| **SNRPA1** | 1.19E-22 | 0.235734322 | 0.677 | 0.516 | 2.88E-18 |
| **PMAIP1** | 2.76E-05 | 0.235732155 | 0.49 | 0.428 | 0.666169481 |

| | | | | | |
|---|---|---|---|---|---|
| **XBP1** | 5.78E-06 | 0.235712464 | 0.677 | 0.645 | 0.139529368 |
| **DHX30** | 2.39E-25 | 0.235478888 | 0.798 | 0.631 | 5.78E-21 |
| **ZCWPW1** | 2.11E-21 | 0.235441549 | 0.543 | 0.39 | 5.09E-17 |
| **TAGLN3** | 9.12E-33 | 0.235340381 | 0.256 | 0.113 | 2.2E-28 |
| **ADA2** | 7.92E-46 | 0.235315458 | 0.24 | 0.084 | 1.91E-41 |
| **CYP27A1** | 5.8E-25 | 0.235177829 | 0.468 | 0.299 | 1.4E-20 |
| **WDR93** | 3.78E-29 | 0.235140997 | 0.728 | 0.532 | 9.13E-25 |
| **KNDC1** | 2.15E-27 | 0.235088273 | 0.66 | 0.47 | 5.2E-23 |
| **AC008915.2** | 2.6E-23 | 0.234354135 | 0.571 | 0.406 | 6.28E-19 |
| **PRDX3** | 1.58E-26 | 0.234229954 | 0.874 | 0.75 | 3.82E-22 |
| **CEP126** | 1.82E-22 | 0.233826063 | 0.987 | 0.935 | 4.41E-18 |
| **SURF1** | 6.58E-23 | 0.233767345 | 0.774 | 0.619 | 1.59E-18 |
| **AC073370.1** | 1.82E-36 | 0.233446939 | 0.2 | 0.072 | 4.4E-32 |
| **FBXL13** | 3.14E-31 | 0.23344251 | 0.539 | 0.335 | 7.58E-27 |
| **ZNHIT1** | 4.49E-27 | 0.233441648 | 0.964 | 0.846 | 1.08E-22 |
| **AP3M2** | 1.16E-24 | 0.233346543 | 0.57 | 0.399 | 2.79E-20 |
| **ANKRD65** | 1.56E-24 | 0.233290459 | 0.776 | 0.602 | 3.78E-20 |
| **SYNGAP1** | 2.37E-24 | 0.233017056 | 0.586 | 0.414 | 5.71E-20 |
| **DYRK3** | 5.96E-22 | 0.232804158 | 0.576 | 0.419 | 1.44E-17 |
| **MSH6** | 1.2E-22 | 0.232753111 | 0.53 | 0.372 | 2.91E-18 |
| **NFU1** | 2.49E-28 | 0.232603478 | 0.632 | 0.438 | 6E-24 |
| **METTL5** | 2.44E-22 | 0.232535397 | 0.819 | 0.664 | 5.9E-18 |
| **LINC02363** | 8.55E-32 | 0.232518082 | 0.46 | 0.268 | 2.06E-27 |
| **WDR31** | 3.81E-27 | 0.232502585 | 0.632 | 0.434 | 9.2E-23 |
| **TCP1** | 5.24E-28 | 0.232100411 | 0.928 | 0.805 | 1.27E-23 |
| **KRT18** | 1.58E-06 | 0.231916288 | 0.896 | 0.918 | 0.038137316 |
| **SIAH3** | 2.29E-65 | 0.231532624 | 0.241 | 0.066 | 5.54E-61 |
| **URB1-AS1** | 2.97E-22 | 0.231486026 | 0.451 | 0.299 | 7.16E-18 |
| **CHCHD1** | 2.08E-28 | 0.231166622 | 0.934 | 0.797 | 5.02E-24 |
| **AL035587.2** | 3.2E-40 | 0.230991239 | 0.34 | 0.156 | 7.74E-36 |
| **POMT2** | 8.62E-25 | 0.230928122 | 0.327 | 0.183 | 2.08E-20 |
| **RND3** | 0.000948425 | 0.230686085 | 0.599 | 0.582 | 1 |
| **AP000692.2** | 3.05E-30 | 0.230455635 | 0.192 | 0.075 | 7.37E-26 |
| **COMMD8** | 9.73E-26 | 0.230439922 | 0.769 | 0.595 | 2.35E-21 |
| **NDUFA12** | 2.67E-22 | 0.230378821 | 0.773 | 0.607 | 6.44E-18 |
| **HSP90AA1** | 6.59E-14 | 0.230283085 | 1 | 1 | 1.59E-09 |
| **LYRM2** | 7.95E-26 | 0.230275343 | 0.937 | 0.799 | 1.92E-21 |
| **CEP41** | 3.21E-23 | 0.230235621 | 0.719 | 0.563 | 7.75E-19 |

| | | | | | |
|---|---|---|---|---|---|
| **MT-ATP6** | 0.384727914 | 0.230185754 | 0.998 | 1 | 1 |
| **TSPAN19** | 8.42E-17 | 0.23015508 | 0.954 | 0.927 | 2.03E-12 |
| **OAS2** | 4.77E-19 | 0.230000157 | 0.429 | 0.287 | 1.15E-14 |
| **HILPDA** | 1.49E-14 | 0.22983366 | 0.597 | 0.483 | 3.61E-10 |
| **APPL2** | 5.63E-21 | 0.22979976 | 0.672 | 0.521 | 1.36E-16 |
| **TRIM3** | 1.54E-20 | 0.229740517 | 0.291 | 0.167 | 3.72E-16 |
| **PYURF** | 5.91E-24 | 0.229728907 | 0.654 | 0.481 | 1.43E-19 |
| **TMEM173** | 5.71E-23 | 0.229306516 | 0.834 | 0.701 | 1.38E-18 |
| **TSPAN2** | 2.03E-25 | 0.229148228 | 0.237 | 0.114 | 4.9E-21 |
| **DHFR** | 9.77E-21 | 0.229120057 | 0.337 | 0.206 | 2.36E-16 |
| **FSIP1** | 9.3E-35 | 0.229051487 | 0.432 | 0.236 | 2.25E-30 |
| **GEMIN6** | 2.56E-23 | 0.228750529 | 0.582 | 0.423 | 6.17E-19 |
| **HYDIN** | 8.49E-22 | 0.228609951 | 0.994 | 0.975 | 2.05E-17 |
| **POLR2L** | 1.87E-32 | 0.228508581 | 0.984 | 0.966 | 4.51E-28 |
| **TNFRSF19** | 5.84E-15 | 0.228317677 | 0.744 | 0.642 | 1.41E-10 |
| **TTC16** | 7.75E-34 | 0.228300031 | 0.494 | 0.287 | 1.87E-29 |
| **GGCT** | 3.44E-23 | 0.22825127 | 0.627 | 0.45 | 8.3E-19 |
| **DNAJC7** | 8.02E-20 | 0.227915102 | 0.852 | 0.737 | 1.94E-15 |
| **PLAAT2** | 2.67E-16 | 0.227882918 | 0.552 | 0.402 | 6.44E-12 |
| **CTNNAL1** | 2.28E-17 | 0.22773906 | 0.633 | 0.499 | 5.51E-13 |
| **FAM177A1** | 1.92E-19 | 0.227736047 | 0.839 | 0.705 | 4.64E-15 |
| **SAP18** | 1.16E-33 | 0.227657839 | 0.992 | 0.979 | 2.8E-29 |
| **LIMS1** | 2.59E-14 | 0.227566696 | 0.933 | 0.897 | 6.26E-10 |
| **SRPK2** | 7.5E-27 | 0.227531003 | 0.822 | 0.629 | 1.81E-22 |
| **FGF14-AS2** | 8.63E-40 | 0.227397956 | 0.333 | 0.152 | 2.08E-35 |
| **ESD** | 1.53E-20 | 0.227236427 | 0.66 | 0.501 | 3.69E-16 |
| **WDR45B** | 9.29E-21 | 0.227184174 | 0.813 | 0.686 | 2.24E-16 |
| **FSD1L** | 5.01E-24 | 0.226863188 | 0.742 | 0.563 | 1.21E-19 |
| **PSMD9** | 4.32E-17 | 0.226749709 | 0.799 | 0.675 | 1.04E-12 |
| **INPP4B** | 4.09E-05 | 0.226604165 | 0.509 | 0.464 | 0.988401757 |
| **H1FX** | 1.86E-14 | 0.226015902 | 0.664 | 0.54 | 4.49E-10 |
| **ALDH3B1** | 1.74E-18 | 0.22588597 | 0.953 | 0.932 | 4.2E-14 |
| **ST13** | 1.09E-25 | 0.225651493 | 0.982 | 0.947 | 2.64E-21 |
| **NT5C3A** | 1.38E-21 | 0.225470233 | 0.686 | 0.529 | 3.34E-17 |
| **AGBL4** | 2.04E-41 | 0.225469761 | 0.339 | 0.153 | 4.92E-37 |
| **STIP1** | 2.4E-24 | 0.225457625 | 0.716 | 0.529 | 5.8E-20 |
| **MRPL58** | 2.34E-26 | 0.225342178 | 0.597 | 0.409 | 5.65E-22 |
| **YWHAE** | 2.57E-35 | 0.225093302 | 0.998 | 0.994 | 6.22E-31 |

| | | | | | |
|---|---|---|---|---|---|
| **SLC2A12** | 5.76E-19 | 0.225091124 | 0.417 | 0.276 | 1.39E-14 |
| **C18orf32** | 4.86E-32 | 0.225020072 | 0.982 | 0.922 | 1.17E-27 |
| **LY6G5C** | 2.82E-27 | 0.224975464 | 0.441 | 0.267 | 6.8E-23 |
| **WDR90** | 1.76E-17 | 0.224821013 | 0.778 | 0.624 | 4.25E-13 |
| **AL121820.2** | 3.11E-50 | 0.224608261 | 0.237 | 0.078 | 7.51E-46 |
| **C6orf226** | 9.25E-23 | 0.224400592 | 0.609 | 0.445 | 2.23E-18 |
| **CASTOR3** | 4.22E-18 | 0.224223253 | 0.498 | 0.359 | 1.02E-13 |
| **CNIH3** | 2.02E-20 | 0.224182286 | 0.394 | 0.251 | 4.88E-16 |
| **DDAH2** | 7.79E-15 | 0.22415878 | 0.583 | 0.461 | 1.88E-10 |
| **RNF8** | 1.01E-20 | 0.224110255 | 0.496 | 0.344 | 2.43E-16 |
| **MISP3** | 2.57E-32 | 0.224080287 | 0.466 | 0.269 | 6.21E-28 |
| **ERLEC1** | 1.15E-24 | 0.224056391 | 0.784 | 0.612 | 2.78E-20 |
| **HMGN3** | 2.34E-32 | 0.223930557 | 0.997 | 0.999 | 5.65E-28 |
| **CCT6B** | 1.38E-30 | 0.223691863 | 0.383 | 0.21 | 3.34E-26 |
| **SINHCAF** | 5.17E-20 | 0.223681984 | 0.799 | 0.655 | 1.25E-15 |
| **SDHAF4** | 1.17E-28 | 0.223514197 | 0.464 | 0.282 | 2.84E-24 |
| **IQCA1** | 1.72E-15 | 0.223433562 | 0.71 | 0.606 | 4.14E-11 |
| **CACNG4** | 6.02E-39 | 0.223370889 | 0.307 | 0.135 | 1.45E-34 |
| **ANG** | 2.43E-29 | 0.223368879 | 0.392 | 0.22 | 5.88E-25 |
| **SF3B2** | 1.34E-28 | 0.223105087 | 0.979 | 0.923 | 3.24E-24 |
| **PARD6A** | 7.01E-34 | 0.223032543 | 0.313 | 0.151 | 1.69E-29 |
| **GOSR1** | 5.52E-25 | 0.222958995 | 0.829 | 0.655 | 1.33E-20 |
| **ABHD14A** | 2.58E-21 | 0.222785757 | 0.45 | 0.302 | 6.22E-17 |
| **POLE3** | 2.97E-24 | 0.222688372 | 0.478 | 0.315 | 7.17E-20 |
| **LRGUK** | 1.57E-28 | 0.22267405 | 0.46 | 0.277 | 3.8E-24 |
| **CENPF** | 3.31E-23 | 0.222576242 | 0.417 | 0.259 | 7.98E-19 |
| **CLU** | 0.000117136 | 0.222252614 | 0.886 | 0.95 | 1 |
| **ISCA1** | 9.3E-20 | 0.222124176 | 0.692 | 0.562 | 2.25E-15 |
| **CD38** | 1.52E-16 | 0.221876462 | 0.65 | 0.519 | 3.66E-12 |
| **RABGAP1L** | 6.63E-21 | 0.221865293 | 0.901 | 0.776 | 1.6E-16 |
| **ABHD6** | 1E-18 | 0.221800591 | 0.352 | 0.227 | 2.42E-14 |
| **IFT140** | 2.27E-22 | 0.221717506 | 0.684 | 0.514 | 5.47E-18 |
| **PPM1G** | 2.02E-21 | 0.22169376 | 0.926 | 0.831 | 4.87E-17 |
| **SNAPC4** | 3.1E-25 | 0.221448506 | 0.478 | 0.307 | 7.48E-21 |
| **TMC4** | 6.51E-25 | 0.221442545 | 0.962 | 0.843 | 1.57E-20 |
| **TMC5** | 1.4E-16 | 0.221371943 | 0.999 | 0.995 | 3.38E-12 |
| **SPAG17** | 6.86E-18 | 0.221349152 | 0.983 | 0.953 | 1.66E-13 |
| **DAPP1** | 5.16E-18 | 0.220849675 | 0.852 | 0.736 | 1.25E-13 |

| | | | | | |
|---|---|---|---|---|---|
| **DUSP19** | 5.86E-24 | 0.220601119 | 0.504 | 0.337 | 1.41E-19 |
| **ABITRAM** | 8.88E-21 | 0.22036444 | 0.647 | 0.502 | 2.14E-16 |
| **AK8** | 1.46E-28 | 0.220239406 | 0.534 | 0.339 | 3.54E-24 |
| **TTF1** | 2.68E-21 | 0.220203049 | 0.622 | 0.465 | 6.47E-17 |
| **IL13RA1** | 3.82E-21 | 0.220141037 | 0.773 | 0.595 | 9.22E-17 |
| **AC010624.1** | 2.35E-34 | 0.219991834 | 0.364 | 0.185 | 5.67E-30 |
| **RABL2B** | 2.08E-18 | 0.219482109 | 0.964 | 0.902 | 5.02E-14 |
| **CCT7** | 2.24E-24 | 0.219423339 | 0.879 | 0.754 | 5.42E-20 |
| **SF3B4** | 3.31E-24 | 0.219030194 | 0.613 | 0.437 | 8E-20 |
| **TDRP** | 6.39E-26 | 0.218982659 | 0.374 | 0.218 | 1.54E-21 |
| **AC097534.2** | 1E-34 | 0.218852613 | 0.341 | 0.167 | 2.43E-30 |
| **MROH9** | 5.15E-17 | 0.218639702 | 0.391 | 0.262 | 1.24E-12 |
| **AC027117.2** | 4.59E-37 | 0.218304029 | 0.274 | 0.117 | 1.11E-32 |
| **MRPL20** | 5.71E-27 | 0.218272993 | 0.971 | 0.905 | 1.38E-22 |
| **CCNDBP1** | 1.61E-22 | 0.218234821 | 0.847 | 0.696 | 3.89E-18 |
| **UPF3A** | 3.05E-23 | 0.21773773 | 0.838 | 0.672 | 7.36E-19 |
| **TULP3** | 1.05E-19 | 0.217365007 | 0.706 | 0.56 | 2.53E-15 |
| **TP73** | 2.86E-22 | 0.217339067 | 0.711 | 0.541 | 6.9E-18 |
| **RPAP3** | 1.81E-22 | 0.217265541 | 0.683 | 0.515 | 4.37E-18 |
| **RNF6** | 7.23E-21 | 0.217051403 | 0.837 | 0.708 | 1.75E-16 |
| **LINC01014** | 8.61E-42 | 0.217047581 | 0.242 | 0.089 | 2.08E-37 |
| **DDX60** | 9.57E-10 | 0.217040917 | 0.356 | 0.264 | 2.31E-05 |
| **UQCRB** | 9.84E-33 | 0.216964148 | 0.994 | 0.987 | 2.38E-28 |
| **CASC2** | 3.47E-23 | 0.216951818 | 0.789 | 0.616 | 8.37E-19 |
| **MDK** | 6.64E-14 | 0.216796398 | 0.37 | 0.539 | 1.6E-09 |
| **PTRHD1** | 1.32E-17 | 0.216759498 | 0.813 | 0.678 | 3.2E-13 |
| **DNAJC15** | 1.02E-21 | 0.216663781 | 0.703 | 0.524 | 2.47E-17 |
| **IQCH** | 4.56E-26 | 0.216641098 | 0.629 | 0.438 | 1.1E-21 |
| **PFN2** | 2.08E-18 | 0.216632699 | 0.943 | 0.845 | 5.03E-14 |
| **MGAT5** | 7.61E-14 | 0.216418081 | 0.463 | 0.348 | 1.84E-09 |
| **ANK3-DT** | 5.88E-41 | 0.215399768 | 0.291 | 0.122 | 1.42E-36 |
| **ECI2** | 9.27E-18 | 0.215356934 | 0.339 | 0.216 | 2.24E-13 |
| **MSL3** | 3.85E-20 | 0.215162388 | 0.683 | 0.537 | 9.3E-16 |
| **FAM227B** | 3.5E-28 | 0.21503652 | 0.41 | 0.239 | 8.45E-24 |
| **TMEM125** | 3.48E-21 | 0.214990187 | 0.749 | 0.586 | 8.4E-17 |
| **JPT2** | 1.16E-15 | 0.214777759 | 0.911 | 0.821 | 2.8E-11 |
| **NDUFB6** | 1.33E-18 | 0.214525326 | 0.777 | 0.619 | 3.2E-14 |
| **CCDC25** | 6.79E-23 | 0.214332823 | 0.754 | 0.574 | 1.64E-18 |

| | | | | | |
|---|---|---|---|---|---|
| **STK40** | 4.38E-18 | 0.214308684 | 0.488 | 0.346 | 1.06E-13 |
| **AL133331.1** | 1.38E-32 | 0.214292655 | 0.186 | 0.069 | 3.33E-28 |
| **CCDC102A** | 1.35E-32 | 0.214019879 | 0.359 | 0.184 | 3.27E-28 |
| **UBE2S** | 8.42E-19 | 0.213862141 | 0.506 | 0.359 | 2.03E-14 |
| **YTHDF3-AS1** | 2.65E-30 | 0.213799157 | 0.398 | 0.22 | 6.41E-26 |
| **AL031666.1** | 3.74E-42 | 0.21370622 | 0.242 | 0.09 | 9.04E-38 |
| **RRAGD** | 2.22E-17 | 0.213520618 | 0.618 | 0.477 | 5.36E-13 |
| **GALK2** | 1.35E-19 | 0.213468822 | 0.574 | 0.43 | 3.25E-15 |
| **SELENBP1** | 6.45E-15 | 0.213413096 | 0.804 | 0.705 | 1.56E-10 |
| **TOMM34** | 2.67E-18 | 0.213262169 | 0.588 | 0.442 | 6.44E-14 |
| **GFER** | 4.63E-24 | 0.213245442 | 0.671 | 0.482 | 1.12E-19 |
| **HNRNPF** | 3.99E-29 | 0.212946283 | 0.993 | 0.976 | 9.64E-25 |
| **CACYBP** | 5.67E-21 | 0.212181681 | 0.71 | 0.549 | 1.37E-16 |
| **TAGLN2** | 2.72E-17 | 0.212021027 | 0.997 | 0.975 | 6.58E-13 |
| **EGLN3** | 1.24E-20 | 0.211814297 | 0.498 | 0.34 | 2.99E-16 |
| **AC127070.2** | 1.37E-29 | 0.211785864 | 0.369 | 0.2 | 3.32E-25 |
| **CITED4** | 5.45E-13 | 0.211727942 | 0.433 | 0.317 | 1.32E-08 |
| **CEP97** | 3.22E-19 | 0.211549423 | 0.8 | 0.676 | 7.77E-15 |
| **ANKRD18A** | 6.39E-15 | 0.211315769 | 0.393 | 0.267 | 1.54E-10 |
| **AL022328.4** | 1.01E-51 | 0.211176683 | 0.226 | 0.07 | 2.44E-47 |
| **ZNF710-AS1** | 8.29E-33 | 0.211154553 | 0.342 | 0.172 | 2E-28 |
| **NASP** | 8.88E-21 | 0.211135639 | 0.838 | 0.679 | 2.15E-16 |
| **CDC5L** | 5.08E-23 | 0.210993833 | 0.801 | 0.631 | 1.23E-18 |
| **FIBP** | 1.73E-18 | 0.210861448 | 0.703 | 0.551 | 4.19E-14 |
| **SLC23A1** | 2.34E-13 | 0.210674429 | 0.343 | 0.237 | 5.65E-09 |
| **COA6** | 5.19E-23 | 0.210382361 | 0.736 | 0.556 | 1.25E-18 |
| **IFT80** | 1.73E-17 | 0.210177997 | 0.889 | 0.801 | 4.18E-13 |
| **ELN-AS1** | 1.37E-45 | 0.210026592 | 0.27 | 0.101 | 3.32E-41 |
| **DOC2A** | 1.45E-26 | 0.209922534 | 0.31 | 0.162 | 3.5E-22 |
| **CRYM** | 8.76E-28 | 0.209700319 | 0.181 | 0.072 | 2.12E-23 |
| **SMIM8** | 6.16E-14 | 0.209362515 | 0.49 | 0.369 | 1.49E-09 |
| **RHPN1** | 3.18E-22 | 0.209345612 | 0.651 | 0.483 | 7.68E-18 |
| **DEK** | 1.57E-15 | 0.209278443 | 0.847 | 0.755 | 3.8E-11 |
| **SLPI** | 0.453409834 | 0.209172254 | 0.997 | 0.999 | 1 |
| **CD200R1L-AS1** | 2.62E-34 | 0.208919729 | 0.321 | 0.152 | 6.33E-30 |
| **IFITM3** | 1.84E-09 | 0.208885044 | 0.469 | 0.601 | 4.44E-05 |
| **CLBA1** | 9.49E-26 | 0.208862009 | 0.502 | 0.32 | 2.29E-21 |
| **TMCO1** | 2.56E-19 | 0.208774353 | 0.882 | 0.753 | 6.18E-15 |

| | | | | | |
|---|---|---|---|---|---|
| **ATL1** | 5.76E-28 | 0.208614044 | 0.394 | 0.224 | 1.39E-23 |
| **ZC2HC1A** | 1.41E-19 | 0.208574018 | 0.884 | 0.768 | 3.41E-15 |
| **HIKESHI** | 2.12E-19 | 0.208363136 | 0.559 | 0.401 | 5.13E-15 |
| **BBS4** | 2.39E-19 | 0.20823567 | 0.637 | 0.493 | 5.78E-15 |
| **LRRC56** | 1.03E-22 | 0.208196824 | 0.569 | 0.394 | 2.49E-18 |
| **AL133320.2** | 3.82E-22 | 0.207993325 | 0.493 | 0.328 | 9.23E-18 |
| **COA5** | 2.98E-19 | 0.2078609 | 0.693 | 0.536 | 7.19E-15 |
| **TTC1** | 3.67E-23 | 0.207697077 | 0.714 | 0.528 | 8.85E-19 |
| **CDKL1** | 8.33E-18 | 0.207683825 | 0.748 | 0.591 | 2.01E-13 |
| **KIAA0556** | 3.28E-20 | 0.207664418 | 0.704 | 0.553 | 7.93E-16 |
| **C9orf72** | 4.37E-18 | 0.207162496 | 0.82 | 0.695 | 1.06E-13 |
| **CCT5** | 3.91E-21 | 0.207008946 | 0.929 | 0.813 | 9.44E-17 |
| **LARP6** | 8.5E-12 | 0.206940997 | 0.447 | 0.351 | 2.05E-07 |
| **DNAJC1** | 4.83E-20 | 0.206889065 | 0.558 | 0.4 | 1.17E-15 |
| **ISCU** | 8.65E-19 | 0.206723909 | 0.889 | 0.764 | 2.09E-14 |
| **SLC25A14** | 5.76E-27 | 0.206723502 | 0.511 | 0.322 | 1.39E-22 |
| **CFAP97D2** | 3.08E-10 | 0.206393613 | 0.369 | 0.277 | 7.43E-06 |
| **MUCL1** | 1.12E-11 | 0.206251455 | 0.342 | 0.242 | 2.7E-07 |
| **CCDC187** | 4.39E-22 | 0.206179299 | 0.783 | 0.608 | 1.06E-17 |
| **STX18** | 5.94E-18 | 0.205986501 | 0.599 | 0.457 | 1.43E-13 |
| **CCDC40** | 6.46E-19 | 0.205891871 | 0.903 | 0.788 | 1.56E-14 |
| **SVBP** | 5.1E-21 | 0.205727669 | 0.618 | 0.444 | 1.23E-16 |
| **AC010642.2** | 2.44E-19 | 0.205727669 | 0.594 | 0.443 | 5.88E-15 |
| **MDP1** | 5.44E-20 | 0.20548669 | 0.496 | 0.347 | 1.31E-15 |
| **C3orf67** | 7.07E-26 | 0.205463861 | 0.492 | 0.309 | 1.71E-21 |
| **CERKL** | 1.57E-22 | 0.204915956 | 0.743 | 0.564 | 3.79E-18 |
| **C2orf50** | 2.65E-29 | 0.204538655 | 0.368 | 0.199 | 6.39E-25 |
| **PDZK1IP1** | 3.14E-08 | 0.204529341 | 0.141 | 0.086 | 0.00075859 |
| **GTF2A2** | 1.78E-21 | 0.204526816 | 0.924 | 0.808 | 4.29E-17 |
| **SIX1** | 2.87E-18 | 0.204493937 | 0.897 | 0.806 | 6.94E-14 |
| **PDE6B** | 1.65E-35 | 0.204426218 | 0.314 | 0.146 | 3.97E-31 |
| **MICOS13** | 6.86E-20 | 0.204403159 | 0.981 | 0.929 | 1.66E-15 |
| **BICC1** | 2.42E-16 | 0.204356911 | 0.281 | 0.173 | 5.86E-12 |
| **MID1IP1** | 9.85E-18 | 0.204265006 | 0.747 | 0.609 | 2.38E-13 |
| **RSPH10B** | 4E-29 | 0.204237805 | 0.463 | 0.272 | 9.67E-25 |
| **NDUFB10** | 7.45E-20 | 0.203972678 | 0.964 | 0.912 | 1.8E-15 |
| **MRPL51** | 6.17E-19 | 0.203749718 | 0.92 | 0.82 | 1.49E-14 |
| **ZNF232** | 3.29E-28 | 0.203722901 | 0.342 | 0.184 | 7.94E-24 |

| | | | | | |
|---|---|---|---|---|---|
| **EMC6** | 7.78E-18 | 0.203494325 | 0.861 | 0.737 | 1.88E-13 |
| **AC027644.3** | 7.72E-20 | 0.203424742 | 0.606 | 0.448 | 1.87E-15 |
| **LRRC45** | 8.58E-19 | 0.203306843 | 0.679 | 0.531 | 2.07E-14 |
| **CYB5D2** | 8.08E-14 | 0.203256335 | 0.448 | 0.326 | 1.95E-09 |
| **CCDC91** | 4.76E-20 | 0.203192273 | 0.677 | 0.505 | 1.15E-15 |
| **AC005476.2** | 2.32E-35 | 0.20312679 | 0.314 | 0.146 | 5.6E-31 |
| **PPP6R1** | 5.19E-20 | 0.203076245 | 0.784 | 0.63 | 1.25E-15 |
| **FBXO16** | 1.1E-24 | 0.202719031 | 0.434 | 0.268 | 2.67E-20 |
| **YEATS4** | 2.85E-21 | 0.202548683 | 0.463 | 0.307 | 6.89E-17 |
| **CCL14** | 2.63E-37 | 0.202464851 | 0.264 | 0.11 | 6.35E-33 |
| **VPS13B-DT** | 3.03E-33 | 0.202365504 | 0.31 | 0.148 | 7.33E-29 |
| **SIGIRR** | 3.86E-17 | 0.202315472 | 0.778 | 0.652 | 9.33E-13 |
| **METAP2** | 5.39E-19 | 0.202310624 | 0.817 | 0.672 | 1.3E-14 |
| **PRPF6** | 4.78E-21 | 0.202061603 | 0.813 | 0.655 | 1.15E-16 |
| **COPS5** | 2.08E-20 | 0.202028244 | 0.654 | 0.492 | 5.03E-16 |
| **AC016876.1** | 1.77E-32 | 0.202009285 | 0.296 | 0.139 | 4.28E-28 |
| **IFNAR2** | 2.99E-20 | 0.201647674 | 0.619 | 0.46 | 7.23E-16 |
| **DNAJC19** | 3.98E-23 | 0.201516421 | 0.651 | 0.465 | 9.62E-19 |
| **TSPYL5** | 2.65E-17 | 0.201435349 | 0.403 | 0.269 | 6.4E-13 |
| **CYB561** | 1.69E-18 | 0.201348787 | 0.896 | 0.787 | 4.09E-14 |
| **CISD3** | 4.12E-17 | 0.201292331 | 0.548 | 0.407 | 9.96E-13 |
| **POLR1C** | 1.69E-20 | 0.201230045 | 0.521 | 0.363 | 4.07E-16 |
| **AP001453.1** | 1.92E-53 | 0.201152714 | 0.211 | 0.06 | 4.64E-49 |
| **ZNFX1** | 3.23E-15 | 0.201024056 | 0.648 | 0.516 | 7.8E-11 |
| **CD4** | 5.95E-33 | 0.201015089 | 0.261 | 0.114 | 1.44E-28 |
| **PSMG1** | 2.85E-21 | 0.201001781 | 0.359 | 0.219 | 6.88E-17 |
| **MRPS7** | 8.62E-20 | 0.200919212 | 0.687 | 0.528 | 2.08E-15 |
| **DANCR** | 1.01E-14 | 0.200866638 | 0.603 | 0.471 | 2.44E-10 |
| **RBM23** | 1.24E-19 | 0.200593693 | 0.777 | 0.61 | 3E-15 |
| **MXI1** | 2.84E-13 | 0.200583319 | 0.647 | 0.533 | 6.86E-09 |
| **NWD1** | 2.42E-14 | 0.200453803 | 0.966 | 0.938 | 5.83E-10 |
| **SPART** | 2.41E-18 | 0.200154469 | 0.843 | 0.709 | 5.81E-14 |
| **AKR1B10** | 1.04E-10 | -0.200041384 | 0.146 | 0.244 | 2.5E-06 |
| **TPD52L1** | 6.33E-11 | -0.200289106 | 0.201 | 0.298 | 1.53E-06 |
| **M6PR** | 1.84E-11 | -0.200535298 | 0.367 | 0.468 | 4.45E-07 |
| **MAP3K2** | 7.02E-10 | -0.201371761 | 0.621 | 0.679 | 1.7E-05 |
| **MYL6** | 1.47E-21 | -0.201819941 | 0.999 | 1 | 3.55E-17 |
| **DHRS7** | 2.77E-14 | -0.201895166 | 0.159 | 0.275 | 6.7E-10 |

| | | | | | |
|---|---|---|---|---|---|
| **KIAA1109** | 9.1E-09 | -0.2022275 | 0.419 | 0.496 | 0.000219711 |
| **LINC00472** | 1.66E-15 | -0.202389813 | 0.15 | 0.276 | 4.02E-11 |
| **ELL2** | 4.8E-12 | -0.202591414 | 0.244 | 0.352 | 1.16E-07 |
| **NBEAL1** | 2.26E-08 | -0.202620237 | 0.534 | 0.582 | 0.000545381 |
| **AHCYL1** | 2.43E-12 | -0.202636127 | 0.766 | 0.809 | 5.88E-08 |
| **STON2** | 8.93E-15 | -0.203613351 | 0.132 | 0.244 | 2.16E-10 |
| **KDM7A** | 1.27E-11 | -0.203975162 | 0.307 | 0.411 | 3.07E-07 |
| **NPM1** | 1.98E-15 | -0.204346815 | 0.953 | 0.968 | 4.78E-11 |
| **PRKX** | 6.53E-08 | -0.204415638 | 0.48 | 0.541 | 0.001578179 |
| **PATJ** | 1.91E-09 | -0.204730255 | 0.456 | 0.532 | 4.6E-05 |
| **USP34** | 2.22E-09 | -0.205050104 | 0.64 | 0.684 | 5.37E-05 |
| **CD151** | 5.1E-16 | -0.205306239 | 0.11 | 0.225 | 1.23E-11 |
| **RPS13** | 1.12E-25 | -0.205599724 | 0.989 | 1 | 2.7E-21 |
| **SSR2** | 5.3E-16 | -0.20641436 | 0.197 | 0.328 | 1.28E-11 |
| **BAZ2B** | 2.25E-09 | -0.206781636 | 0.602 | 0.671 | 5.43E-05 |
| **POGZ** | 4.9E-13 | -0.207011474 | 0.321 | 0.432 | 1.18E-08 |
| **KHDC4** | 3.53E-15 | -0.207043905 | 0.367 | 0.503 | 8.54E-11 |
| **KLF10** | 3.19E-10 | -0.207494717 | 0.324 | 0.418 | 7.7E-06 |
| **HNRNPR** | 2.33E-11 | -0.208033567 | 0.716 | 0.751 | 5.64E-07 |
| **YIPF4** | 1.68E-10 | -0.208052239 | 0.538 | 0.605 | 4.06E-06 |
| **SYNE2** | 5.07E-08 | -0.208075963 | 0.97 | 0.974 | 0.001224125 |
| **CLIP1** | 1.35E-13 | -0.208093765 | 0.787 | 0.867 | 3.25E-09 |
| **MED13L** | 5.73E-09 | -0.208132037 | 0.448 | 0.514 | 0.000138492 |
| **TMEM33** | 8.29E-12 | -0.208166935 | 0.414 | 0.509 | 2E-07 |
| **CAPZA1** | 2.39E-11 | -0.208186627 | 0.651 | 0.695 | 5.76E-07 |
| **FAM118A** | 1.3E-15 | -0.208339256 | 0.132 | 0.249 | 3.14E-11 |
| **SELENOP** | 2.35E-20 | -0.208491201 | 0.064 | 0.19 | 5.69E-16 |
| **GSPT1** | 1.11E-10 | -0.208982584 | 0.654 | 0.699 | 2.68E-06 |
| **SERPINB5** | 5E-16 | -0.209144663 | 0.264 | 0.402 | 1.21E-11 |
| **TTC14** | 3.21E-13 | -0.209259914 | 0.278 | 0.395 | 7.75E-09 |
| **KANSL1** | 4.09E-11 | -0.209597378 | 0.468 | 0.561 | 9.88E-07 |
| **HMGA1** | 7.84E-15 | -0.209785494 | 0.147 | 0.261 | 1.89E-10 |
| **TMEM161B-AS1** | 4.96E-10 | -0.209788427 | 0.572 | 0.634 | 1.2E-05 |
| **PFN1** | 8.26E-18 | -0.209896662 | 0.844 | 0.91 | 1.99E-13 |
| **RBM5** | 5.02E-11 | -0.210410092 | 0.491 | 0.564 | 1.21E-06 |
| **SCARB2** | 5.53E-12 | -0.210534943 | 0.423 | 0.532 | 1.34E-07 |
| **MLF 2.00** | 5.68E-12 | -0.210680588 | 0.479 | 0.564 | 1.37E-07 |
| **SF3B1** | 8.14E-12 | -0.210819646 | 0.868 | 0.873 | 1.97E-07 |

| | | | | | |
|---|---|---|---|---|---|
| **MT1E** | 1.99E-11 | -0.210920143 | 0.171 | 0.273 | 4.81E-07 |
| **SLC6A14** | 4.16E-20 | -0.211214172 | 0.097 | 0.233 | 1E-15 |
| **GPATCH2L** | 7.5E-15 | -0.211972414 | 0.263 | 0.392 | 1.81E-10 |
| **PLEC** | 5.89E-10 | -0.212375454 | 0.543 | 0.613 | 1.42E-05 |
| **MTUS1** | 1.82E-11 | -0.213221312 | 0.49 | 0.579 | 4.4E-07 |
| **ANKRD36C** | 1.19E-11 | -0.213300746 | 0.414 | 0.538 | 2.88E-07 |
| **ARHGEF12** | 1.99E-09 | -0.21382342 | 0.501 | 0.568 | 4.82E-05 |
| **ADGRG7** | 2.98E-24 | -0.214231621 | 0.009 | 0.121 | 7.19E-20 |
| **PRNP** | 5.48E-14 | -0.214776919 | 0.201 | 0.315 | 1.32E-09 |
| **PPM1L** | 3.22E-16 | -0.214957447 | 0.188 | 0.318 | 7.78E-12 |
| **ARHGAP5** | 2.09E-10 | -0.215122811 | 0.837 | 0.846 | 5.04E-06 |
| **GPC1** | 1.53E-14 | -0.215137954 | 0.186 | 0.306 | 3.69E-10 |
| **MYOF** | 2.12E-15 | -0.215213626 | 0.631 | 0.756 | 5.13E-11 |
| **RPS17** | 7.72E-20 | -0.215239573 | 0.987 | 0.994 | 1.86E-15 |
| **EMC10** | 2.37E-14 | -0.215444215 | 0.273 | 0.394 | 5.73E-10 |
| **ARRDC3** | 1.58E-19 | -0.215948033 | 0.076 | 0.2 | 3.81E-15 |
| **SYPL1** | 2.96E-11 | -0.216059087 | 0.588 | 0.654 | 7.16E-07 |
| **JUP** | 2.89E-14 | -0.216089791 | 0.337 | 0.459 | 6.97E-10 |
| **RPS27A** | 8.22E-30 | -0.216609808 | 0.996 | 1 | 1.99E-25 |
| **RBM4** | 6.61E-12 | -0.216906471 | 0.631 | 0.688 | 1.6E-07 |
| **MYH14** | 9.44E-12 | -0.21720271 | 0.806 | 0.818 | 2.28E-07 |
| **INTS6** | 9.87E-13 | -0.217863979 | 0.259 | 0.367 | 2.38E-08 |
| **EIF4A2** | 4.19E-14 | -0.217979573 | 0.853 | 0.932 | 1.01E-09 |
| **SMARCC1** | 1.69E-10 | -0.218074812 | 0.507 | 0.569 | 4.07E-06 |
| **KIAA1217** | 9.05E-13 | -0.218537017 | 0.606 | 0.692 | 2.19E-08 |
| **ITPRID2** | 1E-14 | -0.218885196 | 0.3 | 0.422 | 2.42E-10 |
| **PCMTD1** | 1.19E-11 | -0.219175383 | 0.433 | 0.526 | 2.88E-07 |
| **PABPC4** | 2.63E-14 | -0.219434971 | 0.288 | 0.412 | 6.34E-10 |
| **STEAP3** | 7.41E-10 | -0.219482751 | 0.664 | 0.719 | 1.79E-05 |
| **ARHGAP21** | 4.02E-13 | -0.219778194 | 0.447 | 0.557 | 9.7E-09 |
| **ABRACL** | 4.45E-13 | -0.219832173 | 0.422 | 0.537 | 1.07E-08 |
| **TMEM167A** | 6.58E-13 | -0.220442637 | 0.376 | 0.485 | 1.59E-08 |
| **CAPN2** | 6.28E-15 | -0.220540177 | 0.864 | 0.889 | 1.52E-10 |
| **RPL17** | 9.44E-29 | -0.22101227 | 0.996 | 0.999 | 2.28E-24 |
| **SLC38A1** | 3.94E-16 | -0.221119844 | 0.133 | 0.253 | 9.51E-12 |
| **ANK3** | 6.96E-08 | -0.221157154 | 0.887 | 0.868 | 0.001680237 |
| **SSR4** | 2.69E-16 | -0.221464514 | 0.944 | 0.955 | 6.49E-12 |
| **S100A16** | 2.5E-26 | -0.221692764 | 0.07 | 0.226 | 6.04E-22 |

| | | | | | |
|---|---|---|---|---|---|
| **MIB1** | 9.7E-12 | -0.221979314 | 0.41 | 0.502 | 2.34E-07 |
| **ZNF587** | 4.68E-19 | -0.22204282 | 0.179 | 0.322 | 1.13E-14 |
| **ERBIN** | 1.22E-13 | -0.222298159 | 0.631 | 0.704 | 2.95E-09 |
| **AGO3** | 1.68E-17 | -0.222588465 | 0.202 | 0.342 | 4.05E-13 |
| **EIF4G1** | 2.69E-12 | -0.222651459 | 0.691 | 0.737 | 6.49E-08 |
| **TES** | 8.62E-12 | -0.222830553 | 0.473 | 0.562 | 2.08E-07 |
| **MIF** | 2.28E-17 | -0.222846221 | 0.989 | 0.996 | 5.5E-13 |
| **KIDINS220** | 1.13E-12 | -0.222882906 | 0.347 | 0.452 | 2.72E-08 |
| **ARL6IP5** | 2.52E-14 | -0.222918667 | 0.716 | 0.771 | 6.09E-10 |
| **SEPTIN2** | 3.07E-14 | -0.223047721 | 0.79 | 0.825 | 7.41E-10 |
| **RPL7A** | 5.72E-35 | -0.224113055 | 0.993 | 1 | 1.38E-30 |
| **HNRNPU** | 6.64E-16 | -0.224146448 | 0.928 | 0.961 | 1.6E-11 |
| **PGAM1** | 5.66E-13 | -0.224291189 | 0.336 | 0.442 | 1.37E-08 |
| **EIF1AX** | 5.83E-12 | -0.224550393 | 0.479 | 0.568 | 1.41E-07 |
| **C3orf52** | 2.69E-19 | -0.224616955 | 0.147 | 0.286 | 6.49E-15 |
| **OGA** | 6.34E-11 | -0.224764748 | 0.447 | 0.524 | 1.53E-06 |
| **CAND1** | 2.22E-12 | -0.224821282 | 0.599 | 0.649 | 5.36E-08 |
| **MPZL2** | 7.6E-19 | -0.225093753 | 0.249 | 0.402 | 1.84E-14 |
| **RPL35** | 3.28E-16 | -0.225240581 | 0.957 | 0.986 | 7.91E-12 |
| **UGCG** | 1.98E-13 | -0.225267287 | 0.663 | 0.727 | 4.79E-09 |
| **OTUD1** | 8.7E-15 | -0.22579798 | 0.307 | 0.435 | 2.1E-10 |
| **ERGIC2** | 1.8E-13 | -0.226025291 | 0.451 | 0.547 | 4.34E-09 |
| **UQCRH** | 4.54E-17 | -0.226522709 | 0.89 | 0.925 | 1.1E-12 |
| **RPL24** | 2.2E-23 | -0.22688642 | 0.991 | 0.998 | 5.31E-19 |
| **CHD9** | 2.32E-13 | -0.228211235 | 0.57 | 0.67 | 5.61E-09 |
| **SSR1** | 1.43E-14 | -0.228310226 | 0.322 | 0.441 | 3.46E-10 |
| **IER3** | 7.68E-25 | -0.228581017 | 0.092 | 0.256 | 1.85E-20 |
| **AP2M1** | 3.35E-14 | -0.228851837 | 0.724 | 0.784 | 8.09E-10 |
| **HNRNPAB** | 2.24E-14 | -0.229235846 | 0.536 | 0.638 | 5.41E-10 |
| **JMJD1C** | 6.34E-11 | -0.229410141 | 0.572 | 0.641 | 1.53E-06 |
| **PTTG1IP** | 6.9E-15 | -0.229686353 | 0.629 | 0.74 | 1.67E-10 |
| **NPC2** | 1.45E-11 | -0.230192829 | 0.891 | 0.905 | 3.5E-07 |
| **CHMP1B** | 1.04E-15 | -0.230254391 | 0.306 | 0.434 | 2.5E-11 |
| **TUBA1C** | 1.84E-19 | -0.23044967 | 0.181 | 0.329 | 4.44E-15 |
| **RPS24** | 3.02E-32 | -0.230525216 | 0.996 | 1 | 7.28E-28 |
| **SH3PXD2A** | 1.34E-15 | -0.230755146 | 0.162 | 0.28 | 3.23E-11 |
| **MFSD14C** | 6.57E-14 | -0.231556775 | 0.506 | 0.594 | 1.59E-09 |
| **SPG7** | 4.81E-13 | -0.23247513 | 0.49 | 0.579 | 1.16E-08 |

| | | | | | |
|---|---|---|---|---|---|
| **CNN2** | 1.14E-18 | -0.23274166 | 0.154 | 0.291 | 2.76E-14 |
| **SLK** | 1.06E-13 | -0.232862614 | 0.701 | 0.767 | 2.56E-09 |
| **GOLGA4** | 6.79E-13 | -0.232876813 | 0.671 | 0.767 | 1.64E-08 |
| **MT-CO3** | 2.56E-28 | -0.233135922 | 0.999 | 1 | 6.18E-24 |
| **UHMK1** | 8.86E-13 | -0.233197163 | 0.61 | 0.672 | 2.14E-08 |
| **TNFRSF21** | 3.51E-12 | -0.234212836 | 0.444 | 0.539 | 8.47E-08 |
| **HSD17B13** | 1.7E-14 | -0.234229662 | 0.101 | 0.206 | 4.11E-10 |
| **XIAP** | 7.72E-15 | -0.234239497 | 0.484 | 0.597 | 1.86E-10 |
| **MMP10** | 9.13E-13 | -0.234329451 | 0.027 | 0.101 | 2.21E-08 |
| **GALNT4** | 9.53E-17 | -0.234516247 | 0.27 | 0.407 | 2.3E-12 |
| **EEF1B2** | 3.52E-20 | -0.234690025 | 0.901 | 0.944 | 8.5E-16 |
| **ATP6V0E1** | 1.34E-21 | -0.235064926 | 0.96 | 0.966 | 3.25E-17 |
| **CPLANE1** | 6E-11 | -0.235312901 | 0.757 | 0.781 | 1.45E-06 |
| **FABP5** | 5.03E-21 | -0.235325486 | 0.263 | 0.465 | 1.21E-16 |
| **CST3** | 7.23E-19 | -0.235491232 | 0.952 | 0.972 | 1.75E-14 |
| **PSD3** | 1.5E-10 | -0.23602767 | 0.611 | 0.653 | 3.63E-06 |
| **ZNF292** | 1.37E-13 | -0.236558312 | 0.493 | 0.587 | 3.32E-09 |
| **RPS2** | 3.19E-31 | -0.236646801 | 0.992 | 1 | 7.71E-27 |
| **MCL1** | 1.11E-16 | -0.236712497 | 0.468 | 0.609 | 2.67E-12 |
| **TENT5A** | 1.38E-16 | -0.237131626 | 0.167 | 0.293 | 3.32E-12 |
| **BACE2** | 2.11E-13 | -0.237797796 | 0.321 | 0.439 | 5.09E-09 |
| **MT2A** | 3.48E-17 | -0.23822627 | 0.433 | 0.586 | 8.4E-13 |
| **KMT2C** | 1.43E-11 | -0.238604089 | 0.528 | 0.596 | 3.44E-07 |
| **ZKSCAN1** | 2.67E-18 | -0.238622338 | 0.93 | 0.961 | 6.45E-14 |
| **PLEKHA1** | 4.66E-16 | -0.238627745 | 0.199 | 0.324 | 1.13E-11 |
| **PBX1** | 1.08E-11 | -0.239186411 | 0.501 | 0.583 | 2.6E-07 |
| **ANKRD11** | 2.13E-15 | -0.239706292 | 0.862 | 0.877 | 5.14E-11 |
| **HMGCS1** | 3.22E-12 | -0.241025692 | 0.224 | 0.33 | 7.78E-08 |
| **RPS23** | 8.74E-29 | -0.24128227 | 0.988 | 0.999 | 2.11E-24 |
| **NAMPT** | 1.5E-18 | -0.243085925 | 0.234 | 0.379 | 3.63E-14 |
| **WNT7B** | 4.32E-13 | -0.243182155 | 0.317 | 0.425 | 1.04E-08 |
| **TPM1** | 1.93E-24 | -0.243922158 | 0.2 | 0.373 | 4.65E-20 |
| **SRSF4** | 7.48E-16 | -0.244606188 | 0.684 | 0.757 | 1.81E-11 |
| **RPSA** | 5.6E-25 | -0.244649004 | 0.987 | 0.997 | 1.35E-20 |
| **CFH** | 8.56E-12 | -0.244786347 | 0.413 | 0.516 | 2.07E-07 |
| **RPL5** | 2.07E-33 | -0.246285118 | 0.992 | 0.997 | 4.99E-29 |
| **CCPG1** | 1.51E-14 | -0.246970524 | 0.72 | 0.788 | 3.65E-10 |
| **PALLD** | 1.28E-11 | -0.247386382 | 0.584 | 0.662 | 3.09E-07 |

| | | | | | |
|---|---|---|---|---|---|
| **ALDH3A2** | 8.43E-15 | -0.248093132 | 0.66 | 0.743 | 2.04E-10 |
| **FAT1** | 1.2E-10 | -0.249008277 | 0.45 | 0.532 | 2.89E-06 |
| **SET** | 4.42E-19 | -0.24934054 | 0.899 | 0.93 | 1.07E-14 |
| **EIF3H** | 7.12E-17 | -0.249374242 | 0.422 | 0.549 | 1.72E-12 |
| **IKZF2** | 4.46E-13 | -0.249713281 | 0.67 | 0.721 | 1.08E-08 |
| **CD164** | 9.09E-18 | -0.250163369 | 0.939 | 0.946 | 2.2E-13 |
| **CXADR** | 1.58E-16 | -0.251350319 | 0.496 | 0.617 | 3.83E-12 |
| **MUC4** | 1.46E-20 | -0.251932806 | 0.874 | 0.935 | 3.54E-16 |
| **RPL11** | 7.95E-38 | -0.251977331 | 0.996 | 1 | 1.92E-33 |
| **STT3B** | 2.19E-16 | -0.25301865 | 0.334 | 0.458 | 5.28E-12 |
| **MTSS1** | 1.84E-22 | -0.253221841 | 0.163 | 0.321 | 4.44E-18 |
| **LRRN1** | 9.27E-18 | -0.25360888 | 0.129 | 0.255 | 2.24E-13 |
| **RPL23A** | 3.98E-37 | -0.25379162 | 0.987 | 0.999 | 9.62E-33 |
| **ATRX** | 8.33E-19 | -0.2542616 | 0.829 | 0.887 | 2.01E-14 |
| **NCOA4** | 2.96E-16 | -0.255311242 | 0.399 | 0.525 | 7.15E-12 |
| **ITGA3** | 4.75E-15 | -0.255516187 | 0.439 | 0.541 | 1.15E-10 |
| **LYPD2** | 5.25E-09 | -0.255660965 | 0.552 | 0.699 | 0.00012688 |
| **MAT2A** | 6.03E-20 | -0.256481164 | 0.193 | 0.335 | 1.46E-15 |
| **EIF4B** | 3.08E-17 | -0.256518757 | 0.761 | 0.81 | 7.44E-13 |
| **SERINC1** | 5.84E-18 | -0.256754414 | 0.302 | 0.438 | 1.41E-13 |
| **LAPTM4B** | 6.89E-24 | -0.25694197 | 0.119 | 0.272 | 1.66E-19 |
| **RNF13** | 5.15E-15 | -0.257751886 | 0.43 | 0.534 | 1.24E-10 |
| **DENND11** | 1.43E-19 | -0.257905502 | 0.2 | 0.341 | 3.44E-15 |
| **SORL1** | 2.47E-21 | -0.258913459 | 0.126 | 0.27 | 5.96E-17 |
| **P4HB** | 5.61E-20 | -0.260890365 | 0.359 | 0.513 | 1.35E-15 |
| **KRT15** | 1.63E-17 | -0.261507104 | 0.134 | 0.265 | 3.93E-13 |
| **FOSB** | 1.19E-11 | -0.262226053 | 0.364 | 0.478 | 2.87E-07 |
| **PNN** | 2.32E-16 | -0.262736376 | 0.493 | 0.606 | 5.61E-12 |
| **BIRC6** | 1.56E-16 | -0.263034406 | 0.486 | 0.593 | 3.76E-12 |
| **USP22** | 6.04E-18 | -0.26325701 | 0.476 | 0.592 | 1.46E-13 |
| **RPS6** | 5.07E-43 | -0.26337353 | 0.997 | 1 | 1.23E-38 |
| **GLTP** | 8.06E-18 | -0.263639229 | 0.29 | 0.425 | 1.95E-13 |
| **CD59** | 4.55E-29 | -0.263822379 | 1 | 1 | 1.1E-24 |
| **B4GALT5** | 5.98E-29 | -0.26614531 | 0.316 | 0.527 | 1.44E-24 |
| **CAV2** | 2.67E-27 | -0.266234984 | 0.078 | 0.237 | 6.45E-23 |
| **TKT** | 2.14E-19 | -0.266257317 | 0.733 | 0.827 | 5.16E-15 |
| **GLUL** | 1.08E-20 | -0.266369881 | 0.89 | 0.955 | 2.62E-16 |
| **GARS-DT** | 6.48E-20 | -0.267457716 | 0.264 | 0.411 | 1.57E-15 |

| | | | | | |
|---|---|---|---|---|---|
| **KCNQ1OT1** | 2.65E-16 | -0.267802397 | 0.383 | 0.532 | 6.4E-12 |
| **SYNCRIP** | 2.21E-17 | -0.267952771 | 0.636 | 0.71 | 5.33E-13 |
| **GCLC** | 6.47E-16 | -0.268004671 | 0.809 | 0.87 | 1.56E-11 |
| **LAMB3** | 8.28E-25 | -0.268216178 | 0.038 | 0.171 | 2E-20 |
| **PHF3** | 7.32E-15 | -0.268709058 | 0.626 | 0.692 | 1.77E-10 |
| **ZBTB44** | 2.57E-17 | -0.268816758 | 0.558 | 0.664 | 6.21E-13 |
| **SLC22A16** | 8.43E-39 | -0.269026804 | 0.01 | 0.18 | 2.04E-34 |
| **KDM5B** | 1.31E-19 | -0.269146456 | 0.776 | 0.834 | 3.16E-15 |
| **LNPEP** | 3.43E-21 | -0.269605723 | 0.242 | 0.396 | 8.28E-17 |
| **CNN3** | 1.07E-19 | -0.269892225 | 0.428 | 0.585 | 2.58E-15 |
| **CAP1** | 2.68E-18 | -0.270311739 | 0.544 | 0.656 | 6.48E-14 |
| **ITGA2** | 5.42E-17 | -0.271188847 | 0.746 | 0.845 | 1.31E-12 |
| **RPS18** | 6.18E-42 | -0.271717044 | 0.996 | 1 | 1.49E-37 |
| **GK5** | 4.83E-17 | -0.272219601 | 0.437 | 0.549 | 1.17E-12 |
| **GSN** | 2.09E-24 | -0.272250039 | 0.996 | 0.997 | 5.06E-20 |
| **PPP3CA** | 4.84E-20 | -0.272412917 | 0.228 | 0.375 | 1.17E-15 |
| **RPS16** | 5.33E-44 | -0.272719258 | 0.994 | 0.999 | 1.29E-39 |
| **AFF4** | 2.55E-17 | -0.272733247 | 0.461 | 0.572 | 6.15E-13 |
| **RPLP1** | 2.53E-41 | -0.272786313 | 0.999 | 1 | 6.11E-37 |
| **NF1** | 1.61E-18 | -0.273404028 | 0.337 | 0.468 | 3.9E-14 |
| **CDK6** | 7.16E-21 | -0.273593663 | 0.139 | 0.28 | 1.73E-16 |
| **PCDH7** | 7.06E-14 | -0.273641806 | 0.426 | 0.535 | 1.71E-09 |
| **INSIG1** | 9.85E-15 | -0.274184115 | 0.299 | 0.426 | 2.38E-10 |
| **PCSK7** | 6.06E-23 | -0.274607662 | 0.246 | 0.408 | 1.46E-18 |
| **EEF1G** | 1.62E-35 | -0.275606576 | 0.99 | 0.995 | 3.91E-31 |
| **ARID1B** | 6.3E-19 | -0.275781377 | 0.71 | 0.778 | 1.52E-14 |
| **H3F3B** | 1.1E-27 | -0.276772321 | 0.998 | 0.999 | 2.66E-23 |
| **ASAH1** | 7E-24 | -0.276870534 | 0.51 | 0.682 | 1.69E-19 |
| **LEPROT** | 2.22E-19 | -0.276884176 | 0.443 | 0.581 | 5.35E-15 |
| **PTP4A2** | 8.95E-25 | -0.277625366 | 0.653 | 0.781 | 2.16E-20 |
| **RPLP2** | 7.59E-42 | -0.278108459 | 0.992 | 0.999 | 1.83E-37 |
| **ASPH** | 1.78E-20 | -0.278576857 | 0.399 | 0.552 | 4.3E-16 |
| **EGFR** | 4.63E-19 | -0.279639412 | 0.284 | 0.425 | 1.12E-14 |
| **SLC9A3R1** | 1.12E-22 | -0.280306348 | 0.768 | 0.862 | 2.7E-18 |
| **MUC20** | 4.85E-22 | -0.280429785 | 0.851 | 0.917 | 1.17E-17 |
| **LINC00511** | 1.71E-16 | -0.282605766 | 0.451 | 0.56 | 4.13E-12 |
| **JPX** | 2.49E-19 | -0.283827436 | 0.75 | 0.797 | 6E-15 |
| **PTPN3** | 1.34E-20 | -0.284626065 | 0.387 | 0.528 | 3.24E-16 |

| | | | | | |
|---|---|---|---|---|---|
| **PPIA** | 2.36E-22 | -0.284636696 | 0.873 | 0.932 | 5.7E-18 |
| **TUG1** | 1.46E-20 | -0.284779776 | 0.419 | 0.549 | 3.52E-16 |
| **EPPK1** | 6.26E-13 | -0.285151963 | 0.706 | 0.765 | 1.51E-08 |
| **RDX** | 7.89E-22 | -0.285907066 | 0.603 | 0.71 | 1.91E-17 |
| **RDH10** | 2.39E-29 | -0.286885738 | 0.183 | 0.379 | 5.77E-25 |
| **RPL3** | 7.41E-54 | -0.289495969 | 0.99 | 0.999 | 1.79E-49 |
| **CFL1** | 1.03E-26 | -0.289547983 | 0.886 | 0.942 | 2.49E-22 |
| **RPL14** | 1.29E-38 | -0.289586935 | 0.979 | 0.995 | 3.13E-34 |
| **RPS15A** | 3.97E-45 | -0.289690773 | 0.99 | 0.999 | 9.59E-41 |
| **LPP** | 2.13E-20 | -0.289695701 | 0.552 | 0.684 | 5.15E-16 |
| **CLTC** | 3.15E-27 | -0.289780697 | 0.921 | 0.948 | 7.61E-23 |
| **KLF3** | 1.99E-22 | -0.290253929 | 0.45 | 0.6 | 4.81E-18 |
| **DHRS3** | 1.3E-18 | -0.290280641 | 0.54 | 0.658 | 3.15E-14 |
| **DYNC1H1** | 1.22E-26 | -0.290413821 | 0.853 | 0.912 | 2.95E-22 |
| **ADGRG1** | 1.78E-21 | -0.291568644 | 0.313 | 0.466 | 4.29E-17 |
| **ANKRD50** | 1.94E-29 | -0.291705408 | 0.099 | 0.269 | 4.69E-25 |
| **NKTR** | 5.74E-20 | -0.292218934 | 0.431 | 0.567 | 1.39E-15 |
| **PSAP** | 1.39E-29 | -0.292656557 | 0.883 | 0.947 | 3.36E-25 |
| **B4GALT1** | 7.33E-21 | -0.293277761 | 0.441 | 0.583 | 1.77E-16 |
| **VPS13C** | 1.33E-23 | -0.293617587 | 0.937 | 0.95 | 3.22E-19 |
| **TM9SF3** | 1.22E-22 | -0.29391335 | 0.669 | 0.771 | 2.95E-18 |
| **RPL27A** | 5.18E-39 | -0.295919509 | 0.989 | 0.999 | 1.25E-34 |
| **NACA** | 1.54E-41 | -0.296679282 | 0.984 | 0.997 | 3.71E-37 |
| **AC005261.1** | 4.39E-24 | -0.29782878 | 0.292 | 0.463 | 1.06E-19 |
| **PDIA3** | 9.51E-21 | -0.298866754 | 0.663 | 0.76 | 2.3E-16 |
| **PPP1R14B** | 3.04E-29 | -0.300278696 | 0.262 | 0.467 | 7.33E-25 |
| **DSG2** | 1.22E-19 | -0.300627083 | 0.454 | 0.584 | 2.94E-15 |
| **PMEPA1** | 1.91E-24 | -0.300932548 | 0.026 | 0.151 | 4.62E-20 |
| **TBL1XR1** | 1.86E-22 | -0.302508697 | 0.683 | 0.765 | 4.5E-18 |
| **SKIL** | 1.41E-15 | -0.303029742 | 0.506 | 0.61 | 3.4E-11 |
| **RPS10** | 8.53E-39 | -0.306404892 | 0.983 | 0.996 | 2.06E-34 |
| **IFI16** | 3.57E-23 | -0.307156365 | 0.521 | 0.673 | 8.63E-19 |
| **RPS14** | 2.53E-63 | -0.308703075 | 0.992 | 1 | 6.11E-59 |
| **IFNGR1** | 2.45E-23 | -0.309193517 | 0.28 | 0.438 | 5.91E-19 |
| **RPL13A** | 3.86E-54 | -0.309260839 | 0.996 | 0.999 | 9.33E-50 |
| **CHD3** | 4.05E-23 | -0.309658602 | 0.434 | 0.578 | 9.78E-19 |
| **AGL** | 1.27E-20 | -0.309722164 | 0.484 | 0.608 | 3.08E-16 |
| **HP1BP3** | 1E-19 | -0.309761576 | 0.4 | 0.529 | 2.42E-15 |

| | | | | | |
|---|---|---|---|---|---|
| **NUPR1** | 1.88E-24 | -0.309976777 | 0.697 | 0.874 | 4.54E-20 |
| **RPL12** | 9.71E-44 | -0.310232229 | 0.988 | 0.999 | 2.34E-39 |
| **LITAF** | 2.43E-24 | -0.311925055 | 0.721 | 0.834 | 5.87E-20 |
| **RBP1** | 8.62E-24 | -0.314008269 | 0.128 | 0.288 | 2.08E-19 |
| **DDIT4** | 1.76E-26 | -0.316045036 | 0.187 | 0.374 | 4.25E-22 |
| **MT1X** | 2.02E-61 | -0.316152324 | 0.068 | 0.364 | 4.89E-57 |
| **HIF1A** | 4.22E-31 | -0.316267755 | 0.156 | 0.348 | 1.02E-26 |
| **MIR205HG** | 2.26E-29 | -0.317220759 | 0.764 | 0.931 | 5.47E-25 |
| **NFIA** | 6.4E-25 | -0.317990026 | 0.888 | 0.912 | 1.55E-20 |
| **CSDE1** | 5.39E-33 | -0.3190219 | 0.919 | 0.954 | 1.3E-28 |
| **CCNI** | 6.22E-33 | -0.319089263 | 0.942 | 0.965 | 1.5E-28 |
| **ADGRV1** | 3.71E-23 | -0.31939034 | 0.197 | 0.356 | 8.97E-19 |
| **CFD** | 3.4E-11 | -0.319420493 | 0.221 | 0.319 | 8.22E-07 |
| **F3** | 3.07E-35 | -0.319762654 | 0.181 | 0.414 | 7.42E-31 |
| **RPL4** | 2.47E-41 | -0.320070107 | 0.973 | 0.989 | 5.96E-37 |
| **MEIS2** | 1.96E-22 | -0.320957561 | 0.596 | 0.692 | 4.74E-18 |
| **ITM2B** | 8.1E-35 | -0.321613174 | 0.988 | 0.997 | 1.96E-30 |
| **CPD** | 1.13E-25 | -0.321819318 | 0.516 | 0.671 | 2.74E-21 |
| **NCKAP1** | 6.19E-24 | -0.322552417 | 0.497 | 0.623 | 1.5E-19 |
| **DDX3X** | 7.27E-28 | -0.322873833 | 0.8 | 0.88 | 1.75E-23 |
| **RPL32** | 5.9E-54 | -0.324493321 | 0.991 | 0.998 | 1.42E-49 |
| **RPS27L** | 7.91E-28 | -0.325214131 | 0.723 | 0.838 | 1.91E-23 |
| **EIF2S3** | 6.44E-24 | -0.326929638 | 0.4 | 0.537 | 1.56E-19 |
| **RPS19** | 1.17E-54 | -0.32787296 | 0.993 | 1 | 2.83E-50 |
| **KRT5** | 1.95E-33 | -0.328145953 | 0.01 | 0.161 | 4.72E-29 |
| **MUC16** | 1.68E-27 | -0.33102197 | 0.97 | 0.982 | 4.07E-23 |
| **CYP24A1** | 3.05E-09 | -0.33113598 | 0.142 | 0.226 | 7.37E-05 |
| **NFIB** | 7.89E-23 | -0.333783637 | 0.388 | 0.54 | 1.91E-18 |
| **SEC11A** | 1.01E-27 | -0.334567482 | 0.439 | 0.599 | 2.44E-23 |
| **AHR** | 2.17E-26 | -0.335076109 | 0.433 | 0.597 | 5.25E-22 |
| **FN1** | 1.95E-21 | -0.335260894 | 0.018 | 0.124 | 4.71E-17 |
| **LIMA1** | 8.48E-28 | -0.336206152 | 0.334 | 0.525 | 2.05E-23 |
| **CYP2S1** | 5.31E-28 | -0.337289197 | 0.361 | 0.546 | 1.28E-23 |
| **BZW1** | 4.41E-32 | -0.339676825 | 0.719 | 0.837 | 1.06E-27 |
| **PABPC1** | 1.41E-34 | -0.339778043 | 0.902 | 0.958 | 3.4E-30 |
| **PNISR** | 4.46E-35 | -0.339880898 | 0.77 | 0.869 | 1.08E-30 |
| **EEF1D** | 1.17E-37 | -0.340831369 | 0.778 | 0.897 | 2.83E-33 |
| **TUBB** | 3.03E-40 | -0.341902844 | 0.116 | 0.336 | 7.32E-36 |

| | | | | | |
|---|---|---|---|---|---|
| **ZNF207** | 1.07E-34 | -0.342246974 | 0.75 | 0.828 | 2.58E-30 |
| **ALCAM** | 3.87E-48 | -0.344350612 | 0.989 | 0.998 | 9.34E-44 |
| **KLF5** | 3.7E-36 | -0.344388945 | 0.836 | 0.927 | 8.95E-32 |
| **GPX2** | 5.91E-18 | -0.345258598 | 0.523 | 0.631 | 1.43E-13 |
| **MDM4** | 9.56E-32 | -0.347993843 | 0.36 | 0.55 | 2.31E-27 |
| **ASH1L** | 4.78E-27 | -0.348121202 | 0.55 | 0.692 | 1.15E-22 |
| **RPS9** | 1.57E-63 | -0.348889427 | 0.993 | 0.999 | 3.8E-59 |
| **GAS5** | 3.07E-37 | -0.350762595 | 0.138 | 0.348 | 7.42E-33 |
| **ARPC5** | 2.38E-36 | -0.351017726 | 0.372 | 0.593 | 5.75E-32 |
| **SOX4** | 1.58E-39 | -0.352126459 | 0.626 | 0.859 | 3.82E-35 |
| **UBA52** | 3.58E-57 | -0.354340883 | 0.989 | 0.998 | 8.64E-53 |
| **ARGLU1** | 1.18E-32 | -0.355064308 | 0.68 | 0.817 | 2.85E-28 |
| **EIF3E** | 3.62E-34 | -0.355664013 | 0.724 | 0.819 | 8.73E-30 |
| **SNHG14** | 9.82E-27 | -0.356726026 | 0.403 | 0.574 | 2.37E-22 |
| **RPL26** | 1.21E-67 | -0.35707266 | 0.99 | 0.999 | 2.92E-63 |
| **PTMS** | 3.03E-38 | -0.357821764 | 0.15 | 0.372 | 7.32E-34 |
| **RPS15** | 2.55E-76 | -0.358790968 | 0.994 | 1 | 6.16E-72 |
| **LAMP2** | 1.84E-28 | -0.359242297 | 0.414 | 0.569 | 4.43E-24 |
| **TXNIP** | 5.85E-42 | -0.359424036 | 0.847 | 0.968 | 1.41E-37 |
| **CTSD** | 5.51E-36 | -0.359693646 | 0.939 | 0.984 | 1.33E-31 |
| **FNDC3B** | 3.17E-28 | -0.361087321 | 0.302 | 0.48 | 7.66E-24 |
| **RPL15** | 9.06E-85 | -0.363995491 | 0.996 | 1 | 2.19E-80 |
| **WSB1** | 2.47E-34 | -0.364782875 | 0.587 | 0.746 | 5.95E-30 |
| **LRATD1** | 2.22E-21 | -0.365990634 | 0.511 | 0.639 | 5.37E-17 |
| **MYH9** | 1.5E-45 | -0.36697634 | 0.241 | 0.495 | 3.62E-41 |
| **ADAM9** | 8.74E-37 | -0.368275471 | 0.251 | 0.475 | 2.11E-32 |
| **RPL10A** | 5.77E-64 | -0.37125892 | 0.981 | 0.999 | 1.39E-59 |
| **RACK1** | 1.68E-54 | -0.373334954 | 0.972 | 0.993 | 4.05E-50 |
| **SDC1** | 8.18E-43 | -0.377052544 | 0.128 | 0.362 | 1.98E-38 |
| **FAM3C** | 1.95E-32 | -0.37717783 | 0.556 | 0.721 | 4.7E-28 |
| **RPL9** | 5.4E-89 | -0.380826001 | 0.996 | 1 | 1.31E-84 |
| **CTNNB1** | 8.26E-28 | -0.381165136 | 0.713 | 0.805 | 2E-23 |
| **GBP3** | 1.23E-36 | -0.381575463 | 0.491 | 0.683 | 2.96E-32 |
| **BAG1** | 5.64E-45 | -0.382489868 | 0.828 | 0.931 | 1.36E-40 |
| **BTG1** | 4.54E-38 | -0.384665087 | 0.397 | 0.626 | 1.1E-33 |
| **HSPA5** | 1.09E-32 | -0.386640862 | 0.788 | 0.896 | 2.63E-28 |
| **SLC38A2** | 1.4E-41 | -0.388361175 | 0.12 | 0.344 | 3.38E-37 |
| **TPM4** | 9.68E-58 | -0.390043838 | 0.147 | 0.431 | 2.34E-53 |

| | | | | | |
|---|---|---|---|---|---|
| **RPL13** | 1.54E-95 | -0.390353235 | 0.997 | 1 | 3.73E-91 |
| **RPL18** | 3.37E-81 | -0.390534493 | 0.987 | 1 | 8.14E-77 |
| **SERINC2** | 2.94E-38 | -0.394305208 | 0.352 | 0.575 | 7.1E-34 |
| **CTSB** | 8.93E-38 | -0.39684835 | 0.646 | 0.85 | 2.16E-33 |
| **CD24** | 1.42E-62 | -0.397078018 | 0.999 | 1 | 3.44E-58 |
| **LDHA** | 1.64E-53 | -0.397636044 | 0.047 | 0.288 | 3.95E-49 |
| **RPL23** | 1.04E-66 | -0.397748043 | 0.989 | 0.999 | 2.52E-62 |
| **TMSB4X** | 1.43E-67 | -0.398548393 | 0.789 | 0.98 | 3.47E-63 |
| **GRN** | 4.54E-43 | -0.400110738 | 0.124 | 0.362 | 1.1E-38 |
| **RPS11** | 1.33E-92 | -0.400218358 | 0.993 | 1 | 3.21E-88 |
| **CD81** | 2.74E-39 | -0.40301064 | 0.684 | 0.818 | 6.61E-35 |
| **LMO4** | 1.97E-44 | -0.40416137 | 0.324 | 0.581 | 4.76E-40 |
| **FTX** | 2.73E-39 | -0.406584319 | 0.364 | 0.587 | 6.59E-35 |
| **NFAT5** | 1.34E-38 | -0.406717363 | 0.35 | 0.559 | 3.23E-34 |
| **AHNAK** | 9.67E-47 | -0.412705249 | 0.764 | 0.915 | 2.34E-42 |
| **RPL29** | 1.54E-97 | -0.415260386 | 0.991 | 0.999 | 3.72E-93 |
| **CALR** | 2.9E-38 | -0.415618926 | 0.667 | 0.82 | 7.01E-34 |
| **RPS3A** | 1.48E-104 | -0.415870612 | 0.993 | 1 | 3.59E-100 |
| **IGFBP3** | 1.59E-44 | -0.423306802 | 0.139 | 0.395 | 3.83E-40 |
| **ITGAV** | 2.91E-47 | -0.427860483 | 0.186 | 0.431 | 7.03E-43 |
| **ATP1B3** | 2.28E-52 | -0.428973047 | 0.068 | 0.316 | 5.5E-48 |
| **TRAM1** | 9.31E-44 | -0.430702896 | 0.441 | 0.667 | 2.25E-39 |
| **BHLHE40** | 3.16E-37 | -0.431256746 | 0.178 | 0.393 | 7.63E-33 |
| **RUNX1** | 5.27E-45 | -0.43166884 | 0.719 | 0.839 | 1.27E-40 |
| **ETS2** | 9.18E-35 | -0.433279537 | 0.518 | 0.676 | 2.22E-30 |
| **RPL8** | 1.08E-100 | -0.441142777 | 0.997 | 1 | 2.61E-96 |
| **N4BP2L2** | 6.03E-54 | -0.442753427 | 0.741 | 0.883 | 1.46E-49 |
| **LINC00342** | 1.09E-32 | -0.448533474 | 0.394 | 0.615 | 2.62E-28 |
| **RPL10** | 9.33E-100 | -0.45789859 | 0.987 | 0.999 | 2.25E-95 |
| **PKM** | 5.47E-53 | -0.458498922 | 0.861 | 0.95 | 1.32E-48 |
| **RPL18A** | 2.23E-95 | -0.462940407 | 0.99 | 1 | 5.38E-91 |
| **ABCA13** | 4.7E-31 | -0.463031771 | 0.948 | 0.976 | 1.14E-26 |
| **EEF1A1** | 1.19E-144 | -0.463977811 | 0.999 | 1 | 2.88E-140 |
| **EEF2** | 2.69E-61 | -0.464299004 | 0.831 | 0.922 | 6.49E-57 |
| **SLC25A6** | 2.83E-54 | -0.469333786 | 0.396 | 0.654 | 6.83E-50 |
| **SQSTM1** | 7.4E-61 | -0.473254086 | 0.761 | 0.916 | 1.79E-56 |
| **TMPRSS4** | 1.2E-44 | -0.47377436 | 0.383 | 0.616 | 2.91E-40 |
| **APLP2** | 3.8E-56 | -0.474313022 | 0.878 | 0.946 | 9.18E-52 |

| | | | | | |
|---|---|---|---|---|---|
| **PERP** | 5.61E-103 | -0.477222703 | 0.998 | 1 | 1.36E-98 |
| **LAMP1** | 4.09E-58 | -0.478539337 | 0.371 | 0.637 | 9.88E-54 |
| **HNRNPH1** | 3.77E-55 | -0.481821931 | 0.333 | 0.594 | 9.12E-51 |
| **SNHG5** | 4.25E-42 | -0.486927137 | 0.353 | 0.561 | 1.03E-37 |
| **RPL28** | 1.66E-117 | -0.487829111 | 0.993 | 1 | 4E-113 |
| **ACTG1** | 4.21E-56 | -0.497593888 | 0.862 | 0.979 | 1.02E-51 |
| **RPS4X** | 1.36E-112 | -0.503742533 | 0.99 | 0.999 | 3.27E-108 |
| **RPLP0** | 3.1E-97 | -0.518650477 | 0.992 | 0.999 | 7.48E-93 |
| **NORAD** | 3.54E-59 | -0.519897249 | 0.623 | 0.802 | 8.55E-55 |
| **DGKH** | 3.55E-52 | -0.525601334 | 0.842 | 0.917 | 8.57E-48 |
| **VMP1** | 5.09E-58 | -0.528587129 | 0.756 | 0.934 | 1.23E-53 |
| **DDX17** | 4.39E-95 | -0.542759796 | 0.918 | 0.98 | 1.06E-90 |
| **RPS3** | 3.1E-117 | -0.544923263 | 0.966 | 0.997 | 7.49E-113 |
| **SPON2** | 4.85E-57 | -0.549771094 | 0.38 | 0.659 | 1.17E-52 |
| **APP** | 5.17E-58 | -0.555633504 | 0.543 | 0.754 | 1.25E-53 |
| **SYT8** | 9.6E-46 | -0.567481604 | 0.333 | 0.574 | 2.32E-41 |
| **RPL7** | 1.07E-133 | -0.57876996 | 0.98 | 0.997 | 2.57E-129 |
| **AHNAK2** | 2.19E-61 | -0.600044052 | 0.706 | 0.897 | 5.29E-57 |
| **ITGB1** | 1.32E-75 | -0.602137995 | 0.509 | 0.772 | 3.18E-71 |
| **RPS5** | 1.17E-112 | -0.613627607 | 0.929 | 0.987 | 2.83E-108 |
| **GABPB1-AS1** | 2.43E-71 | -0.63052293 | 0.707 | 0.86 | 5.88E-67 |
| **SERPINB3** | 2.28E-64 | -0.645714413 | 0.069 | 0.374 | 5.5E-60 |
| **SNHG29** | 9.11E-105 | -0.647274341 | 0.379 | 0.756 | 2.2E-100 |
| **AKR1C2** | 4.54E-59 | -0.648075373 | 0.19 | 0.488 | 1.1E-54 |
| **XIST** | 1.06E-39 | -0.6549785 | 0 | 0.167 | 2.56E-35 |
| **ZFP36L1** | 5.41E-83 | -0.659554367 | 0.592 | 0.868 | 1.31E-78 |
| **POLR2J3** | 7.33E-95 | -0.663365477 | 0.29 | 0.642 | 1.77E-90 |
| **CCDC39** | 1.26E-128 | -0.677281147 | 0.742 | 0.937 | 3.04E-124 |
| **ABLIM1** | 9.65E-74 | -0.680700722 | 0.532 | 0.785 | 2.33E-69 |
| **AKR1C1** | 3.9E-69 | -0.706386181 | 0.117 | 0.423 | 9.42E-65 |
| **EPAS1** | 1.01E-101 | -0.73576594 | 0.181 | 0.588 | 2.43E-97 |
| **S100A2** | 4.02E-106 | -0.762559214 | 0.09 | 0.5 | 9.7E-102 |
| **PRSS23** | 6.62E-106 | -0.804447337 | 0.343 | 0.727 | 1.6E-101 |
| **GPNMB** | 1.76E-67 | -0.856354187 | 0.016 | 0.288 | 4.25E-63 |
| **HSPB1** | 6.55E-96 | -0.86281939 | 0.401 | 0.772 | 1.58E-91 |
| **PSCA** | 2.33E-81 | -1.058089281 | 0.961 | 0.985 | 5.62E-77 |
| **NEAT1** | 4.03E-255 | -1.067485953 | 0.969 | 1 | 9.72E-251 |
| **FMO3** | 1.79E-99 | -1.153339541 | 0.044 | 0.42 | 4.32E-95 |
